# Insular holobionts: persistence and seasonal plasticity of the Balearic wall lizard (*Podarcis lilfordi*) gut microbiota

**DOI:** 10.1101/2022.05.19.492253

**Authors:** Laura Baldo, Giacomo Tavecchia, Andreu Rotger, José Manuel Igual, Joan Lluís Riera

**Affiliations:** Department of Evolutionary Biology, Ecology and Environmental Sciences, University of Barcelona, Spain; Institute for Research on Biodiversity (IRBio), University of Barcelona, Barcelona, Spain; Animal Demography and Ecology Unit, IMEDEA, CSIC-UIB, Esporles, Spain

**Keywords:** within and among population diversity, 16S rRNA Illumina, sex, season, stable isotopes, fecal diet, plasticity, adaptation, island

## Abstract

Integrative studies of animals and associated microbial assemblages (i.e., the holobiont) are rapidly changing our perspectives on organismal ecology and evolution. Islands provide ideal natural systems to understand the biogeographic patterns that shape these symbiotic associations, their resilience and plasticity over temporal and spatial scales, and ultimately their role in the host ecological adaptation. Here we used the Balearic wall lizard *Podarcis lilfordi* to address the diversification of the holobiont in an insular context by dissecting the drivers of the gut microbiota diversity within and across host allopatric populations. By extensive fecal sampling of individually identified lizards from three closed populations/islets in the South of Mallorca (Na Moltona, Na Guardis and En Curt) along two years and two seasons (spring and autumn), we sorted out the effect of islet, year, season, sex and partly life stage on the microbiota composition. We further related microbiota distances to host genetics and trophic ecology. Overall, the three populations showed a remarkable conservation of the major microbial taxonomic profile, while carrying their unique microbial signature at finer level of taxonomic resolution (Amplicon Sequence Variants (ASVs)). Microbiota distances across populations were compatible with both host genetics (as inferred by microsatellites) and trophic niche distances (as inferred by stable isotopes and fecal content). Within populations, a large proportion of ASVs (30-50%) persisted along the four sampling dates. Microbial diversity was driven by life stage and season, with no annual or sex effect. Seasonal changes within islets were mainly associated with fluctuations in the relative abundances of few bacterial taxa (mostly families Lachnospiraceae and Ruminococcaceae), consistently in both sampled years and without any major compositional turnover. These results support a large resilience of the major compositional aspects of the *P. lilfordi* gut microbiota over the short-term evolutionary divergence of their host allopatric populations (<10,000 years), but also suggest an undergoing process of parallel diversification of the holobiont. The cyclic seasonal fluctuations in gut microbiota composition hint to an important plasticity of these bacterial communities in response to the host annual physiological/metabolic shifts. The importance of these microbial community dynamics in the host ecology and dietary flexibility remains to be investigated.

## Introduction

All organisms live in symbiosis with complex microbial communities (i.e., the host microbiota), which are known to exert fundamental biological functions for their host, effectively providing an extended phenotype for increased ecological adaptation (Alberdi et al., 2016; Henry et al., 2021). The host microbiota, and particularly those communities inhabiting the intestinal tract, known as gut microbiota, are known to affect a multitude of biological functions (Levin et al., 2021), including the host immune response (Thaiss et al., 2016), development (Warne et al., 2019), behavior (Rowe et al., 2020), thermal regulation (Huus & Ley, 2021), dietary preferences (Kohl et al., 2014; Leitão-Gonç alves et al., 2017), trophic niche amplitude (Kohl et al., 2016), digestion rates, and the overall efficiency in resource use (Lindsay et al., 2020). Collectively this indicates a critical role of the gut microbiota in forging the host ecology and influencing its evolutionary outcomes (Alberdi et al., 2016; Shapira, 2016). On the other hand, the gut microbiota structure itself is largely shaped by the host factors, including genotype, diet (Rojas et al., 2021; Youngblut et al., 2019), and geography (Levin et al., 2021), with results varying between captive and wild samples (Youngblut et al., 2019) and depending on the taxonomic scale of observation, both for microbes (from strain to phylum) and hosts (from individuals up to species and families) (Alberdi et al., 2021; Rojas et al., 2021).

Islands represent simplified models to study organismal adaptation due to their isolated nature, small population sizes (with no immigration or emigration events), reduced selective pressures, such as predation or food competitors, and smaller ecological networks (MacArthur & Wilson, 1967), overall providing natural laboratories for dissecting the deterministic and stochastic mechanisms responsible of organismal diversification and fine-tune adaptation to their environments (Bittkau & Comes, 2005; Velo-Antó et al., 2012). These same critical aspects make islands a neat system also for integrative studies of both host and microbial associates (i.e., the holobiont) aimed to understand the coevolution of this symbiosis, the factors that shape it, and its potential contribution to the host insular adaptation (Baldo et al., 2018; Davison et al., 2018; Lankau et al., 2012; Michel et al., 2018).

The Balearic wall lizard *Podarcis lilfordi*, also known as the Lilford’s wall lizard, represents a particularly suitable system for this purpose (Baldo et al., 2018). The species is endemic to the Balearic Islands and currently comprises several island populations in the archipelagos of Cabrera, Mallorca and Menorca (Salvador, 2009). During the last ice age, the ancestral populations present in the mainland of Mallorca and Menorca dispersed to offshore islets following a vicariance process; after the Post-Messinian isolation that started about 2.6 million years ago (Brown et al., 2008; Terrasa et al., 2009), populations remained confined, while Mallorca and Menorca ancestral populations were driven to extinction by the introduction of predators (Alcover, 2000).

All sister populations from these small continental islands are bonded by their historical legacy (common ancestry and a similar genetic background) (Brown et al., 2008; Buades et al., 2013; Pérez-Cembranos et al., 2020; Terrasa et al., 2009), while representing evolutionary units (Pérez-Cembranos et al., 2020) given their discrete geographic boundaries that have limited both host and gut microbe dispersion since their divergence. In most of these islets, the community of terrestrial vertebrates is dominated by lizards (only few islets show presence of geckos, mice, rats and rabbits), reducing chances of interspecific interactions. Furthermore, their shared climate, reduced area and low biotic diversity make these populations largely controllable and comparable systems, facilitating the study of the major common factors forging and maintaining this symbiotic association (Baldo et al., 2018; Lankau et al., 2012; Michel et al., 2018). In a previous study on seven Menorcan populations we have characterised for the first time the composition of the Lilford’s wall lizard gut microbiota, revealing a large conservation of the taxonomic profiles (resembling the typical vertebrate microbiota), with a potential impact of the phylogeographic history and ecological drift in shaping microbial diversity (Baldo et al., 2018). However, the sampling design and size (five individuals per population) did not allow us to explore drivers of within population microbiota diversity, particularly sex and temporal dynamics, and the putative functional role of these communities in lizard adaptation.

In the present study we focused on three well-studied populations of Lilford’s wall lizard found in three close islets south of Mallorca: Na Moltona, Na Guardis and En Curt (Figure 1). The three islets are less than five km apart, share nearly the same Mediterranean climate and biotic environment, and differ primarily in size and biotic index (Rotger et al., 2020). In particular, the smallest islet (En Curt) hosts a reduced plant community compared to the other two islets (Rotger et al., 2021). The lizard represents the major vertebrate species on each islet, with density ranging from 350 to 2500 ind/ha (Rotger et al., 2016). Since 2010, the three populations have been under a demographic study: every spring and autumn, individuals are sampled through a capture and recapture method, photo-identified by digitally recording the pattern of ventral scales (Moya et al., 2015), sexed and measured (i.e., body length and weight). This has provided important individual-level and longitudinal data for each population, showing that the three populations differ in demographic parameters and life history traits such as body growth rate, fecundity, survival, and density(Rotger et al., 2016, 2020). A recent genetic study based on microsatellites indicates that the three populations diverged about 5,000 - 8,000 years ago (with no recent relevant immigration events), and that Na Moltona and Na Guardis represent closer sister populations (Rotger et al., 2021). Diet composition of the latter islets has been recently characterized based on fecal content of several individuals, showing that these two populations share a similar trophic ecology, primarily based on arthropods (with plant integration), with sex differences and seasonal variations in response to resource availability (Santamaría et al., 2019).

**Figure 1:**
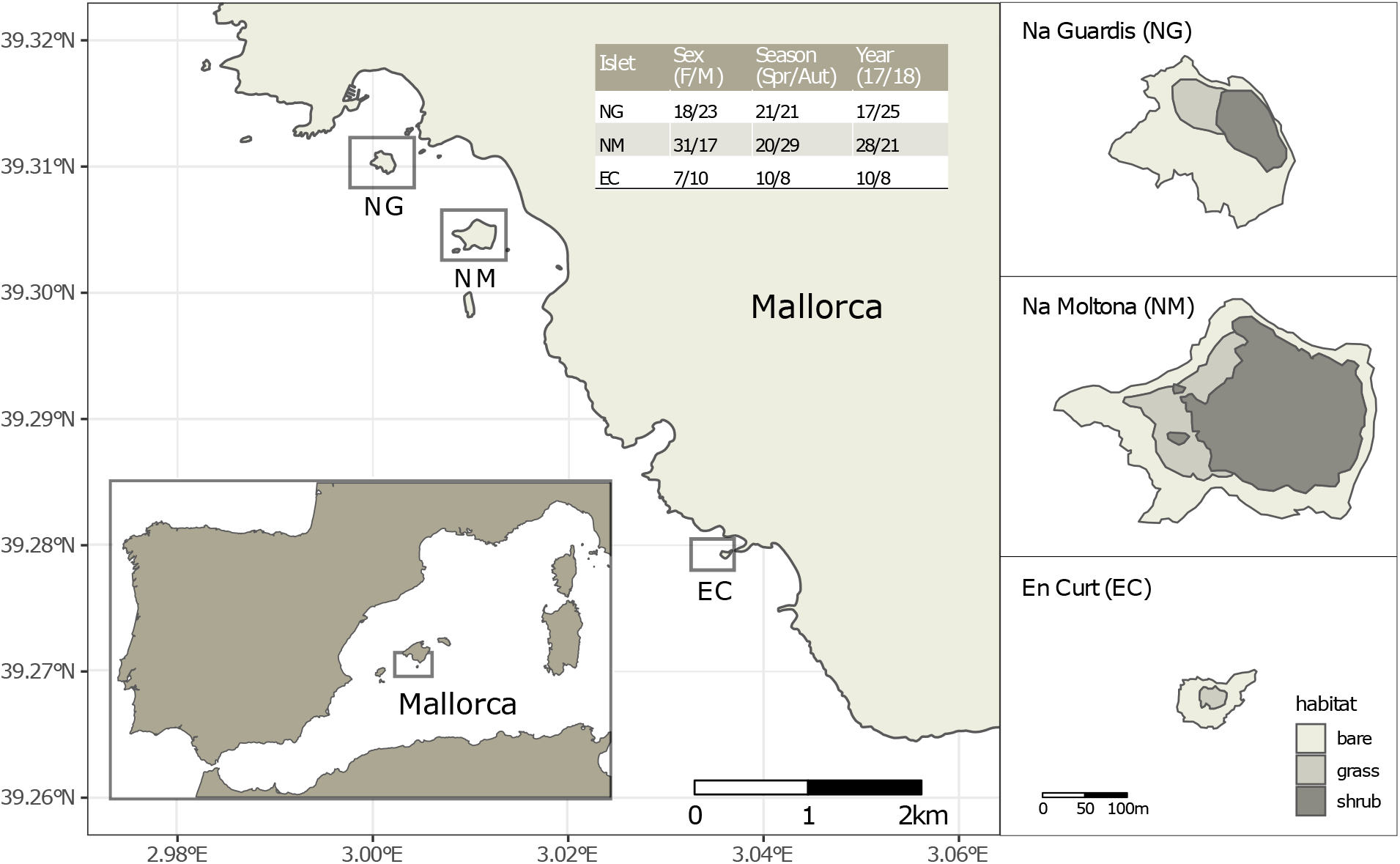
Location of the three islets under study and sample statistics. The table lists the number of fecal samples per sex (females and males, without including one unsexed juvenile per islet), season (spring and autumn) and year (2017 and 2018). See Appendix S1, Table S1, for sample metadata.

This extensive background population and individual-level information provides a powerful setting to begin addressing patterns of the Lilford’s wall lizard gut microbiota variation within and among populations, within their short temporal scale of divergence (< 10,000 years). This variation can also inform on the potential role of the gut microbiota in the adaptation of the lizards to their insular environment.

Here we specifically surveyed the impact of the islet phylogeographic distance (as inferred by microsatellites), host trophic niche (according to both fecal content and stable isotopes) and intrinsic factors (sex and life stage) in shaping the gut microbial communties. We also explored, for the first time in this species, the level of persistence and plasticity of these microbial communities over time, by repeated sampling over two years and two seasons. The aim was to understand the strength of this association and the degree to which these communities change in composition along seasons. Recent studies in vertebrates have shown that the microbiota can be highly plastic (Alberdi et al., 2016; Gomez et al., 2019), and changes its composition in response to the host’s physiological changes for optimization of the energy metabolism (Huus & Ley, 2021; Sommer et al., 2016) effectively boosting the host ecological adaptation by increasing its phenotypic plasticity (Guo et al., 2021; Hicks et al., 2018; Smits et al., 2017). As lizard populations experience a strong limitation in resources, particularly critical during the dry autumn season (Santamaría et al., 2019), we set to investigate whether microbiota showed evidence of seasonal plasticity that might hint to a functional role of the microbiota in buffering the lizard’s seasonal metabolic needs.

To address these goals, we sampled fecal matter from a subset of individually identified lizards of each of the three populations (both males and females) for two years, during spring and autumn seasons, and characterized the fecal microbiota by amplicon 16S rRNA sequencing.

The following major questions were targeted: what is the relative contribution of the host phylogeographic history and trophic ecological adaptation in driving gut microbiota divergence among allopatric populations? What are the major factors that structure the microbiota within these closed populations? What is the level of persistence of microbial taxa within a population over time? Does the gut microbiota show evidence of seasonal plasticity? Finally, are the observed microbial dynamics similar across islets/populations, suggesting a deterministic pattern?

## Materials and Methods

### Microbiota sampling and metadata collection

Faeces were collected from a total of 109 individually identified specimens of *P. lilfordi* in three islets off the south shore of Mallorca Island: Na Guardis (NG) (42 samples), Na Moltona (NM) (49) and En Curt (EC) (18) (Figure 1, Appendix S1, Table S1 for sample metadata). Sampling was performed during spring (April) and autumn (October) in 2017 and 2018, for a total of four sampling dates (“Spring-17”, “Autumn-17”, “Spring-18” and “Autumn-18”, each referred simply as to “Date”). The reproduction period is extended from Spring to the end of the Summer. Animals begin to lay eggs in May and females can lay two to three clutches (Castilla & Bauwens, 2000), usually of two to four eggs (Rotger et al., 2020). All specimens were caught in georeferenced pitfall traps containing sterile fruit juices placed along paths and vegetation edges. Specimens were weighed, and body size measured as snout to vent length (SVL). Life stage (adults, subadults and juveniles) was assigned based on the SVL, according to the mean values for the smallest described subspecies (Salvador, 2009). Age of sexual maturity is reached between 1 and 1.5 years old (juveniles are <1 year old; Rotger et al. 2016, Rotger et al. 2020). Individuals were sexed by inspection of femoral pores and counting of row ventral scales (males are larger than females and show pores with visible lipophilic compounds (Salvador, 2009). Individual-based data of chest images were taken for all individuals and analyzed through the APHIS program for specimen identification and confirmation of sex (Moya et al., 2015).

After gentle pressing of the specimen abdomen, fecal drops were collected from the cloaca directly into a sterile 2 ml tube filled with 100% ethanol. Samples were placed at – 20 °C within the first 24 hours from collection and kept refrigerated until processing. Sample preservation in 95-100% ethanol was shown to be effective in maintaining microbial community composition, even for storage up to one week at room temperature (Song et al., 2016).

Stable isotopes analysis was conducted at the Laboratorio de Isótopos Estables (LIE-EBD/CSIC, Spain) on a subset of blood samples collected from 71 specimens in spring 2016. Following a 1-2 cm tail cut, drops of blood were immediately collected in capillaries, preserved in ethanol 70% and stored at −20 °C. Samples were dried for 48 hours at 60 °C and analyzed before combustion at 1020 °C using a continuous flow isotope-ratio mass spectrometry system. The isotopic composition is reported in the conventional delta (δ) per mil notation (‰), relative to Vienna Pee Dee Belemnite (δ^13^C) and atmospheric N_2_ (δ^15^N).

The species is currently listed as endangered according to the IUCN red list. Specimen sampling and manipulation were carried out in accordance with the ethics guidelines and recommendations of the Species Protection Service (Department of Agriculture, Environment and Territory, Government of the Balearic Islands), under annual permits given to GT.

### DNA extractions and 16S rRNA Illumina sequencing

Fecal samples were briefly centrifuged, ethanol removed, and the pellet used for DNA extractions with the DNAeasy Powersoil kit (Qiagen), following the manufacturer’s protocol. Samples were homogenized with 0.1 mm glass beads at 5,500 rpm, 2 × 45 sec using a Precellys Evolution instrument (Bertin Technologies). DNA quality was assessed with Nanodrop and sent to the Centre for Genomic Regulation (CRG) in Barcelona (Spain) for amplicon generation and sequencing. The region V3-V4 of 16S rRNA was amplified using a pool of five forward and reverse primers (including a frameshift to increase diversity) with Nextera overhangs (Appendix S1, Table S2). For each sample, amplicons were generated in three-replicates using KAPA Hifi DNA polymerase (Roche), with a first round of PCR (25 cycles); amplicons were then pooled and a 5 μl purified aliquot was used to seed the second PCR (8 cycles) for individual barcoding. Two negative controls (water only) and two mock communities (HM-277D and HM-276D from BEI Resources) were processed along with sample DNA. Barcoded amplicons were pooled at equimolar concentrations and the final library cleaned with the Sequal kit (Invitrogen). The library was sequenced on Illumina MiSeq v3 (600-cycle cartridge, 300 paired-end reads). The final sample dataset did not include any recaptured specimens.

### Amplicon sequence analyses

Demultiplexed sequences were input into Qiime2 (Caporaso et al., 2010), primers were removed, and reads were joined with “join-pairs” and filtered with “quality-filter q-score-joined”. Sequences were denoised with DEBLUR version 1.1.0 (trim-length=400, min-reads=5) (Amir et al., 2017) to produce Amplicon Sequence Variants (ASVs). Taxonomic assignment was performed on a trained classifier using the Greengenes database version 13_5 (McDonald et al., 2012). ASVs classified as mitochondria and chloroplasts or present in the controls (water and mock communities) were discarded (Appendix S1, Table S3). Sequences were aligned with Mafft in Qiime2, and hypervariable regions masked. Columns with gaps present in more than 50% of the sequences were removed using trimal (Capella-Gutiérrez et al., 2009). A rooted phylogenetic tree was built with FastTree (Price et al., 2009)and used for the unweighted Unifrac analysis. To limit bias in sample sequencing effort, data was rarefied to the minimum sample size (26003 sequences) and imported into the R environment using the phyloseq package(McMurdie & Holmes, 2013).

### Taxonomic composition and diversity

Taxonomic barplots were built with *ggplot* function in the ggplot2 R package. Alpha diversity was estimated according to Chao1 and Shannon indexes on seasonal datasets using the function *plot_richness* in phyloseq. Differences by islet and season were tested with two-way analysis of variance models.

Beta diversity was visually explored with principal coordinates analysis (PCoA) on Bray-Curtis distances calculated from square root transformed ASV rarefied data using function *cmdscale* in the R stats package. This distance was made euclidean by taking the square root before analysis. Differences in microbiota composition according to islet, sex, life stage, season, and year were assessed with permutational multivariate analysis of variance (PERMANOVA) on the same distance matrix after checking for homogeneity in multivariate dispersion. Model selection was performed by first fitting a model with all main terms and all two-way interactions, then refitting the model without the interaction terms with large p-values (p > 0.1) in the full model based on marginal tests with 10000 permutations. Unlike sequential tests, marginal tests evaluate each term against a model containing all other terms. Therefore, the refitted model contains tests for the chosen interactions and for the main terms that do not form part of an interaction term. PERMANOVA was done with function *anova2*, and multivariate homogeneity in dispersions with function *betadisper*, both in the R package ‘vegan’ (Oksanen et al., 2020).

### Microbiota and host genetic and trophic distances

Microbiota distances among islets were calculated as the islet centroids computed from the Bray-Curtis and unweighted Unifrac distance matrices using the function *distance* in the phyloseq package and function *dist_between_centroids* in the usedist package (Bittinger, 2020). To take into account intrapopulation microbiota variance in centroid estimates, multiple distance matrices were built on distinct core datasets (50 to 90%), where the core is a subset of ASVs shared among a cutoff percentage of individuals within a population. Host genetic distances among the three islets/populations were inferred using average *Fst* distances according to published microsatellites data(Rotger et al., 2021).

Differences in mean values of stable isotopes among islets were tested with generalised least squares (GLS) to account for strong heteroskedasticity. Post-hoc pairwise comparisons were performed using the Satterthwaite approximation for degrees of freedom and the Tukey method for p-value adjustment. Models were fitted with function *gls* in the ‘nlme’ R package (Pinheiro and Bates, 2022), and pairwise comparisons with the ‘emmeans’ package (Lenth, 2022).

### Islets microbial markers

Bacteria taxa driving differences in microbiota composition across populations (i.e., islet biomarkers) were searched through a double approach: the Dufrene-Legendre Indicator Species analysis using the *indval* function in the labdvs R package (Roberts, 2019) and the Linear discriminant analysis Effect Size (LEfSe) for biomarker discovery (Segata et al., 2011). Both approaches retrieve differences across pairs of groups considering both presence-absence and differential abundances. Results obtained from the two methods were intersected to account for potential methodological biases (Nearing et al., 2022).

For both approaches, the input dataset was rarefied, retaining only ASVs with more than 100 total sum counts to reduce sparsity issues (Nearing et al., 2022), and the dataset split into spring and autumn samples, performing the analysis on seasonal datasets. Juveniles were excluded from this analysis due to insufficient representation in the sample. *Indval* analyses were run on all taxonomic levels (from ASV to phylum), binning counts with the function *aggregate* in R stats package. Significant ASVs/taxa were retained when relfreq ≥ 0.6 (minimum relative frequency of occurrence within a population for ASV/taxa to be retained) and p < 0.01. Results from spring and autumn datasets were crossed to obtain season-independent discriminatory features. LEfSe analyses were run in the Galaxy web application setting the class to “Islet” (Kruskal-Wallis among classes p = 0.01, and pairwise Wilcoxon test between subclasses, p = 0.01), and a threshold on the logarithmic LDA set to 3, with one-against-all strategy. The analyses were run on ASV and higher taxa levels separately. Only ASV/taxa retrieved in both seasonal datasets were retained as islet biomarkers.

### Persistence of ASVs over sampling dates and seasonal microbial markers

Recurrent occurrence of the same microbial ASVs over time, here referred to as persistence, was evaluated on the two major islets NG and NM. For each islet, we subsetted the data according to “Date” (season-year) and for each subset we estimated the 50% core ASVs. The four core datasets were then compared to retrieve common ASVs, i.e., the microbial component present along the four sampling dates. Comparing the 50% core by date, instead of using the full microbial diversity per date, reduces the probability that only a few specimens per population contributed to the observed pattern. Venn diagrams were produced using the online tool at http://bioinformatics.psb.ugent.be/cgi-bin/liste/Venn/calculate_venn.htpl.

To estimate bacterial features responsible for seasonal differences within a population (i.e. seasonal markers), we undertook a similar double approach, performing both LEfse and indval analyses. The analyses were run on NM and NG, which had large sample representation for both autumn and spring 2017 and 2018 (above 15 samples each season/year). Same season samples within each population were treated as a single group and discriminatory features were estimated as above (for Lefse analysis, class was set to “Season”).

## Results

We sequenced the fecal microbiota of 109 specimens from the three closed populations/islets south of Mallorca: Na Moltona (NM), Na Guardis (NG), and En Curt (EC) (Figure 1). Fecal samples were associated to four major categorical variables: islet, sex, life stage, season (spring and autumn) and year (2017 and 2018) (see Appendix S1, Table S1 for sample metadata). The final microbiota dataset encompassed an even representation of each variable, except for life stage (nearly 80% of the specimens were adults).

After extensive quality filtering and removal of taxa found in the controls (Appendix S1, Table S3), we obtained a total of 6195163 sequences and 2313 ASVs (10 minimum reads) (abundance matrix with taxonomy is available at Mendeley Data, doi: 10.17632/bc5nxsxgxd.1): 1360 ASVs in EC, 1647 in NG and 1677 in NM. Of these ASVs, 91 (EC), 74 (NG) and 70 (NM) were present in at least 80% of the specimens within each islet/population (i.e., they form the core microbiota). According to rarefaction curves, sequencing effort was sufficient to approach the maximum diversity for most samples (Supporting Information, Figure S1). Data was nonetheless rarefied to the minimum sample depth of 26003 reads (corresponding to 1933 ASVs) to account for potential bias in sequencing effort and sparsity, and used for all subsequent analyses.

### Highly conserved microbial taxonomic profile among wall Lilford’s wall lizard allopatric populations

The overall fecal microbial dataset comprises a total of 18 unique phyla, 36 classes, 64 orders, 94 families, 134 genera, and 66 species. The taxonomic profile of the most abundant taxa was remarkably conserved at phylum, family, and genus level across all individuals (Supporting Information, Figure S2), with no major compositional differences across islets, between males and females and along the four sampling dates (Figure 2). In all cases, the two most abundant phyla were Firmicutes (43%) and Bacteroidetes (38%), with similar relative abundances, followed by Proteobacteria (8%) and Tenericutes (6%) (Figure 2). Dominant families were Bacteroidaceae (21%), Lachnospiraceae (15%), Ruminococcaceae (8%) and Porphyromonadaceae (8%). The most abundant genera were *Bacteroides* (21%), *Parabacteroides* (7%), *Anaeroplasma* (5%), *Oscillospira* (4%), *Odoribacter* (3.5%) and *Roseburia* (2.6%) (Figure 2). Only 8% of the ASVs (154 out of 1933) reached species classification (confidence threshold 80%); the most abundant species were *Parabacteroides gordonii* (4.4%), *Clostridium ramosum, Parabacteroides distasonis* and *Akkermansia muciniphila* (all <1%).

**Figure 2:**
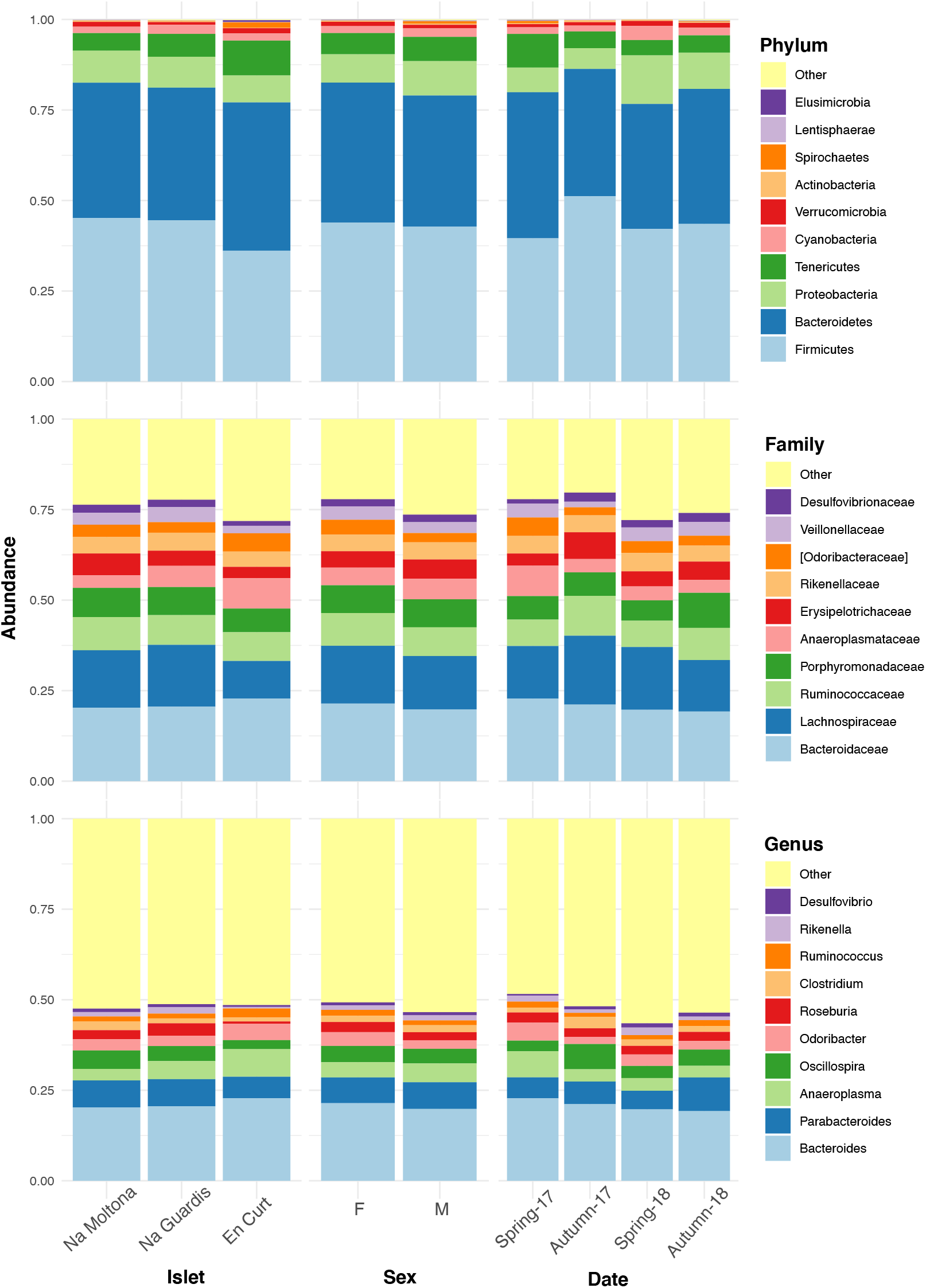
Taxonomic composition of the *P. lilfordi* gut microbiota at phylum, family, and genus levels according to “islet”, “sex” and sampling “date” (season-year). Juveniles (n = 9) were excluded. The legends list only the top 10 taxa (all remaining taxa were included in “Others”). No major taxonomic differences were observed as a function of any of the variables. For individual specimen taxonomic profile see Supporting Information, Figure S2.

### Islet and season as major variables shaping the microbiota structure

PERMANOVA analysis on the entire dataset (i.e., including all life stages, i.e., juveniles, subadults and adults) indicated statistically significant clustering by the interaction between islet and season (P ≤ 0.0001), and islet and life stage (p = 0.005), marginal differences by year (P = 0.0449), and no sex effects (P = 0.1197) (Table 1a). This suggests potential ontogenetic differences in gut microbiota. However, juveniles were underrepresented in our dataset (n = 9 out of 109 individuals). When excluding juveniles, PERMANOVA analysis shows again strongly significant islet by season interactions (P ≤ 0.0001) and marginal year effects (P = 0396), yet no differences by either sex or life stage (Table 1b). This suggests that differences in life stage were mostly due to the juvenile stage, with no differences between adults and subadults. Therefore, juveniles were excluded from all subsequent analyses.

**Table 1:** Results of PERMANOVA (9999 permutations). All tests are marginal.

|  | Df | Sum of squares | R<sup>2</sup> | pseudo F | Pr(>F) |
| --- | --- | --- | --- | --- | --- |
| *a) Entire dataset* |  |  |  |  |  |
| Sex | 2 | 0.658 | 0.0183 | 1.1061 | 0.1197 |
| Year | 1 | 0.376 | 0.0104 | 1.2641 | 0.0449 |
| Islet:Season | 2 | 0.998 | 0.0277 | 1.6778 | **0.0001** |
| Islet:Life_Stage | 6 | 2.099 | 0.0583 | 1.1759 | **0.0050** |
| Residual | 94 | 27.970 | 0.7760 |  |  |
| Total | 108 | 36.043 | 1 |  |  |
| *b) Without juveniles* |  |  |  |  |  |
| Sex | 1 | 0.309 | 0.0095 | 1.0432 | 0.2876 |
| Life_Stage | 1 | 0.309 | 0.0095 | 1.0433 | 0.2957 |
| Year | 1 | 0.379 | 0.0117 | 1.2818 | 0.0396 |
| Islet:Season | 2 | 1.050 | 0.0323 | 1.7744 | **0.0002** |
| Residual | 91 | 26.921 | 0.8270 |  |  |
| Total | 99 | 32.552 | 1 |  |  |

Principal coordinates analysis showed that EC hosted a clearly distinct microbial community, while NG and NM substantially overlapped on the subspace defined by PCo1 and PCo2 (Figure 3). In addition, post-hoc tests by islet showed that season was a statistically significant factor in every case, but most clearly in Na Moltona and Na Guardis (NM: P = 0.0002, NG: P ≤ 0.0001, EC: P = 0.008) (Figure 3).

**Figure 3:**
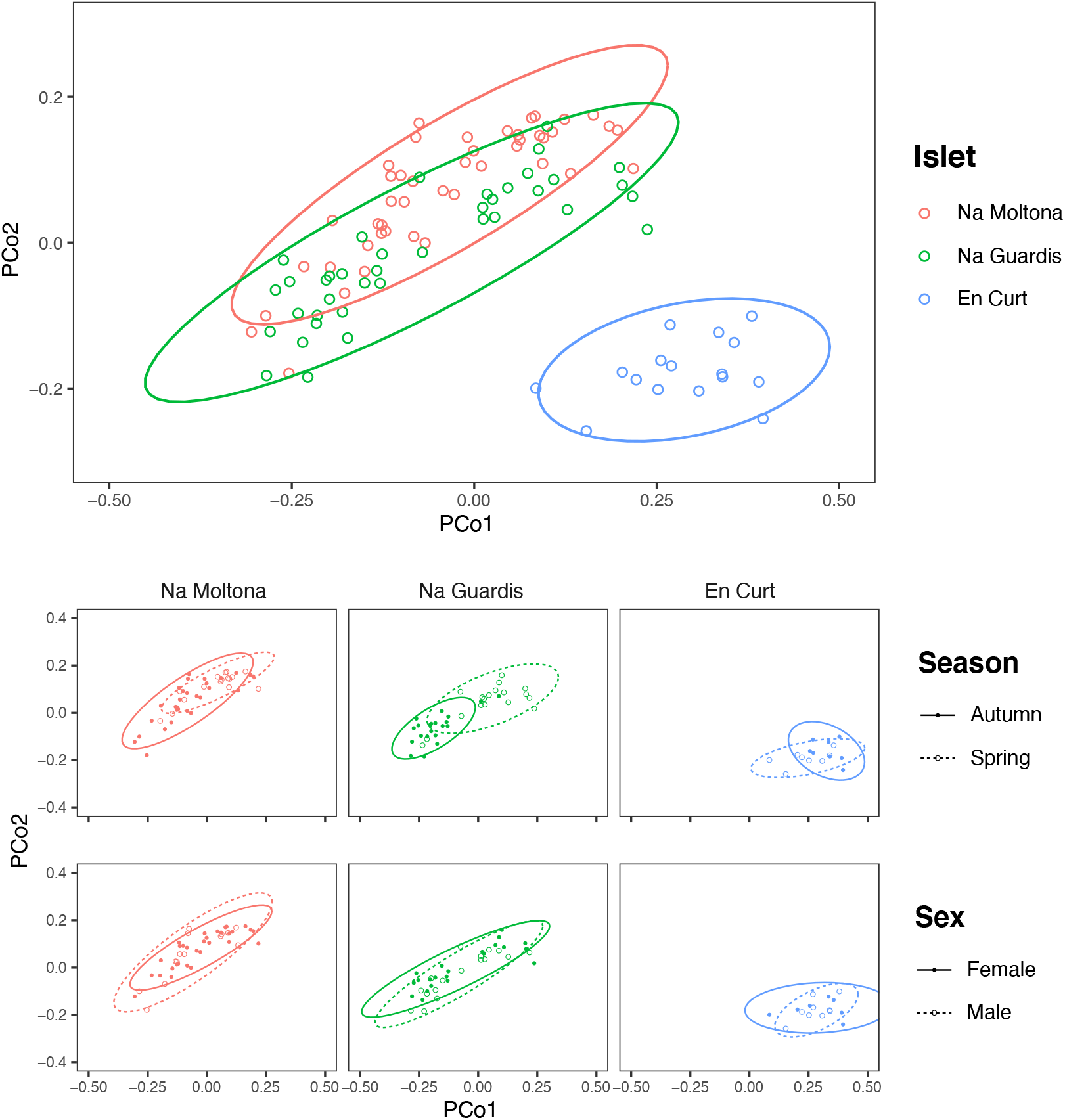
PCoA based on Bray-Curtis distances of the *P. lilfordi* gut microbiota depicting diversity among populations (“islets”) and within populations, according to “season” and “sex”. Dots represent specimens. Juveniles (n = 9) were excluded. Ellipses (calculated with stat_ellipse in ggplot2) enclose 95% of the expected values around centroids assuming a *t* distribution. Data were square root transformed. Microbiota differences were driven by “islet”, “season” (within each islet), but not “sex”.

According to alpha diversity analyses on seasonal datasets (Figure 4), spring showed a highly homogenous pattern of diversity, with no major differences across any islet pairwise (both Chao1 and Shannon, p > 0.05), whereas autumn marked a large difference among all three populations, with EC being the most diverse (Chao1 and Shannon, p< 0.05 for all islet pairwises, except for NG-EC, p > 0.05 Shannon). Within individual populations, spring showed a richer community in NG (p < 0.001 for Chao1, but not significant for Shannon) but not in NM (p > 0.05 both indexes), while the opposite pattern was observed in EC, with autumn being most diverse (p < 0.05 both indexes). No statistically significant differences in alpha diversity were found between sexes within individual islets (p > 0.1 for both indexes).

**Figure 4:**
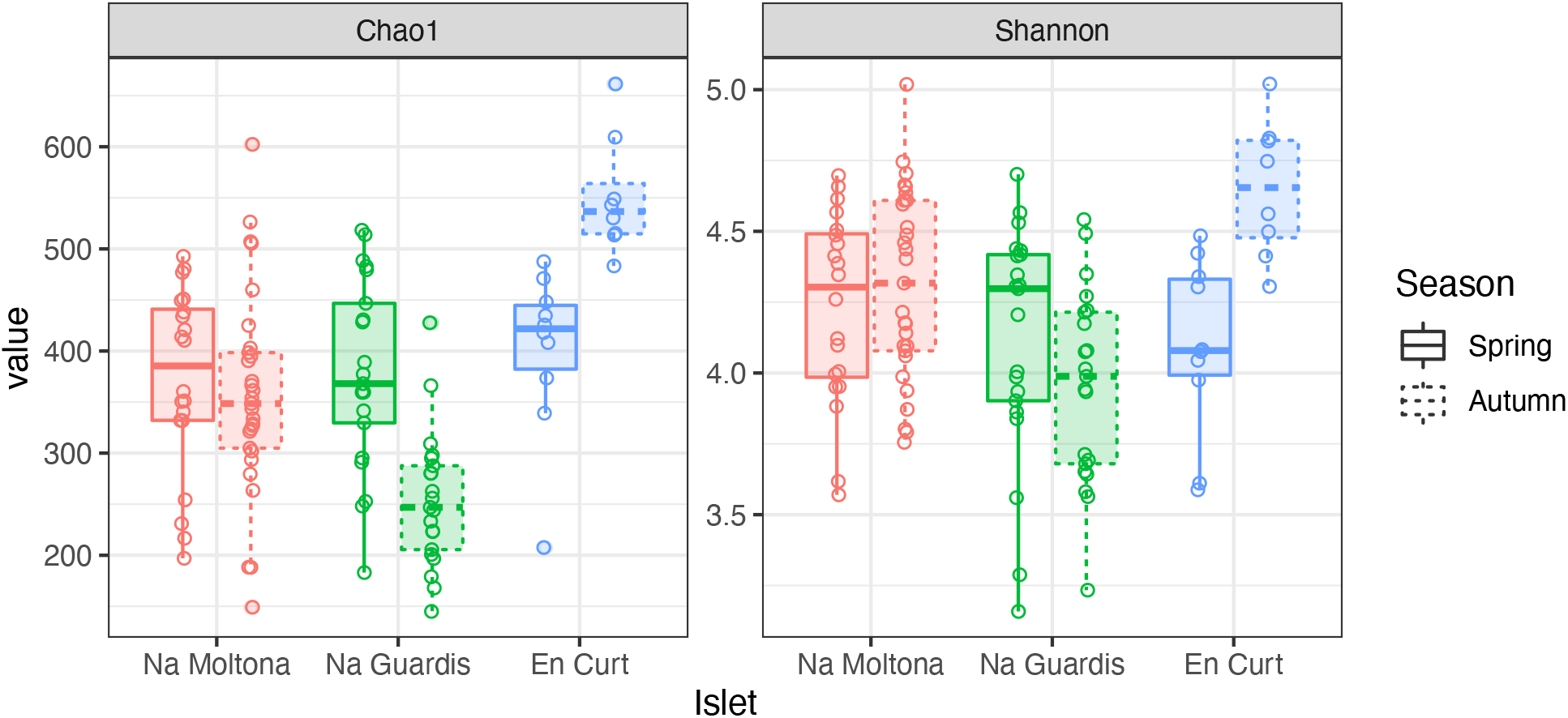
Gut microbiota alpha diversity by islet according to Chao1 and Shannon, estimated on seasonal datasets. Differences among islets are observed only for the autumn dataset.

### Among population microbiota diversity is explained by both lizard phylogeography and trophic niche

To explore the “islet/population” effect on the microbiota diversity (Figure 3 and 4), we searched for biomarkers of each islet using a double approach (*indval* and Lefse analyses, see Methods). A total of 14 ASVs and two taxa were retrieved by the two methods, which discriminated across islets according to both autumn and spring datasets (Figure 5 and Appendix S1, Table S4 for results and taxonomic classification). In accordance with the PCoA clustering (Figure 3), most discriminatory features were enriched in the EC islet and virtually found only on this islet, with no occurrence in either NM or NG (relabund values close to 0 for both NM and NG, Table S4). Only three ASVs were found to be specifically enriched in NM, although not unique to this islet, while NG showed no islet-specific microbial markers. Most discriminatory ASVs belonged to the order Bacteroidales, and fermentative families Bacteroidaceae and Porphyromonadaceae. Their abundance across individuals was highly comparable between spring and autumn, suggesting stability in biomarkers relative abundance over time (Figure 5). At taxa level, the islet EC showed a unique enrichment in the phylum Elusimicrobia (virtually absent in the other islets) and in the genus *Vibrio*, specifically in the species *Vibrio rumoiensis* (phylum Proteobacteria) (Figure 5). No specific taxa markers were detected for either NG or NM.

**Figure 5:**
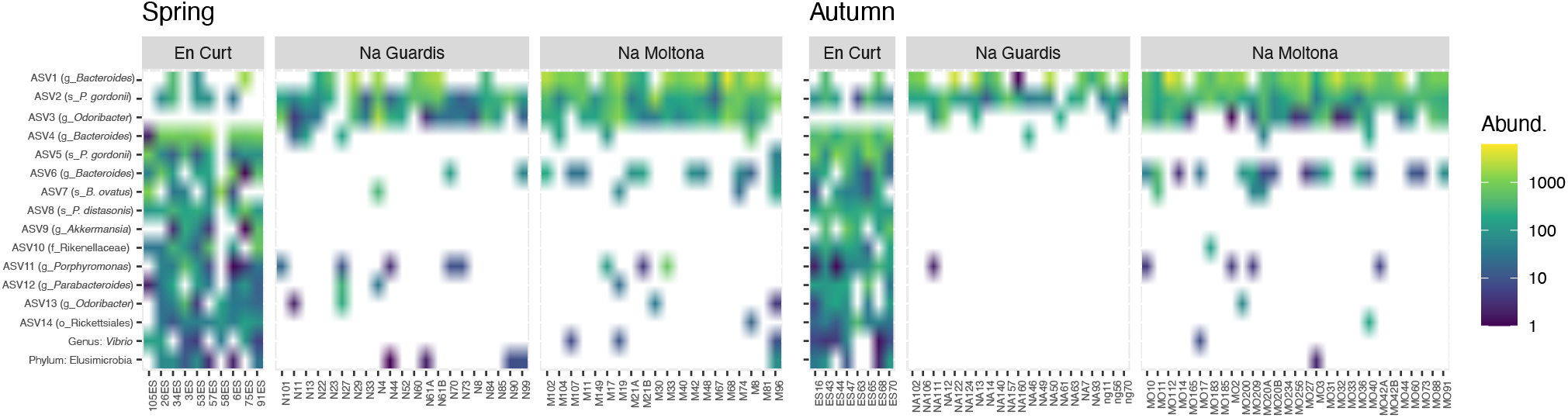
Heatmap of the microbial markers (14 ASVs and two taxa) driving differences across islets, consistently in spring and autumn. Pattern of ASV relative abundance per specimen (x axis, ordered by their mean abundances in spring) is highly concordant between seasons, despite the datasets include different sets of individuals. Heatmaps were built on log-transformed data grouped by islet and season. Data were restricted to ASVs and taxa retrieved by both indval and LEfSe approaches (for indavl, relfreq ≥ 0.6 and p < 0.01; for LEfSe LDA > 3 and p < 0.01). See Appendix S1, Table S4 for full taxonomic classification and statistics.

Bray-Curtis microbiota centroid distances among islets, calculated using distinct core subsets per islet (50, 60, 70, 80%), indicated high microbial community relatedness between the two largest islets, NG and NM, with EC being the most differentiated (Figure 6A, see Figure S3 for Unifrac distances). These microbiota distances were concordant with the host population genetic distances based on previously estimated *Fst* values of microsatellite diversity (Rotger et al., 2021).

**Figure 6:**
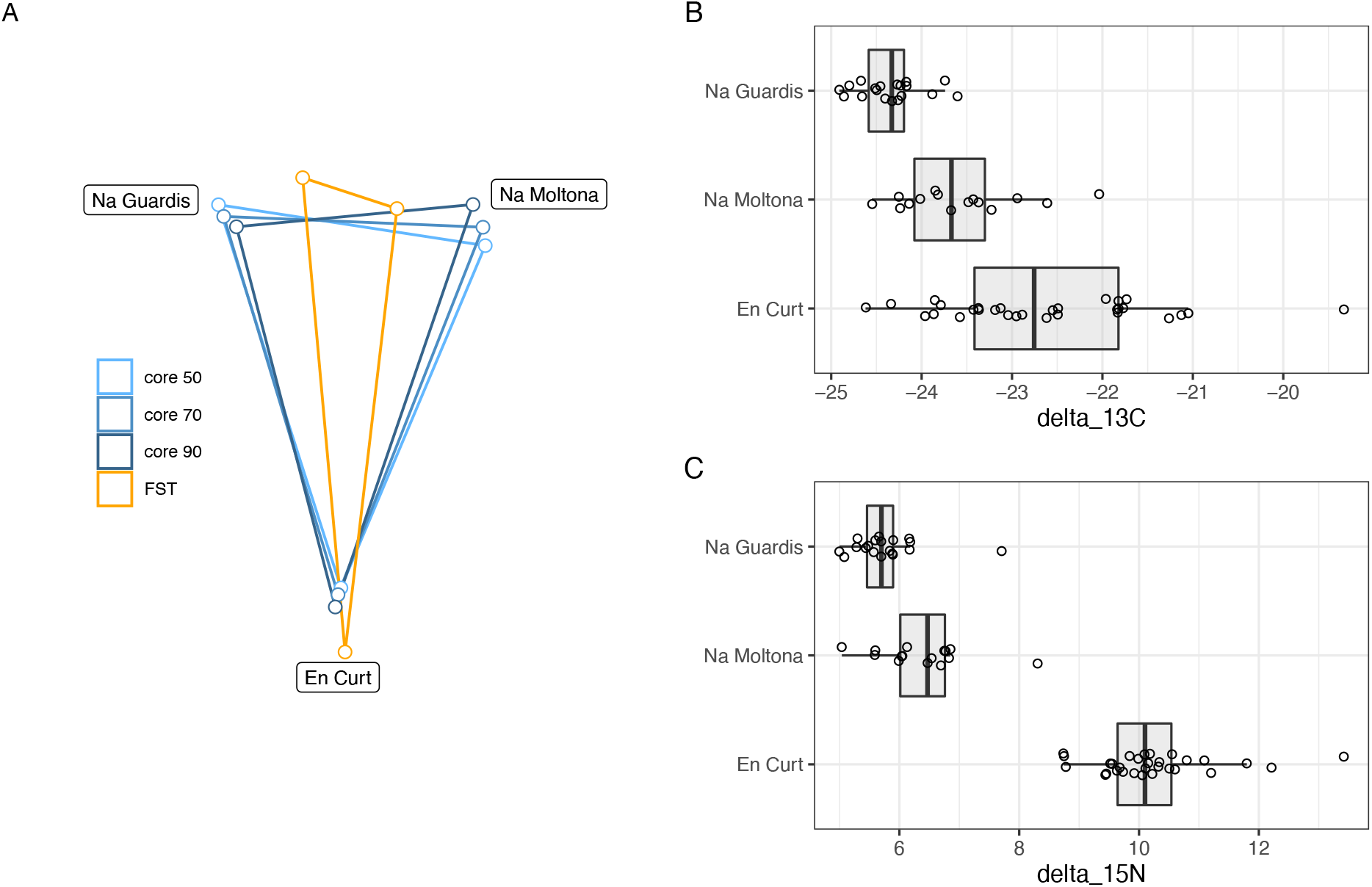
Host genetic and gut microbiota distances among the three islets/populations (A) and host trophic niches (B). A) Superimposed map of microbiota centroids distances (Bray-Curtis) per islet calculated on different core subsets, and host population genetics distance based on *Fst* values estimated on available microsatellites from a previous study (pairwise *Fst*, EC-NM: 0.135, EC-NG: 0.144, NM-NG: 0.03, p=0.001 for all pairwises) (Rotger et al., 2021). See Supporting Information, Figure S3 for results based on Unifrac distances). B) Stable isotopes estimated for each population based on a dataset from spring 2016 (Appendix S1, Table S5). EC displayed higher values of both carbon-13 and particularly of nitrogen-15, with minor differences among NM and NG. The relative distances among islets according to host genetics, trophic ecology and gut microbiota are highly congruent.

Trophic niche distances among the three populations were investigated through stable isotopes on sample sets from 2016 (data available at Appendix S1, Table S5). Findings indicated that both carbon-13 and nitrogen-15 differed among islets (Figure 6B), while post-hoc pairwise analyses showed that in both cases EC displayed higher values of both carbon-13 and particularly of nitrogen-15 with respect to both NG (N-15: t[46.8] = 19.7, P < 0.001; C-13: t[37.1] = 7.57, P < 0.001), and NM (N-15: t[35.9] = 14.38, P < 0.001; C-13: t[41.8] = 3.44, p = 0.038), while the latter islets differed for carbon-13 (t[20.1] = 4.01, P = 0.0019) but only marginally for nitrogen-15 (t[25.8] = 2.58, P = 0.0407).

Overall, the microbiota distances were consistent with both the host population genetic and ecological distances.

### Within island/population microbiota diversity: microbiota persistence and seasonal effect

Microbial diversity within islets was primarily driven by “season” (p = 0.001, no juveniles) (Table 1 and Figure 3). To evaluate the level of microbiota plasticity within a population over time (i.e., degree of changes in relative abundance and/or turnover of microbial taxa), we investigated both persistence of microbial ASVs along the four sampling “dates” (spring-17, autumn-17, spring-18 and autumn-18) and enrichment patterns as a function of season (spring vs autumn). The analyses were restricted to the two islets with the largest sample representation per date, NM and NG (Figure 1).

Persistence was assessed as the portion of the microbial ASVs that was consistently retrieved in all four sampling dates, considering only those ASVs that occurred in at least 50% of the specimens within a single date. Of this total core ASV diversity (441 for NM and 334 for NG), 30.5% (102 ASVs for NG) and 49% (151 ASVs for NM) were recurrent in all four dates (Supporting Information, Figure S4 and Appendix S1, Table S6 and S7), with 82 ASVs being common to both islets. Taxonomic profiles of these persistent ASVs were largely congruent between islets, with the majority belonging to the family Ruminococcaceae and genus *Oscillospira*, followed by members of the Lachnospiraceae and Bacteroidaceae (particularly of the genus *Bacteroides*) (Figure 7).

**Figure 7:**
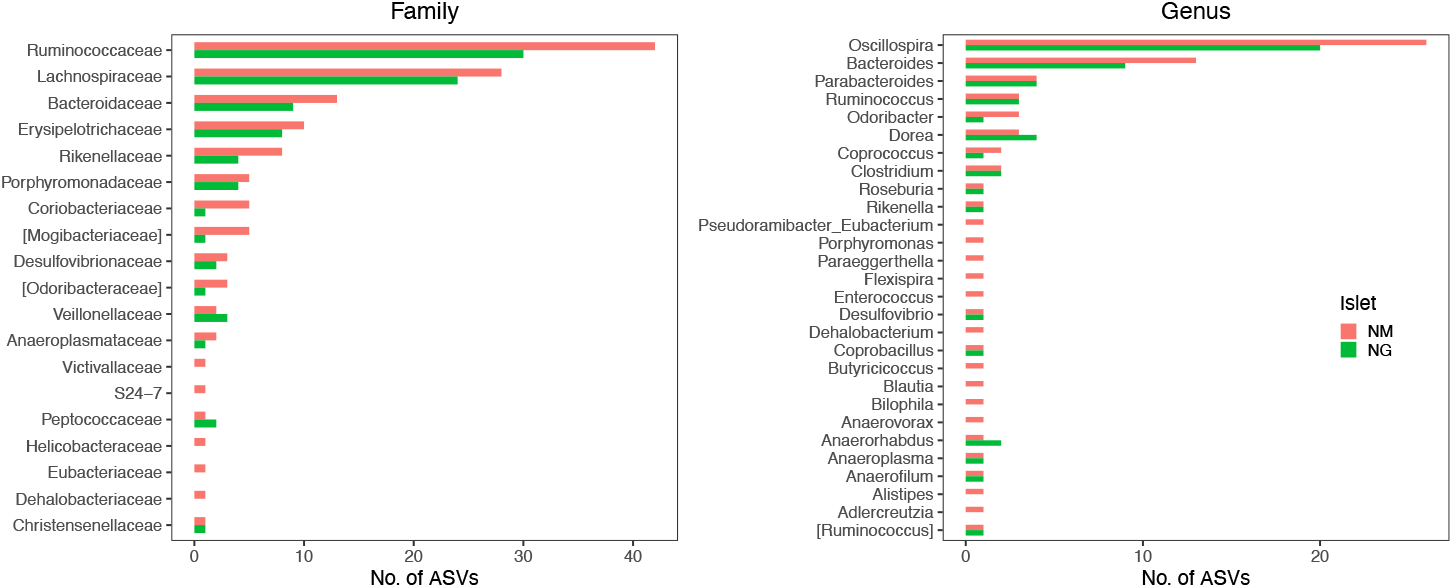
Family and genus-level taxonomic profile of persistent ASVs retrieved along the four sampling dates. The bars indicate the frequency of ASV per each taxon. The two islets shared a highly similar taxonomic profile.

According to microbiota centroids by “Date” on Bray-Curtis distances, same season microbiotas sampled in 2017 and 2018 were highly similar and diverged from microbiotas collected in distinct seasons (Figure 8A), indicating that the microbiota structure alternates across seasons in a quite conserved manner: the microbiota configuration state shifts from spring 2017 to autumn 2017, then goes back to a similar state in spring 2018 and finally shifts again to the autumn configuration in 2018. This pattern was observed in both islets and was robust to the use of different core subsets, up to 90% (Figure S5), suggesting it is largely driven by quantitative changes in relative abundances of core ASVs.

**Figure 8:**
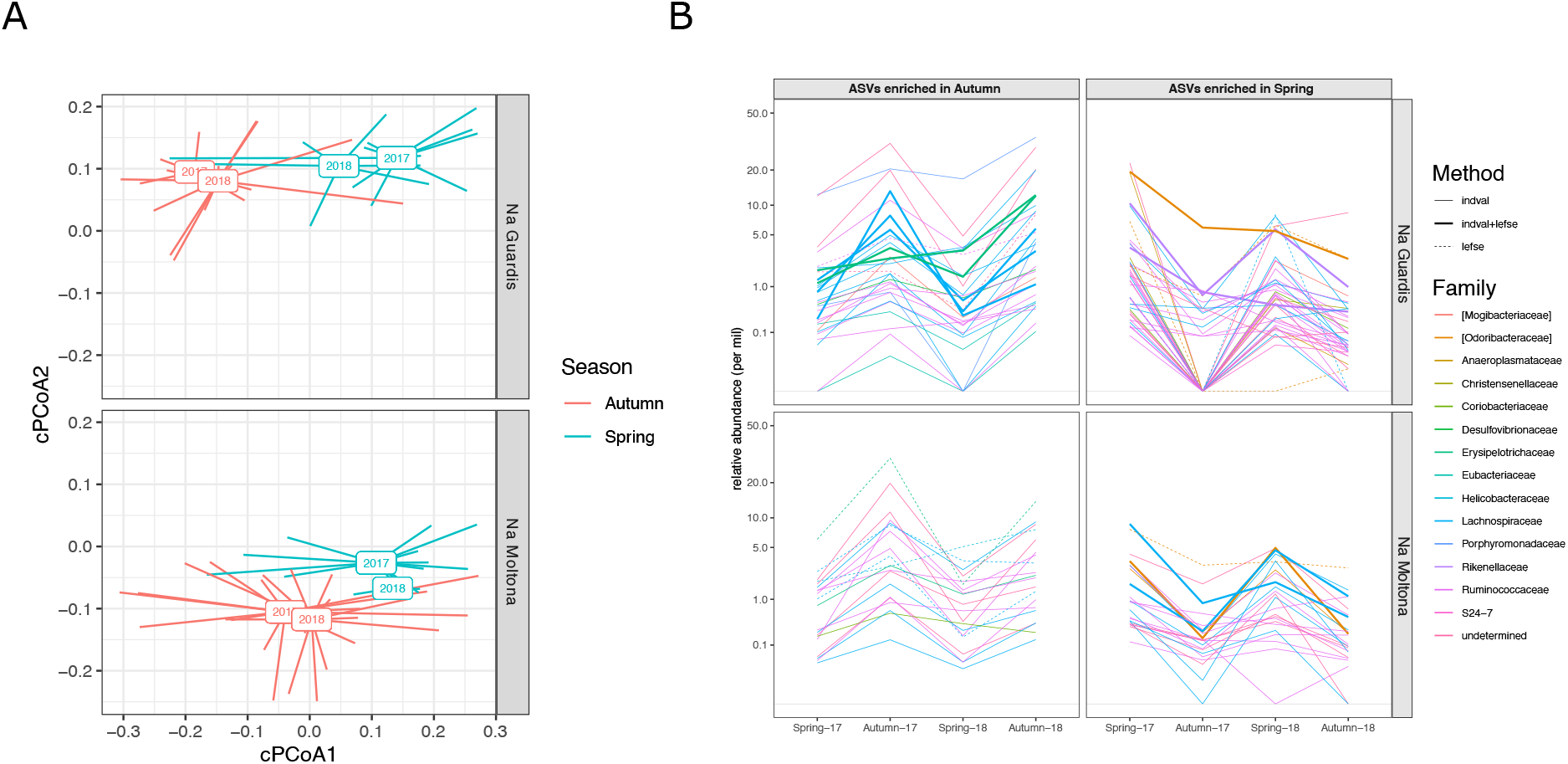
Microbiota seasonal changes (spring and autumn) in NG and NM along two years sampling (2017-18). A) PCoA based on Bray-Curtis distances on square root transformed values. Square boxes depict centroids for each year and season, with lines connecting centroids with individual observations. Microbiota configuration changes across seasons in a repetitive manner and consistently in the two populations. Results were robust to the use of different core subsets (see Supporting Information, Figure S5). B) Variation in mean relative abundance along “dates” of ASVs that were significantly enriched in either spring or autumn according to LEfSe and/or indval analyses. A clear pattern of seasonal fluctuations can be observed for most ASVs, with the majority belonging to the families Ruminococcaceae and Lachnospiraceae. A total of 20 ASVs (NG) and 11 ASVs (NM) were consistently retrieved by both methods (see also Appendix S1, Table S8).

Enrichment analyses with *indval* and Lefse analyses identified 63 and 105 ASVs that were differentially abundant between spring and autumn in NM and NG, respectively (of these, 20 ASVs (NG) and 11 ASVs (NM) were consistently retrieved by both methods, Table S8). Most of these ASV seasonal markers (spring and autumn enriched) showed a clear fluctuation in relative abundance along the four sampling dates, for both islets (Figure 8B). A proportion of them corresponded to persistent ASVs (7 of the 20 ASVs in NG, and 5 of the 11 in NM), with the majority showing an average abundance different from zero for all dates (see in particular NM autumn-enriched ASVs) (Figure 8B). Only few ASVs dropped below the threshold of detection during one/two dates (particularly in NG, Autumn-17), while being present on all other dates. The taxonomic profile of enriched ASVs was largely similar between seasons and islets: most ASVs (~70%) belonged to the order Clostridiales and family Ruminococcaceae and Lachnospiraceae (Figure 8B).

A similar pattern was retrieved for higher taxonomic levels, with most taxa being consistently retrieved across all dates, while fluctuating in relative abundances in both islets (Supporting Information, Figure S6). Notably, few species showed a clear seasonal-associated presence/absence pattern, including the disease-associated species *Enterobacter hormaechei* and *Lawsonia intracellularis* (autumn-specific for NG). While the taxonomic composition of enriched taxa was overall largely specific to each islet, few taxa displayed a highly congruent pattern between islets, supported by both methods (LEfse and indval): the genus *Odoribacter* and family *Odoribacteraceae* (enriched in spring), and the genus *Anareofilum* (enriched in autumn). Both taxa form part of the core microbiota (present in at least 80% of all specimens).

## Discussion

A fundamental question in the study of the host-microbes symbiosis is to which extent this association is resilient to spatio-temporal changes and what are the processes influencing such (lack of) divergence, which ultimately affects patterns of coevolution in animals (Mallott & Amato, 2021). The use of relatively simple natural systems (small closed populations) and individual-level data are critical to address this question.

Here we explored the trajectories of early diversification of the gut microbiota among and within three sister islet populations of the Balearic wall lilford’s lizard, focusing on the impact of host phylogeographic history, dietary niches, host intrinsic traits (sex and lifestage) and temporal variables (year and season), with the major aim of sheding light on the strength of this symbiotic association, its level of plasticity and putative role in the host metabolic adaptation.

### Microbiota diversity across islet populations

What drives the early steps in microbiota diversification among populations once the reproductive boundaries are set? And in which aspects do these associated communities start diverging following the host genetics and adaptation to the new environment?

Gut microbiota divergence across host allopatric populations is largely a function of timing since population divergence (i.e., the phylogeographic history), putative adjustments/transitions in the host ecological niche following separation, and exposure to distinct pools of environmental bacteria (Lankau et al., 2012; Michel et al., 2018). The relative impact of these processes on microbial community changes will depend on the plasticity of the gut microbiota in response to both host selectivity/filtering (underpinned by the host genetics) (Alberdi et al., 2016; Gomez et al., 2019), the level of microbial transmission across generations (both vertical and horizontal) and the impact of stochastic events (i.e., ecological drift) (Baldo et al., 2018; Lankau et al., 2012).

Here we observed a largely homogeneous taxonomic profile of the gut microbiota among three populations, without any major differences according to the studied variables (Figure 2 and Supporting Information, Figure S2). This is largely consistent with our previous study based on microbial content of seven populations of the same species from Menorca, using full intestine tissues (Baldo et al., 2018), with both Menorca and Mallorca populations presenting a similar dominance of the phyla Firmicutes and Bacteroidetes, families Lachnospiraceae, Ruminococcaceae and Porphyromonadaceae and genera *Bacteroides* and *Parabacteroides* (Figure 2). This conservatism of the taxonomic profile likely results from common host genetic constraints (Mallott & Amato, 2021), exerting a strong imprinting and stabilising effect on the major microbial membership composition, which overcomes the exposure to distinct local environments (Baldo et al., 2018; Rotger et al., 2021), a pattern that has been previously observed in recently diverged species (e.g., in the Galapagos finch (Michel et al., 2018).

Nonetheless, at finer taxonomic level, i.e., in terms of ASVs, the three islet populations carried their unique microbial signature (Figure 3). Once excluding the effect of life stage and season (therefore working only on adults/subadults and individual seasons), islet was indeed a statistically significant clustering factor, driven by a limited number of ASVs and taxa, mostly specific to the smallest islets EC (Figure 4). Islet biomarkers had comparable enrichment patterns across the spring and autumn datasets (showing independence from a seasonal effect) and might represent new uptakes, although limited, from the local microbial pool. Their functional role to the host adaptation is worth further investigation. In particular, the phylum Elusimicrobia was largely specific of EC and represents a recently identified animal-associated phylum which rely on fermentation (Méheust et al., 2020).

Microbiota distances among the three populations/islets were consistent with their host population genetic distances, suggesting a phylogeographic scenario of microbiota divergence following their host diversification (Figure 6A). According to published microsatellites data, NM and NG diverged about 2,000 – 4,000 years ago, with EC representing the most distant population (Rotger et al., 2021). NM and NG are the geographically and genetically closest populations and hosted the most similar microbial communities (Figure 1 and Figure 6A). A putative event of gene flow might have occurred between NM and NG in recent times (< 200 years, associated to a single specimen translocation) (Rotger et al., 2021), which could have resulted into a potential dispersion of microbial taxa and a homogenization of the overall microbial diversity of these two populations. However, these two islets show clear differences in their morphological and life history traits (smaller body size and marked senescence in NG, Rotger et al. 2020), indicating an ongoing process of divergence between the two islets (Rotger et al., 2020, 2021). Their closed microbiota similarity and divergence from the the islet EC might still be compatible with a process of codivergence of microbes with the host over time, as observed in several instances across the animal kingdom (Lim & Bordenstein, 2020; Mallott & Amato, 2021). Comparisons of captive *versus* natural populations have indeed proved the ability of lizards to transmit microbes across generations (Colston, 2017; Kohl et al., 2017), suggesting a scenario of putative retention of ancestral bacteria following specimen segregation during the wall lilford’s lizard vicariance process (Baldo et al., 2018).

Whilst microbial communities might carry a phylogeographic signal, the analysis of the trophic niche by stable isotopes provided a seemingly compatible clustering: EC deviates in its trophic niche from both NG and NM, probably due to the rocky nature of this small islet (0.30 ha) (Figure 1) with a very low biotic index and limited resources, causing metabolic stress and a higher incidence of cannibalism. A wider range of both carbon-13 and nitrogen-15 values for this population (Figure 5B) indicate a larger trophic niche breadth, putatively including also marine food items obtained along the shore (molluscs and crustaceans), not predominant in either NM or NG (personal observation of behavior). Dietary adjustments can greatly impact the microbiota composition in lizards (Jiang et al., 2017; Kohl et al., 2016; Li et al., 2019; Montoya-Ciriaco et al., 2020), suggesting that the differences in trophic niche among populations can partly drive the observed microbiota clustering.

At present, both the phylogeographic and ecological scenarios are compatible with the observed pattern of islet microbiota divergence, inviting caution in data interpretation and especially in phylosymbiosis claims when ecological aspects are not fully understood. A selection of putatively heritable gut bacteria is currently being analysed at strain level to resolve the evolutionary trajectories of gut microbes in the three populations. Additionally, a more in-depth study of the trophic niche should be undertaken to clarify major ecological/dietary differences across populations.

### Microbiota diversity within populations: lifestage effect and no sex differences

Dissecting the drivers of microbial diversity within natural populations presents several challenges due to the multiple concurrent host variables to consider (e.g., sex and life stage), and the spatial-temporal effect (e.g., year and season). To date, the few available studies on lizard microbiota have indeed targeted variation between natural populations (Kohl et al., 2017; Ren et al., 2016; Zhang et al., 2018), with few of them looking at limited intrapopulation diversity aspects (Kohl et al., 2017; Ren et al., 2016).

The sampling design in the present study allowed us to statistically explore the relative contribution of major players in microbiota diversity within populations. This diversity was driven by temporal variables (season, but not year) and partly life stage, but not by sex (Table 1 and Figure 3). Statistically significant differences between juveniles (<1 year old) and both adults and subadults suggest a change in microbiota composition during the first year of lizard development, putatively associated with niche adjustments (Pérez-Mellado et al., 2015), a preliminary finding that warrants further investigation.

According to previous studies in *P. lilfordi* showing sex-specific diet and niche adjustment along the year (Pérez-Mellado et al., 2015; Santamaría et al., 2019), we also expected a sex and seasonal effect on the gut microbiota. In general, lizards in Mediterranean islands are largely omnivorous and opportunistic in trophic behavior, modulating niche width and food preferences along the year in response to both resource availability and energy requirements (affected by reproductive behavior and external temperature) (Pérez-Cembranos et al., 2016). This transition in the trophic niche is particularly strong between spring (with the highest resource availability) and autumn (lowest), with hot summers marking a progressive limitation of resource availability (Pérez-Cembranos et al., 2016). According to a recent study based on individual fecal content analysis in NM and NG, the *P. lilfordi* seasonal response is sex specific (Santamaría et al., 2019), and particularly noticeable in autumn, when food resources are scarce and males show a despotic behavior (Rotger et al., 2020), restricting the female niche amplitude (Santamaría et al., 2019). Despite a sex influence on both lizard metabolism and trophic niche behavior (Pérez-Mellado et al., 2015), males and females did not carry distinct gut microbiotas in any of the three islets, nor were microbial differences observed between sexes within a season (season-by-sex interaction was not significant for individual islets). These results agree with findings in other lizard species of the family Liolaemidae, where no detectable effects of sex were observed (Kohl et al., 2017). In *P. lilfordi*, a lack of sex effect could be associated to an omnivorous diet (with no clear sex-specific food preferences), a reduced sexual dimorphism (Rotger et al., 2020)and a large microbial metacommunity effect within the discrete boundaries of an island (Miller et al., 2018). Microbes can be largely transmissible in highly social or closed populations, due to the increased probability of contact among specimens (Raulo et al., 2021; Tung et al., 2015), allowing the rapid circulation of bacteria within the population. Furthermore, episodes of cannibalism are known within the genus *Podarcis* due to the restricted resources available in the islands (Cooper et al., 2015). Both sociality and cannibalism could provide a means of bacteria transmission between sexes, resulting in microbiota homogenization.

### Temporal microbiota variation within a population/islet: persistence and seasonal plasticity

Once excluding the life stage effect (i.e., restricting the analyses to subadults/adults only) and pooling males and females, within-population microbial diversity was largely driven by season. To understand the short-term temporal dynamics of the gut microbiota within a population and the strength of the host-microbes association, we explored levels of microbiota plasticity along the two years sampling.

An important fraction of the microbiota (at least 30% of ASVs) persisted within a population over the four sampling dates. This is likely an underestimate once considering a putative failure in ASV sequencing for some of the specimens. As our study did not compare the same set of individuals over time, such persistence should be considered as the maintenance of specific ASVs within the host population microbial metacommunity, not at individual level (Robinson et al., 2019). While we cannot exclude a contribution of the environmental allochthonous component derived from diet, we note that a 50% minimum ASV frequency occurrence among specimens within each sampling date largely excludes a stochastic microbial contribution from diet. This is in line with several studies spanning a wide range of animal taxa and consistently showing a neglectable contribution of the “external” microbiota (Costello et al., 2010; Kohl et al., 2017). Although the diet-derived microbial content of Lilford’s wall lizard has yet to be characterized, the lack of microbial sex-specific differences, whereas diet is largely sex-specific (Santamaría et al., 2019) further suggests that the impact of the allochtonous component might be minor. Future in-depth characterization of the environmental microbiota, including the phyllosphere, will provide a necessary confirmation.

Interestingly, the taxonomic profile of these persistent ASVs was highly congruent between islets (Figure 7), with most taxa being previously identified as highly heritable in vertebrates, including the fermentative families Lachnospiraceae and Ruminococcaceae (Grieneisen et al., 2021 in wild baboons) and the genera *Oscillospira, Bacteroides, Odoribacter, Anaerotruncus* and *Coprobacillus* in lizards (Kohl et al., 2017). The majority represent important metabolic players (Gophna et al., 2017; Kohl et al., 2016, 2017) and are known for their ability to degrade vegetable fibres (particularly the genera *Oscillospira and Bacteroides*) (Gophna et al., 2017; Patnode et al., 2019). While lizards from these islands predominantly consume arthropods, particularly insects, they are known to partly feed also on plant material (including seeds, nectar and pollen) (Pérez-Cembranos et al., 2016; Santamaría et al., 2019)as a derived adaptation to the limited environmental resources (van Damme, 1999). Presence and persistence of these core fermentative bacteria would support a putative role of the gut microbiota in extending the lizard trophic niche amplitude towards the consumption of vegetable matter, an intriguing hypothesis that will need to be corroborated with a target study on metabolic contribution of gut microbes.

We finally looked at compositional changes that the microbiota undergoes along seasons (i.e., seasonal plasticity). Recent studies on humans, wild great apes and mice (Baniel et al., 2021; Guo et al., 2021; Hicks et al., 2018; Maurice et al., 2015; Smits et al., 2017) have shown seasonal cycling of the gut microbiota in response to diet, suggesting that the microbiome not only can optimise energy metabolism for specific diets, but can also confer dietary flexibility to their hosts in response to physiological needs and changes in resource availability over time (Alberdi et al., 2016). To date, this seasonal effect has only been marginally explored in reptiles (Kohl et al., 2017).

Our findings indicated a clear seasonal shift in the gut microbiota configuration, which replicated along the two sampled years according to multiple core microbial subsets (Figure 8A and Supporting Information, Figure S5). This microbial temporal pattern was observed for both NM and NG populations, with a strong seasonal correspondence between islets, which is in line with their similar genetic background (Rotger et al., 2021) and trophic niches (Figure 6B, but also see fecal content from Santamaría et al. 2019), suggesting both a potential host genotype and diet effect.

A large majority of the ASV and taxa seasonal markers were repeatedly observed along most sampling dates (although not in 50% of the population each date) (Figure 8B), indicating that changes were essentially quantitative (i.e., associated with variation in relative abundance, although not absolute counts, of microbial stable associates), excluding a major compositional turnover each season. ASVs seasonal makers were mainly associated to the families Ruminococcaceae and Lachnospiraceae, which have been repeatedly involved in seasonal microbial reconfiguration in mammals (Baniel et al., 2021; Maurice et al., 2015) suggesting they might represent the more plastic component of the microbiota in response to temporal dietary/physiological shifts. Among taxa, both islets showed a clear spring enrichment of the family Odoribacteriaceae and the associated genus *Odoribacter* (order Bacteroidales), a strictly anaerobic and butyrate-producing bacterium. This genus is a stable and abundant associate of lizards (Kohl et al., 2017; Zhang et al., 2018)and more general of vertebrates (Dutton et al., 2021) and it has been identified as critical metabolic players in the human gut (Hiippala et al., 2020), whereas its functional role in the Lilford’s wall lizard gut remains to be investigated.

Overall, these seasonal microbial markers might represent important players for the gut microbiome plasticity. Whether this plasticity confers metabolic flexibility to their host, as recently observed for mammals (Guo et al., 2021; Hicks et al., 2018; Maurice et al., 2015), remains to be investigated. Unlike mammals, reptiles are ectothermic and their metabolic flexibility and putative physiological and energetic dependence on gut microbes might be under different regulatory processes (Moeller et al., 2020), an intriguing avenue that is definitely worth future research.

## Conclusions

This study provides a first in-depth exploration of the trends governing gut microbial dynamics between and within populations of *P. lilfordi*. By taking advantage of individual-based microbiota data and performing comparative analyses of three sister populations found in near islets, we showed that microbial diversity among populations is primarily driven by small qualitative changes, that is by the presence of few islet-specifics bacterial ASVs, with neglectable variation in taxa membership. This suggests that the host genotype largely overrides the effect of geographic barriers and local exposure to different environmental pools in terms of major microbial profiles, while the environment progressively drives the diversification of symbiotic communities at strain level along that of their host populations. It remains unclear to what extent these small differences in community composition are adaptive, for instance in response to population adjustment to the trophic niche, or shaped by ecological drift, including a putative differential retention of ancestral taxa from the common ancestral population. A crucial aspect that still needs to be investigated is to which extent these gut bacteria are transmitted among *Podarcis* individuals and through the host generations.

On the other hand, microbiota diversity within populations is marked by seasonality, resulting in microbial cyclic fluctuations, consistently along the two sampled years and for both populations studied and suggesting a deterministic pattern in microbiome temporal changes over external stimuli (diet/season). These data clearly need to be corroborated by resampling of the same individuals across seasons. Collection of cloacal swabs instead of feces could help in sample size optimization (currently limited by lizard rare defecation) providing a means for longitudinal studies of gut microbiota in lizard natural populations.

Overall, the observed lack of an important turnover (i.e., no significant replacement of major taxa) in the main microbial community composition across allopatric populations of *P. lilfordi* supports a strong resilience of these gut microbial communities along the short-term evolutionary times of their host diversification, implying strength and specificity of this symbiosis. Furthermore, replicated quantitative changes in microbial reconfiguration along seasons indicated that the microbiota is, to some extent, a plastic trait that can potentially adjust to the host temporal/metabolic changes in a rather predictive manner. The challenge is now to understand the impact of microbial community signature and its plasticity on the *P. lilfordi fitness* and ecological adaptation to these small islets, as well as to evaluate the great potential of integrative holobiont studies in monitoring this endangered wild species.

## Acknowledgements

We thank I. Hendriks for aid in sample collection. This study was supported by the Agencia Estatal de Investigación (AEI) and the Fondo Europeo de Desarrollo Regional (FEDER) (CGL2017-82986-C2-2-P) to L.B. and by the project PRD2018/25 (CAIB - Government of the Balearic Islands) to G.T.

## Authors’ contributions

L.B. conceived the ideas, L.B., G.T., J.M.I. and A.R.V. performed the sampling and collected metadata; L.B. and J.L.R. designed methodology and analysed the data; L.B. led the writing of the manuscript. All authors contributed critically to the drafts and gave final approval for publication.

## Data Availability Statement

Raw 16S rRNA Miseq Data is available at the Bioproject PRJNA764850. Original ASV abundance matrix per sample and corresponding taxonomic classification is available on Mendeley Data (doi: 10.17632/bc5nxsxgxd.1).

## Supporting Information

**Figure S1:** Rarefaction curves (step = 1000 counts) summarising sequencing effort per sample/specimen. The dashed line shows the sample with minimum sequence coverage (260003 sequence counts).

**Figure S2:** Microbiota taxonomic composition (phylum and family) at specimen level. Legends list only the top ten most abundant taxa. The remaining were included in “Others”.

**Figure S3:** Microbiota centroid distances according to unweighted Unifrac, estimated on different core subsets, and host genetic distances according to *Fst* values estimated from published microsatellites data (Rotger et al. 2021).

**Figure S4:** Venn diagrams showing shared ASVs across the four sampling dates (i.e. persistent ASVs). For each date, we considered only ASVs found in at least 50% of the specimens.

**Figure S5:** PCoA of microbiota Bray-Curtis distances according to Date, estimated on different core subsets (50, 70, 80 and 90%). The rectangular box represents the centroid per date.

**Figure S6:** Variation in relative abundance along “dates” of taxa that were significantly enriched in either spring or autumn according to LEfSe and/or indval analyses. Fluctuations in relative abundance between dates for all taxonomic levels (Species to Phylum) were calculated as mean relative abundances on reads aggregated by levels (i.e., after adding all all reads corresponding to a particular taxon).

## Appendix S1

**Table S1:** Sample metadata.

**Table S2:** Primers used for amplification of the region V3-V4 of 16S rRNA.

**Table S3:** Abundance matrix of sequence counts found in the controls (PCR negative controls and mock communities) and corresponding taxonomic classification.

**Table S4:** List of ASVs that significantly discriminated among islets based on both seasonal datasets and according to both “indval” (in the *labdsv* R package) and LEfSe analyses.

**Table S5:** Stable isotopes estimated on 71 specimens collected during spring 2016.

**Table S6:** List of persistent ASVs found in NG.

**Table S7:** List of persistent ASVs found in NM.

**Table S8:** List of ASVs that significantly discriminated between seasons within single islets (NG and NM) according to both indval and LEfSe analyses.

## References

Alberdi, A., Aizpurua, O., Bohmann, K., Zepeda-Mendoza, M. L., & Gilbert, M. T. P. (2016). Do Vertebrate Gut Metagenomes Confer Rapid Ecological Adaptation? In Trends in Ecology and Evolution (Vol. 31, Issue 9, pp. 689–699). Elsevier Ltd. https://doi.org/10.1016/j.tree.2016.06.008

Alberdi, A., Andersen, S. B., Limborg, M. T., Dunn, R. R., & Gilbert, M. T. P. (2021). Disentangling host–microbiota complexity through hologenomics. Nature Reviews Genetics. https://doi.org/10.1038/s41576-021-00421-0

Alcover, J. A. (2000). Vertebrate Evolution and Extinction on Western and Central Mediterranean Islands. TROPICS, 10(1), 103–123.

Amir, A., McDonald, D., Navas-Molina, J. A., Kopylova, E., Morton, J. T., Zech Xu, Z., Kightley, E. P., Thompson, L. R., Hyde, E. R., Gonzalez, A., & Knight, R. (2017). Deblur Rapidly Resolves Single-Nucleotide Community Sequence Patterns. MSystems, 2(2). https://doi.org/10.1128/msystems.00191-16

Baldo, L., Riera, J. L., Mitsi, K., & Pretus, J. L. (2018). Processes shaping gut microbiota diversity in allopatric populations of the endemic lizard Podarcis lilfordi from Menorcan islets (Balearic Islands). FEMS Microbiology Ecology, 94(2), 1–14. https://doi.org/10.1093/femsec/fix186

Baniel, A., Amato, K. R., Beehner, J. C., Bergman, T. J., Mercer, A., Perlman, R. F., Petrullo, L., Reitsema, L., Sams, S., Lu, A., & Snyder-Mackler, N. (2021). Seasonal shifts in the gut microbiome indicate plastic responses to diet in wild geladas. Microbiome, 9(1). https://doi.org/10.1186/S40168-020-00977-9

Bittinger, K. (2020). usedist: Distance Matrix Utilities. R package version 0.4.0. Https://CRAN.R-Project.Org/Package=usedist.

Bittkau, C., & Comes, H. P. (2005). Evolutionary processes in a continental island system: Molecular phylogeography of the Aegean Nigella arvensis alliance (Ranunculaceae) inferred from chloroplast DNA. Molecular Ecology, 14(13), 4065–4083. https://doi.org/10.1111/j.1365-294X.2005.02725.x

Brown, R. P., Terrasa, B., Pérez-Mellado, V., Castro, J. A., Hoskisson, P. A., Picornell, A., & Ramon, M. M. (2008). Bayesian estimation of post-Messinian divergence times in Balearic Island lizards. Molecular Phylogenetics and Evolution, 48(1), 350–358. https://doi.org/10.1016/j.ympev.2008.04.013

Buades, J. M., Rodríguez, V., Terrasa, B., Pérez-Mellado, V., Brown, R. P., Castro, J. A., Picornell, A., & Ramon, M. M. (2013). Variability of the mc1r Gene in Melanic and Non-Melanic Podarcis lilfordi and Podarcis pityusensis from the Balearic Archipelago. PLoS ONE, 8(1). https://doi.org/10.1371/journal.pone.0053088

Capella-Gutiérrez, S., Silla-Martínez, J. M., & Gabaldón, T. (2009). trimAl: A tool for automated alignment trimming in large-scale phylogenetic analyses. Bioinformatics, 25(15), 1972–1973. https://doi.org/10.1093/bioinformatics/btp348

Caporaso, J. G., Kuczynski, J., Stombaugh, J., Bittinger, K., Bushman, F. D., Costello, E. K., Fierer, N., Pẽa, A. G., Goodrich, J. K., Gordon, J. I., Huttley, G. A., Kelley, S. T., Knights, D., Koenig, J. E., Ley, R. E., Lozupone, C. A., McDonald, D., Muegge, B. D., Pirrung, M., … Knight, R. (2010). QIIME allows analysis of high-throughput community sequencing data. In Nature Methods (Vol. 7, Issue 5, pp. 335–336). https://doi.org/10.1038/nmeth.f.303

Castilla, A., & Bauwens, D. (2000). Reproductive Characteristics of the Island Lacertid Lizard Podarcis lilfordi. Journal of Herpetology, 34(3), 390–396.

Colston, T. J. (2017). Gut microbiome transmission in lizards. In Molecular ecology (Vol. 26, Issue 4, pp. 972–974). NLM (Medline). https://doi.org/10.1111/mec.13987

Cooper, W. E., Dimopoulos, I., & Pafilis, P. (2015). Sex, age, and population density affect aggressive behaviors in Island lizards promoting cannibalism. Ethology, 121(3), 260–269. https://doi.org/10.1111/eth.12335

Costello, E. K., Gordon, J. I., Secor, S. M., & Knight, R. (2010). Postprandial remodeling of the gut microbiota in Burmese pythons. ISME Journal, 4(11), 1375–1385. https://doi.org/10.1038/ismej.2010.71

Davison, J., Moora, M., Öpik, M., Ainsaar, L., Ducousso, M., Hiiesalu, I., Jairus, T., Johnson, N., Jourand, P., Kalamees, R., Koorem, K., Meyer, J. Y., Püssa, K., Reier, Ü., Pärtel, M., Semchenko, M., Traveset, A., Vasar, M., & Zobel, M. (2018). Microbial island biogeography: isolation shapes the life history characteristics but not diversity of root-symbiotic fungal communities. ISME Journal, 12(9), 2211–2224. https://doi.org/10.1038/s41396-018-0196-8

Dutton, C. L., Subalusky, A. L., Sanchez, A., Estrela, S., Lu, N., Hamilton, S. K., Njoroge, L., Rosi, E. J., & Post, D. M. (2021). The meta-gut: community coalescence of animal gut and environmental microbiomes. Scientific Reports |, 11(23117). https://doi.org/10.1038/s41598-021-02349-1

Gomez, A., Sharma, A. K., Mallott, E. K., Petrzelkova, K. J., Robinson, C. A. J., Yeoman, C. J., Carbonero, F., Pafco, B., Rothman, J. M., Ulanov, A., Vlckova, K., Amato, K. R., Schnorr, S. L., Dominy, N. J., Modry, D., Todd, A., Torralba, M., Nelson, K. E., Burns, M. B., … Leigh, S. R. (2019). Plasticity in the Human Gut Microbiome Defies Evolutionary Constraints. 4, 271–290. https://doi.org/10.1128/mSphere

Gophna, U., Konikoff, T., & Nielsen, H. B. (2017). Oscillospira and related bacteria – From metagenomic species to metabolic features. Environmental Microbiology, 19(3), 835–841. https://doi.org/10.1111/1462-2920.13658

Grieneisen, L., Dasari, M., Gould, T. J., Björk, J. R., Grenier, J. C., Yotova, V., Jansen, D., Gottel, N., Gordon, J. B., Learn, N. H., Gesquiere, L. R., Wango, T. L., Mututua, R. S., Warutere, J. K., Siodi, L., Gilbert, J. A., Barreiro, L. B., Alberts, S. C., Tung, J., … Blekhman, R. (2021). Gut microbiome heritability is nearly universal but environmentally contingent. Science, 373(6551), 181–186. https://doi.org/10.1126/science.aba5483

Guo, N., Wu, Q., Shi, F., Niu, J., Zhang, T., Degen, A. A., Fang, Q., Ding, L., Shang, Z., Zhang, Z., & Long, R. (2021). Seasonal dynamics of diet–gut microbiota interaction in adaptation of yaks to life at high altitude. Npj Biofilms and Microbiomes, 7(1). https://doi.org/10.1038/s41522-021-00207-6

Henry, L. P., Bruijning, M., Forsberg, S. K. G., & Ayroles, J. F. (2021). The microbiome extends host evolutionary potential. In Nature Communications (Vol. 12, Issue 1). Nature Research. https://doi.org/10.1038/s41467-021-25315-x

Hicks, A. L., Jo Lee, K., Couto-Rodriguez, M., Patel, J., Sinha, R., Guo, C., Olson, S. H., Seimon, A., Seimon, T. A., Ondzie, A. U., Karesh, W. B., Reed, P., Cameron, K. N., Lipki, W. I., & Brent, L. W. (2018). Gut microbiomes of wild great apesfluctuateseasonally in response to diet. Nature Communications, 9(1786).

Hiippala, K., Barreto, G., Burrello, C., Diaz-Basabe, A., Suutarinen, M., Kainulainen, V., Bowers, J. R., Lemmer, D., Engelthaler, D. M., Eklund, K. K., Facciotti, F., & Satokari, R. (2020). Novel Odoribacter splanchnicus Strain and Its Outer Membrane Vesicles Exert Immunoregulatory Effects in vitro. Frontiers in Microbiology, 11. https://doi.org/10.3389/fmicb.2020.575455

Huus, K. E., & Ley, R. E. (2021). Blowing Hot and Cold: Body Temperature and the Microbiome. MSystems.

Jiang, H. Y., Ma, J. E., Li, J., Zhang, X. J., Li, L. M., He, N., Liu, H. Y., Luo, S. Y., Wu, Z. J., Han, R. C., & Chen, J. P. (2017). Diets alter the gut microbiome of crocodile lizards. Frontiers in Microbiology, 8(OCT). https://doi.org/10.3389/FMICB.2017.02073

Kohl, K. D., Brun, A., Magallanes, M., Brinkerhoff, J., Laspiur, A., Acosta, J. C., Bordenstein, S. R., & Caviedes-Vidal, E. (2016). Physiological and microbial adjustments to diet quality permit facultative herbivory in an omnivorous lizard. Journal of Experimental Biology, 219(12), 1903–1912. https://doi.org/10.1242/jeb.138370

Kohl, K. D., Brun, A., Magallanes, M., Brinkerhoff, J., Laspiur, A., Acosta, J. C., Caviedes-Vidal, E., & Bordenstein, S. R. (2017). Gut microbial ecology of lizards: insights into diversity in the wild, effects of captivity, variation across gut regions and transmission. Molecular Ecology, 26(4), 1175–1189. https://doi.org/10.1111/mec.13921

Kohl, K. D., Weiss, R. B., Cox, J., Dale, C., & Dearing, M. D. (2014). Gut microbes of mammalian herbivores facilitate intake of plant toxins. Ecology Letters, 17(10), 1238–1246. https://doi.org/10.1111/ele.12329

Lankau, E. W., Hong, P. Y., & MacKie, R. I. (2012). Ecological drift and local exposures drive enteric bacterial community differences within species of Galápagos iguanas. Molecular Ecology, 21(7), 1779–1788. https://doi.org/10.1111/j.1365-294X.2012.05502.x

Leitão-Gonç alves, R., Carvalho-Santos, Z., Patrícia Francisco, A., Tondolo Fioreze, G., Anjos, M., lia Baltazar, C., Paula Elias, A., Itskov, P. M., W Piper, M. D., & Ribeiro, C. (2017). Commensal bacteria and essential amino acids control food choice behavior and reproduction. https://doi.org/10.1371/journal.pbio.2000862

Lenth, R. (2022). emmeans: Estimated Marginal Means, aka Least-Squares Means. R Package Version 1.7.2. Https://CRAN.R-Project.Org/Package=emmeans.

Levin, D., Raab, N., Pinto, Y., Rothschild, D., Zanir, G., Godneva, A., Mellul, N., Futorian, D., Gal, D., Leviatan, S., Zeevi, D., Bachelet, I., & Segal, E. (2021). Diversity and functional landscapes in the microbiota of animals in the wild. Science, 372(6539). https://doi.org/10.1126/SCIENCE.ABB5352

Li, G., Li, J., Kohl, K. D., Yin, B., Wei, W., Wan, X., Zhu, B., & Zhang, Z. (2019). Dietary shifts influenced by livestock grazing shape the gut microbiota composition and co-occurrence networks in a local rodent species. Journal of Animal Ecology, 88(2), 302–314. https://doi.org/10.1111/1365-2656.12920

Lim, S. J., & Bordenstein, S. R. (2020). An introduction to phylosymbiosis. In Proceedings of the Royal Society B: Biological Sciences (Vol. 287, Issue 1922). Royal Society Publishing. https://doi.org/10.1098/rspb.2019.2900

Lindsay, E. C., Metcalfe, N. B., & Llewellyn, M. S. (2020). The potential role of the gut microbiota in shaping host energetics and metabolic rate. In Journal of Animal Ecology (Vol. 89, Issue 11, pp. 2415–2426). Blackwell Publishing Ltd. https://doi.org/10.1111/1365-2656.13327

MacArthur, R. H., & Wilson, E. O. (1967). The theory of island biogeography / Robert H. MacArthur and Edward O. Wilson. Princeton University Press Princeton, N.J.

Mallott, E. K., & Amato, K. R. (2021). Host specificity of the gut microbiome. In Nature Reviews Microbiology (Vol. 19, Issue 10, pp. 639–653). Nature Research. https://doi.org/10.1038/s41579-021-00562-3

Maurice, C. F., Knowles, S. C., Ladau, J., Pollard, K. S., Fenton, A., Pedersen, A. B., & Turnbaugh, P. J. (2015). Marked seasonal variation in the wild mouse gut microbiota. The ISME Journal, 9, 2423–2434. https://doi.org/10.1038/ismej.2015.53

McDonald, D., Price, M. N., Goodrich, J., Nawrocki, E. P., Desantis, T. Z., Probst, A., Andersen, G. L., Knight, R., & Hugenholtz, P. (2012). An improved Greengenes taxonomy with explicit ranks for ecological and evolutionary analyses of bacteria and archaea. ISME Journal, 6(3), 610–618. https://doi.org/10.1038/ismej.2011.139

McMurdie, P. J., & Holmes, S. (2013). phyloseq: An R Package for Reproducible Interactive Analysis and Graphics of Microbiome Census Data. PLOS ONE, 8(4), e61217–. https://doi.org/10.1371/journal.pone.0061217

Méheust, R., Castelle, C. J., Matheus Carnevali, P. B., Farag, I. F., He, C., Chen, L.-X., Amano, Y., Hug, L. A., & Banfield, J. F. (2020). Groundwater Elusimicrobia are metabolically diverse compared to gut microbiome Elusimicrobia and some have a novel nitrogenase paralog. The ISME Journal, 14, 2907–2922. https://doi.org/10.1038/s41396-020-0716-1

Michel, A. J., Ward, L. M., Goffredi, S. K., Dawson, K. S., Baldassarre, D. T., Brenner, A., Gotanda, K. M., McCormack, J. E., Mullin, S. W., O’Neill, A., Tender, G. S., Uy, J. A. C., Yu, K., Orphan, V. J., & Chaves, J. A. (2018). The gut of the finch: Uniqueness of the gut microbiome of the Galápagos vampire finch 06 Biological Sciences 0602 Ecology 05 Environmental Sciences 0502 Environmental Science and Management. Microbiome, 6(1). https://doi.org/10.1186/s40168-018-0555-8

Miller, E. T., Svanbäck, R., & Bohannan, B. J. M. (2018). Microbiomes as Metacommunities: Understanding Host-Associated Microbes through Metacommunity Ecology. In Trends in Ecology and Evolution. https://doi.org/10.1016/j.tree.2018.09.002

Moeller, A. H., Ivey, K., Cornwall, M. B., Herr, K., Rede, J., Taylor, E. N., & Gunderson, A. R. (2020). The lizard gut microbiome changes with temperature and is associated with heat tolerance. Applied and Environmental Microbiology, 86(17). https://doi.org/10.1128/AEM.01181-20

Montoya-Ciriaco, N., Gómez-Acata, S., Muñoz-Arenas, L. C., Dendooven, L., Estrada-Torres, A., Díaz De La Vega-Pérez, A. H., & Navarro-Noya, Y. E. (2020). Dietary effects on gut microbiota of the mesquite lizard Sceloporus grammicus (Wiegmann, 1828) across different altitudes. Microbiome, 8(1). https://doi.org/10.1186/s40168-020-0783-6

Moya, Ó., Pep-Luis, M., Sergio, M., José-Manuel, I., Andreu, R., Antonio, R., & Giacomo, T. (2015). APHIS: A new software for photo-matching in ecological studies. Ecological Informatics, 27, 64–70. https://doi.org/10.1016/j.ecoinf.2015.03.003

Nearing, J. T., Douglas, G. M., Hayes, M. G., Macdonald, J., Desai, D. K., Allward, N., Jones, C. M. A., Wright, R. J., Dhanani, A. S., Comeau, A. M., & Langille, M. G. I. (2022). Microbiome differential abundance methods produce different results across 38 datasets. Nature Communications, 13, 342. https://doi.org/10.1038/s41467-022-28034-z

Oksanen, J., Blanchet, F. G., Friendly, M., Kindt, R., Legendre, P., Mcglinn, D., Minchin, P. R., O’hara, R. B., Simpson, G. L., Solymos, P., Henry, M., Stevens, H., Szoecs, E., & Maintainer, H. W. (2020). Package “vegan” Title Community Ecology Package Version 2.5-7.

Patnode, M. L., Beller, Z. W., Han, N. D., Cheng, J., Peters, S. L., Terrapon, N., Henrissat, B., le Gall, S., Saulnier, L., Hayashi, D. K., Meynier, A., Vinoy, S., Giannone, R. J., Hettich, R. L., & Gordon, J. I. (2019). Interspecies Competition Impacts Targeted Manipulation of Human Gut Bacteria by Fiber-Derived Glycans. Cell, 179(1), 59–73.e13. https://doi.org/10.1016/j.cell.2019.08.011

Pérez-Cembranos, A., León, A., & Pérez-Mellado, V. (2016). Omnivory of an Insular Lizard: Sources of Variation in the Diet of Podarcis lilfordi (Squamata, Lacertidae). https://doi.org/10.1371/journal.pone.0148947

Pérez-Cembranos, A., Pérez-Mellado, V., Alemany, I., Bassitta, M., Terrasa, B., Picornell, A., Castro, J. A., Brown, R. P., & Ramon, C. (2020). Morphological and genetic diversity of the Balearic lizard, Podarcis lilfordi (Günther, 1874): Is it relevant to its conservation? Diversity and Distributions, 26(9), 1122–1141. https://doi.org/10.1111/ddi.13107

Pérez-Mellado, V., García-Díez, T., Hernández-Estévez, J. A., & Tavecchia, & G. (2015). Behavioural processes, ephemeral resources and spring population dynamics of an insular lizard, Podarcis lilfordi (Squamata: Lacertidae). https://doi.org/10.1080/11250003.2015.1093035

Price, M. N., Dehal, P. S., & Arkin, A. P. (2009). Fasttree: Computing large minimum evolution trees with profiles instead of a distance matrix. Molecular Biology and Evolution, 26(7), 1641–1650. https://doi.org/10.1093/molbev/msp077

Raulo, A., Allen, B. E., Troitsky, T., Husby, A., Firth, J. A., Coulson, T., & Knowles, S. C. L. (2021). Social networks strongly predict the gut microbiota of wild mice. ISME Journal. https://doi.org/10.1038/s41396-021-00949-3

Ren, T., Kahrl, A. F., Wu, M., & Cox, R. M. (2016). Does adaptive radiation of a host lineage promote ecological diversity of its bacterial communities? A test using gut microbiota of Anolis lizards. Molecular Ecology, 25(19), 4793–4804. https://doi.org/10.1111/mec.13796

Roberts, D. W. (2019). Ordination and multivariate analysis for ecology. R Package Version 1.5-0. Http://CRAN.R-Project.Org/Package=labdsv.

Robinson, C. D., Bohannan, B. J. M., & Britton, R. A. (2019). Scales of persistence: transmission and the microbiome Graphical Abstract HHS Public Access. Curr Opin Microbiol, 50, 42–49. https://doi.org/10.1016/j.mib.2019.09.009

Rojas, C. A., Ramírez-Barahona, S., Holekamp, K. E., & Theis, K. R. (2021). Host phylogeny and host ecology structure the mammalian gut microbiota at different taxonomic scales. Animal Microbiome, 3(1). https://doi.org/10.1186/s42523-021-00094-4

Rotger, A., Igual, J. M., Genovart, M., Rodríguez, V., Ramon, C., Pérez-Mellado, V., Bibiloni, G., Rita, J., & Tavecchia, G. (2021). Contrasting Adult Body-Size in Sister Populations of the Balearic Lizard, Podarcis lilfordi (Günther 1874) Suggests Anthropogenic Selective Pressures. Herpetological Monographs, 35(1). https://doi.org/10.1655/HERPMONOGRAPHS-D-19-00005

Rotger, A., Igual, J. M., Smith, J. J., & Tavecchia, G. (2016). Relative role of population density and climatic factors in shaping the body growth rate of lilford’s wall lizard (Podarcis lilfordi). Canadian Journal of Zoology, 94(3), 207–215. https://doi.org/10.1139/cjz-2015-0188

Rotger, A., Igual, J. M., & Tavecchia, G. (2020). Contrasting size-dependent life history strategies of an insular lizard. Current Zoology, 66(6), 625–633. https://doi.org/10.1093/CZ/ZOAA019

Rowe, M., Veerus, L., Trosvik, P., Buckling, A., & Pizzari, T. (2020). The Reproductive Microbiome: An Emerging Driver of Sexual Selection, Sexual Conflict, Mating Systems, and Reproductive Isolation. In Trends in Ecology and Evolution (Vol. 35, Issue 3, pp. 220–234). Elsevier Ltd. https://doi.org/10.1016/j.tree.2019.11.004

Salvador, A. (2009). ENCICLOPEDIA VIRTUAL DE LOS VERTEBRADOS ESPAÑOLES Sociedad de Amigos del MNCN-MNCN-CSIC Lagartija balear-Podarcis lilfordi (Günther, 1874). http://www.vertebradosibericos.org/

Santamaría, S., Enoksen, C. A., Olesen, J. M., Tavecchia, G., Rotger, A., Igual, J. M., & Traveset, A. (2019). Diet composition of the lizard Podarcis lilfordi (Lacertidae) on 2 small islands: an individual-resource network approach. Current Zoology, May, 1–11. https://doi.org/10.1093/cz/zoz028

Segata, N., Izard, J., Waldron, L., Gevers, D., Miropolsky, L., Garrett, W. S., & Huttenhower, C. (2011). Metagenomic biomarker discovery and explanation. https://doi.org/10.1186/gb-2011-12-6-r60

Shapira, M. (2016). Gut Microbiotas and Host Evolution: Scaling Up Symbiosis. In Trends in Ecology and Evolution. https://doi.org/10.1016/j.tree.2016.03.006

Smits, S. A., Leach, J., Sonnenburg, E. D., Gonzalez, C. G., Lichtman, J. S., Reid, G., Knight, R., Manjurano, A., Changalucha, J., Elias, J. E., Gloria Dominguez-Bello, M., & Sonnenburg, J. L. (2017). Seasonal cycling in the gut microbiome of the Hadza hunter-gatherers of Tanzania. Science, 357, 802–806. https://www.science.org

Sommer, F., Stå, M., Ilkayeva, O., Newgard, C. B., Frö, O., Bä Ckhed Correspondence, F., Arnemo, J. M., Kindberg, J., Josefsson, J., & Bä Ckhed, F. (2016). The Gut Microbiota Modulates Energy Metabolism in the Hibernating Brown Bear Ursus arctos. Cell Reports, 14. https://doi.org/10.1016/j.celrep.2016.01.026

Song, J., Amir, A., Metcalf, J. L., Amato, K. R., Xu, Z. Z., Humphrey, G., Knight, R., & Dearing, E. M. D. (2016). Preservation Methods Differ in Fecal Microbiome Stability, Affecting Suitability for Field Studies Downloaded from. https://doi.org/10.1128/mSystems.00021-16

Terrasa, B., Pérez-Mellado, V., Brown, R. P., Picornell, A., Castro, J. A., & Ramon, M. M. (2009). Foundations for conservation of intraspecific genetic diversity revealed by analysis of phylogeographical structure in the endangered endemic lizard Podarcis lilfordi. Diversity and Distributions, 15(2), 207–221. https://doi.org/10.1111/j.1472-4642.2008.00520.x

Thaiss, C. A., Zmora, N., Levy, M., & Elinav, E. (2016). The microbiome and innate immunity. In Nature (Vol. 535, Issue 7610, pp. 65–74). Nature Publishing Group. https://doi.org/10.1038/nature18847

Tung, J., Barreiro, L. B., Burns, M. B., Grenier, J. C., Lynch, J., Grieneisen, L. E., Altmann, J., Alberts, S. C., Blekhman, R., & Archie, E. A. (2015). Social networks predict gut microbiome composition in wild baboons. ELife, 2015(4), 1–18. https://doi.org/10.7554/eLife.05224

van Damme, R. (1999). Evolution of Herbivory in Lacertid Lizards: Effects of Insularity and Body Size. Journal of Herpetology, 33(4), 663–674.

Velo-Antó, G., Zamudio, K. R., & Cordero-Rivera, A. (2012). Genetic drift and rapid evolution of viviparity in insular fire salamanders (Salamandra salamandra). Heredity, 108, 410–418. https://doi.org/10.1038/hdy.2011.91

Warne, R. W., Kirschman, | Lucas, & Zeglin, L. (2019). Manipulation of gut microbiota during critical developmental windows affects host physiological performance and disease susceptibility across ontogeny. J Anim Ecol, 88. https://doi.org/10.1111/1365-2656.12973

Youngblut, N. D., Reischer, G. H., Walters, W., Schuster, N., Walzer, C., Stalder, G., Ley, R. E., & Farnleitner, A. H. (2019). Host diet and evolutionary history explain different aspects of gut microbiome diversity among vertebrate clades. Nature Communications, 10(1). https://doi.org/10.1038/s41467-019-10191-3

Zhang, W., Li, N., Tang, X., Liu, N., & Zha, W. (2018). Changes in intestinal microbiota across an altitudinal gradient in the lizard Phrynocephalus vlangalii | Enhanced Reader. Ecology and Evolution, 8, 4695–4703. https://doi.org/10.1002/ece3.4029

