## Appendix S1 for "Insular holobionts: persistence and seasonal plasticity of the Balearic wall lizard (*Podarcis lilfordi*) gut microbiota"

**Table S1:** Sample metadata.

| Specimen_ID | Islet | Sampling_date | Season_Year | Life_Stage | Sex | Weight _gr | SVL_mm | LAT | LON |
|---|---|---|---|---|---|---|---|---|---|
| MO11 | Na Moltona | 18.10.17 | Autumn-17 | Adult | F | 6,4 | 65 | 501017,4694 | 4350644,669 |
| MO112 | Na Moltona | 20.10.17 | Autumn-17 | Adult | F | 6,5 | 66 | 501053,8679 | 4350642,367 |
| MO165 | Na Moltona | 20.10.17 | Autumn-17 | Adult | M | 7,6 | 65 | 500977 | 4350590 |
| MO183 | Na Moltona | 24.10.17 | Autumn-17 | Adult | F | 5,6 | 62 | 501017,4694 | 4350644,669 |
| MO185 | Na Moltona | 24.10.17 | Autumn-17 | Adult | M | 8,6 | 71 | 501017,7571 | 4350639,777 |
| MO2 | Na Moltona | 18.10.17 | Autumn-17 | Adult | M | 12,3 | 75 | 501053,8679 | 4350642,367 |
| MO200 | Na Moltona | 24.10.17 | Autumn-17 | Adult | F | 5,8 | 64 | 501043,0779 | 4350620,93 |
| MO209 | Na Moltona | 24.10.17 | Autumn-17 | Adult | M | 10,3 | 73 | 500984 | 4350580 |
| MO20A | Na Moltona | 18.10.17 | Autumn-17 | Adult | M | 7,7 | 71 | 501048,8326 | 4350631,72 |
| MO234 | Na Moltona | 24.10.17 | Autumn-17 | Adult | F | 5,1 | 63 | 501042,0708 | 4350638,051 |
| MO256 | Na Moltona | 24.10.17 | Autumn-17 | Adult | F | 5,4 | 62 | 501027 | 4350580 |
| MO3 | Na Moltona | 18.10.17 | Autumn-17 | Adult | F | 5,1 | 61 | 501028,4033 | 4350643,23 |
| MO40 | Na Moltona | 18.10.17 | Autumn-17 | Adult | F | 6,2 | 65 | 501035,309 | 4350613,449 |
| MO42A | Na Moltona | 18.10.17 | Autumn-17 | Adult | M | 6,7 | 63 | 501018 | 4350590 |
| MO73 | Na Moltona | 18.10.17 | Autumn-17 | Adult | F | 6,8 | 67 | 501057,3208 | 4350633,303 |
| M102 | Na Moltona | 12.04.17 | Spring-17 | Adult | F | 7,1 | 68 | 501016 | 4350580 |
| M104 | Na Moltona | 12.04.17 | Spring-17 | Adult | F | 8,7 | 69 | 501032 | 4350580 |
| M107 | Na Moltona | 12.04.17 | Spring-17 | Adult | M | 10,9 | 78 | 501017,4694 | 4350644,669 |
| M11 | Na Moltona | 10.04.17 | Spring-17 | Adult | F | 8 | 69 | 500999 | 4350590 |
| M19 | Na Moltona | 10.04.17 | Spring-17 | Adult | M | 9,8 | 70 | 501028,4033 | 4350643,23 |
| M21A | Na Moltona | 10.04.17 | Spring-17 | Adult | F | 6 | 61 | 501021,6415 | 4350635,749 |
| M40 | Na Moltona | 10.04.17 | Spring-17 | Adult | F | 5,6 | 64 | 500991 | 4350590 |
| M48 | Na Moltona | 11.04.17 | Spring-17 | Adult | F | 6,5 | 65 | 501060,1981 | 4350629,562 |
| M68 | Na Moltona | 11.04.17 | Spring-17 | Subadult | F | 4,4 | 53 | 501006 | 4350580 |
| M74 | Na Moltona | 11.04.17 | Spring-17 | Adult | F | 7,8 | 65 | 501023,0802 | 4350628,843 |
| M8 | Na Moltona | 10.04.17 | Spring-17 | Adult | F | 5,2 | 62 | 501035,309 | 4350613,449 |
| M81 | Na Moltona | 11.04.17 | Spring-17 | Adult | F | 6,2 | 63 | 500984 | 4350580 |
| M96 | Na Moltona | 12.04.17 | Spring-17 | Adult | F | 5,7 | 62 | 501012,2901 | 4350650,855 |
| MO10 | Na Moltona | 25.10.18 | Autumn-18 | Adult | F | 4,9 | 60 | 501042,0708 | 4350638,051 |
| MO14 | Na Moltona | 25.10.18 | Autumn-18 | Adult | F | 7,4 | 68 | 501043,0779 | 4350620,93 |
| MO17 | Na Moltona | 25.10.18 | Autumn-18 | Adult | F | 7,4 | 68 | 500984 | 4350580 |
| MO20B | Na Moltona | 25.10.18 | Autumn-18 | Subadult | F | 3,9 | 50 | 501023 | 4350580 |
| MO27 | Na Moltona | 25.10.18 | Autumn-18 | Adult | M | 13,2 | 77 | 501021,6415 | 4350635,749 |
| MO31 | Na Moltona | 25.10.18 | Autumn-18 | Adult | F | 6,6 | 65 | 501042,5024 | 4350614,888 |
| MO32 | Na Moltona | 25.10.18 | Autumn-18 | Adult | M | 10,2 | 71 | 501028,8349 | 4350616,614 |
| MO33 | Na Moltona | 25.10.18 | Autumn-18 | Adult | F | 6,6 | 66 | 501005 | 4350590 |
| MO36 | Na Moltona | 25.10.18 | Autumn-18 | Adult | M | 7,8 | 67 | 501005 | 4350570 |
| MO42B | Na Moltona | 25.10.18 | Autumn-18 | Adult | M | 7,8 | 67 | 501052,1415 | 4350616,039 |
| MO44 | Na Moltona | 25.10.18 | Autumn-18 | Adult | F | 6,3 | 66 | 501042,5024 | 4350614,888 |
| MO60 | Na Moltona | 26.10.18 | Autumn-18 | Adult | F | 7,4 | 63 | 501053,8679 | 4350642,367 |
| MO88 | Na Moltona | 26.10.18 | Autumn-18 | Adult | F | 6,8 | 68 | 500995 | 4350570 |
| MO91 | Na Moltona | 26.10.18 | Autumn-18 | Adult | F | 5,6 | 64 | 501023 | 4350580 |
| M149 | Na Moltona | 02.05.18 | Spring-18 | Juvenile | M | 2,9 | 47 | 501032 | 4350580 |
| M17 | Na Moltona | 20.04.18 | Spring-18 | Juvenile | M | 2,8 | 52 | 501035,309 | 4350613,449 |
| M21B | Na Moltona | 20.04.18 | Spring-18 | Adult | M | 10,7 | 74 | 500984 | 4350580 |
| M30 | Na Moltona | 20.04.18 | Spring-18 | Juvenile | M | 2,1 | 46 | 501041,9269 | 4350650,999 |
| M33 | Na Moltona | 20.04.18 | Spring-18 | Adult | F | 7,5 | 67 | 501023,0802 | 4350628,843 |
| M42 | Na Moltona | 20.04.18 | Spring-18 | Adult | M | 9,2 | 72 | 501042,5024 | 4350614,888 |
| M67 | Na Moltona | 02.05.18 | Spring-18 | Juvenile | Unk | 2,5 | 48 | 501002,3632 | 4350651,574 |
| ng11 | Na Guardis | 17.10.17 | Autumn-17 | Adult | F | 4,3 | 59 | 500135 | 4351239 |
| ng56 | Na Guardis | 17.10.17 | Autumn-17 | Subadult | F | 3,9 | 54 | 500104 | 4351232 |
| ng70 | Na Guardis | 17.10.17 | Autumn-17 | Adult | F | 3,8 | 55 | 500104 | 4351240 |
| N11 | Na Guardis | 04.04.17 | Spring-17 | Adult | M | 7,2 | 63 | 500124 | 4351255 |
| N13 | Na Guardis | 04.04.17 | Spring-17 | Adult | M | 8,4 | 70 | 500127 | 4351246 |
| N22 | Na Guardis | 04.04.17 | Spring-17 | Adult | F | 6,4 | 63 | 500111 | 4351247 |
| N27 | Na Guardis | 04.04.17 | Spring-17 | Adult | F | 5 | 61 | 500152 | 4351262 |
| N29 | Na Guardis | 04.04.17 | Spring-17 | Juvenile | Unk | 2,2 | 45 | 500087 | 4351239 |
| N44 | Na Guardis | 04.04.17 | Spring-17 | Juvenile | F | 2,3 | 48 | 500104 | 4351240 |
| N61A | Na Guardis | 04.04.17 | Spring-17 | Adult | M | 8,2 | 69 | 500140 | 4351245 |
| N70 | Na Guardis | 05.04.17 | Spring-17 | Adult | F | 4,6 | 58 | 500123 | 4351231 |
| N73 | Na Guardis | 05.04.17 | Spring-17 | Adult | F | 6,2 | 64 | 500089 | 4351246 |
| N8 | Na Guardis | 05.04.17 | Spring-17 | Adult | F | 5,9 | 62 | 500115 | 4351229 |
| N84 | Na Guardis | 05.04.17 | Spring-17 | Adult | F | 6,1 | 62 | 500116 | 4351266 |
| N85 | Na Guardis | 05.04.17 | Spring-17 | Adult | F | 4,3 | 57 | 500115 | 4351276 |
| N90 | Na Guardis | 06.04.17 | Spring-17 | Adult | F | 6,7 | 62 | 500097 | 4351239 |
| N99 | Na Guardis | 06.04.17 | Spring-17 | Adult | M | 7,4 | 63 | 500097 | 4351239 |
| NA102 | Na Guardis | 20.10.18 | Autumn-18 | Adult | F | 4,1 | 56 | 500108 | 4351253 |
| NA106 | Na Guardis | 20.10.18 | Autumn-18 | Subadult | M | 5,3 | 61 | 500104 | 4351240 |
| NA111 | Na Guardis | 20.10.18 | Autumn-18 | Adult | M | 9 | 70 | 500097 | 4351239 |
| NA12 | Na Guardis | 20.10.18 | Autumn-18 | Adult | M | 8,2 | 66 | 500111 | 4351247 |
| NA122 | Na Guardis | 20.10.18 | Autumn-18 | Adult | F | 5,6 | 62 | 500140 | 4351245 |
| NA124 | Na Guardis | 20.10.18 | Autumn-18 | Adult | M | 9 | 65 | 500141 | 4351258 |
| NA13 | Na Guardis | 20.10.18 | Autumn-18 | Adult | F | 6,2 | 60 | 500108 | 4351253 |
| NA14 | Na Guardis | 20.10.18 | Autumn-18 | Adult | M | 6,4 | 63 | 500097 | 4351239 |
| NA140 | Na Guardis | 20.10.18 | Autumn-18 | Adult | F | 4,7 | 56 | 500135 | 4351239 |
| NA157 | Na Guardis | 20.10.18 | Autumn-18 | Subadult | F | 3,2 | 53 | 500129 | 4351236 |
| NA160 | Na Guardis | 20.10.18 | Autumn-18 | Adult | F | 6,2 | 64 | 500140 | 4351245 |
| NA46 | Na Guardis | 20.10.18 | Autumn-18 | Adult | M | 7,2 | 69 | 500141 | 4351258 |
| NA49 | Na Guardis | 20.10.18 | Autumn-18 | Subadult | F | 3,3 | 54 | 500133 | 4351261 |
| NA50 | Na Guardis | 20.10.18 | Autumn-18 | Adult | F | 5,1 | 62 | 500115 | 4351276 |
| NA61 | Na Guardis | 20.10.18 | Autumn-18 | Adult | M | 7,1 | 63 | 500108 | 4351253 |
| NA63 | Na Guardis | 20.10.18 | Autumn-18 | Subadult | F | 3,2 | 53 | 500104 | 4351232 |
| NA7 | Na Guardis | 20.10.18 | Autumn-18 | Subadult | M | 4,2 | 55 | 500141 | 4351258 |
| NA93 | Na Guardis | 20.10.18 | Autumn-18 | Adult | M | 6,7 | 64 | 500124 | 4351255 |
| N101 | Na Guardis | 19.04.18 | Spring-18 | Adult | M | 8,2 | 67 | 500115 | 4351276 |
| N23 | Na Guardis | 17.04.18 | Spring-18 | Adult | F | 6,2 | 59 | 500087 | 4351239 |
| N33 | Na Guardis | 18.04.18 | Spring-18 | Adult | M | 6,8 | 65 | 500129 | 4351236 |
| N4 | Na Guardis | 17.04.18 | Spring-18 | Juvenile | M | 1,4 | 39 | 500129 | 4351236 |
| N52 | Na Guardis | 18.04.18 | Spring-18 | Adult | M | 6,7 | 62 | 500096 | 4351256 |
| N60 | Na Guardis | 18.04.18 | Spring-18 | Adult | F | 6,2 | 59 | 500087 | 4351239 |
| N61B | Na Guardis | 18.04.18 | Spring-18 | Adult | M | 8,3 | 66 | 500124 | 4351255 |
| 105ES | En Curt | 01.04.17 | Spring-17 | Juvenile | M | 1,9 | 43 | 503124 | 4347775 |
| 26ES | En Curt | 01.04.17 | Spring-17 | Adult | M | 6 | 66 | 503114 | 4347767 |
| 34ES | En Curt | 01.04.17 | Spring-17 | Adult | M | 8,5 | 75 | 503103 | 4347771 |
| 3ES | En Curt | 01.04.17 | Spring-17 | Subadult | F | 3,4 | 50 | 503101 | 4347755 |
| 53ES | En Curt | 01.04.17 | Spring-17 | Adult | F | 5,1 | 61 | 503115 | 4347786 |
| 57ES | En Curt | 01.04.17 | Spring-17 | Subadult | M | 3,3 | 52 | 503123 | 4347786 |
| 58ES | En Curt | 01.04.17 | Spring-17 | Adult | F | 5,5 | 65 | 503123 | 4347786 |
| 6ES | En Curt | 01.04.17 | Spring-17 | Adult | M | 7,5 | 64 | 503101 | 4347755 |
| 75ES | En Curt | 01.04.17 | Spring-17 | Juvenile | Unk | 1,1 | 37 | 503095 | 4347767 |
| 91ES | En Curt | 01.04.17 | Spring-17 | Adult | M | 5,5 | 62 | 503109 | 4347760 |
| ES16 | En Curt | 16.10.18 | Autumn-18 | Adult | M | 7,6 | 68 | 503129 | 4347778 |
| ES43 | En Curt | 16.10.18 | Autumn-18 | Adult | M | 6,4 | 65 | 503115 | 4347777 |
| ES44 | En Curt | 11.10.18 | Autumn-18 | Adult | M | 7,9 | 70 | 503115 | 4347783 |
| ES47 | En Curt | 09.10.18 | Autumn-18 | Subadult | F | 3,7 | 56 | 503109 | 4347760 |
| ES63 | En Curt | 09.10.18 | Autumn-18 | Subadult | F | 3,3 | 53 | 503124 | 4347775 |
| ES65 | En Curt | 11.10.18 | Autumn-18 | Adult | M | 9,1 | 72 | 503125 | 4347769 |
| ES68 | En Curt | 09.10.18 | Autumn-18 | Subadult | F | 4 | 54 | 503095 | 4347767 |
| ES70 | En Curt | 09.10.18 | Autumn-18 | Subadult | F | 4 | 55 | 503115 | 4347777 |

**Table S2:** Primers used for amplification of the region V3-V4 of 16S rRNA.

| Primer name | full sequence 5'-3' |
|---|---|
| V3-V4-forward | TCGTCGGCAGCGTCAGATGTGTATAAGAGACAGCCTACGGGNGGCWGCAG |
| V3-V4-forward+1 | TCGTCGGCAGCGTCAGATGTGTATAAGAGACAGNCCTACGGGNGGCWGCAG |
| V3-V4-forward+2 | TCGTCGGCAGCGTCAGATGTGTATAAGAGACAGNNCCTACGGGNGGCWGCAG |
| V3-V4-forward+3 | TCGTCGGCAGCGTCAGATGTGTATAAGAGACAGNNNCCTACGGGNGGCWGCAG |
| V3-V4-forward+4 | TCGTCGGCAGCGTCAGATGTGTATAAGAGACAGNNNNCCTACGGGNGGCWGCAG |
| V3-V4-reverse | GTCTCGTGGGCTCGGAGATGTGTATAAGAGACAGGACTACHVGGGTATCTAATCC |
| V3-V4-reverse+1 | GTCTCGTGGGCTCGGAGATGTGTATAAGAGACAGNGACTACHVGGGTATCTAATCC |
| V3-V4-reverse+2 | GTCTCGTGGGCTCGGAGATGTGTATAAGAGACAGNNGACTACHVGGGTATCTAATCC |
| V3-V4-reverse+3 | GTCTCGTGGGCTCGGAGATGTGTATAAGAGACAGNNNGACTACHVGGGTATCTAATCC |
| V3-V4-reverse+4 | GTCTCGTGGGCTCGGAGATGTGTATAAGAGACAGNNNNGACTACHVGGGTATCTAATCC |

**Table S3:** Abundance matrix of ASVs found in the negative PCR controls and mock communtiies

| | PCR controls | | Mock communities | | | | | |
|---|---|---|---|---|---|---|---|---|
| ASV | NTC1 | NTC2 | 276D | 277D | Total_seq_counts_Lizards | Total_seq_counts_PCR_controls | Total_seq_counts_Mock | Taxonomy |
| a41b72842290a9e865efc83d63ad2818 | 0 | 0 | 5416 | 16036 | 0 | 0 | 21452 | p__Firmicutes; c__Bacilli; o__Lactobacillales; f__Streptococcaceae; g__Streptococcus; s__ |
| 591ed1d6afd1e79ae33d6b6cd0c6b185 | 144 | 80 | 5080 | 15717 | 6417 | 224 | 20797 | p__Proteobacteria; c__Gammaproteobacteria; o__Enterobacteriales; f__Enterobacteriaceae |
| be9e7e2786b3ae134712e7b6bedc2259 | 0 | 0 | 3813 | 14677 | 7 | 0 | 18490 | p__Firmicutes; c__Bacilli; o__Bacillales; f__Staphylococcaceae; g__Staphylococcus |
| 67dfacf96a38558ee3a659a1aa141fd5 | 0 | 0 | 12385 | 925 | 0 | 0 | 13310 | p__Proteobacteria; c__Epsilonproteobacteria; o__Campylobacterales; f__Helicobacteraceae; g__Helicobacter; s__pylori |
| dee86d463f6a6a4bad3f90c961d4150b | 0 | 0 | 2966 | 8911 | 0 | 0 | 11877 | p__Proteobacteria; c__Alphaproteobacteria; o__Rhodobacterales; f__Rhodobacteraceae; g__Rhodobacter; s__sphaeroides |
| 29fde6ffd0a41547e218d84de8f4fa03 | 0 | 0 | 10289 | 134 | 0 | 0 | 10423 | p__Bacteroidetes; c__Bacteroidia; o__Bacteroidales; f__Bacteroidaceae; g__Bacteroides; s__ |
| a3a045b51f6ba1e973a9750f7061edbb | 0 | 0 | 6578 | 2744 | 0 | 0 | 9322 | p__Firmicutes; c__Clostridia; o__Clostridiales; f__Clostridiaceae; g__Clostridium; s__butyricum |
| 3876207928742711d18645217094065b | 0 | 0 | 8175 | 582 | 0 | 0 | 8757 | p__Proteobacteria; c__Gammaproteobacteria; o__Pseudomonadales; f__Moraxellaceae; g__Acinetobacter |
| 63dcf1c7a95370bfe7e6b7e39eb9343b | 0 | 0 | 7265 | 492 | 0 | 0 | 7757 | p__Proteobacteria; c__Betaproteobacteria; o__Neisseriales; f__Neisseriaceae; g__Neisseria; s__cinerea |
| c804f27343fbd3504b0ef0766a3942fe | 0 | 0 | 4857 | 2121 | 17 | 0 | 6978 | p__Proteobacteria; c__Gammaproteobacteria; o__Pseudomonadales; f__Pseudomonadaceae |
| bfd4d446a5b960b55647a573d459ff58 | 0 | 0 | 4770 | 1974 | 0 | 0 | 6744 | p__Firmicutes; c__Bacilli; o__Lactobacillales; f__Streptococcaceae; g__Streptococcus; s__agalactiae |
| e5084546676d262b0af7425e5b7a3a7e | 0 | 0 | 4291 | 1764 | 56 | 0 | 6055 | p__Firmicutes; c__Bacilli; o__Bacillales; f__Bacillaceae; g__Bacillus |
| e3e618b6c0b173feb5ba627d9a96d485 | 0 | 0 | 5652 | 255 | 825 | 0 | 5907 | p__Firmicutes; c__Bacilli; o__Lactobacillales; f__Lactobacillaceae; g__Lactobacillus; s__ |
| 475aed25e995894a8ce8204c1c6b63d3 | 0 | 0 | 5702 | 35 | 0 | 0 | 5737 | p__[Thermi]; c__Deinococci; o__Deinococcales; f__Deinococcaceae; g__Deinococcus; s__ |
| b05f5459e068b529b2c41e659efdef34 | 0 | 0 | 4509 | 218 | 14 | 0 | 4727 | p__Actinobacteria; c__Actinobacteria; o__Actinomycetales; f__Propionibacteriaceae; g__Propionibacterium; s__acnes |
| cfc605a1be0fec6ff94c797a789eea8c | 0 | 0 | 4384 | 176 | 0 | 0 | 4560 | p__Firmicutes; c__Bacilli; o__Bacillales; f__Listeriaceae; g__Listeria |
| 162d380468428be715f6403345eaa137 | 0 | 0 | 3645 | 0 | 0 | 0 | 3645 | p__Firmicutes; c__Bacilli; o__Bacillales; f__Staphylococcaceae; g__Staphylococcus |
| 2e3daa713ba0eafcae4ae5d65a774f9f | 0 | 0 | 2562 | 20 | 0 | 0 | 2582 | p__Proteobacteria; c__Gammaproteobacteria; o__Pseudomonadales; f__Pseudomonadaceae; g__Pseudomonas; s__ |
| 0f6d46c70548d4ce709802b6e3162f6f | 0 | 0 | 2292 | 12 | 81 | 0 | 2304 | p__Actinobacteria; c__Actinobacteria; o__Actinomycetales; f__Actinomycetaceae; g__Actinomyces; s__ |
| bca88c697e57d95ec75e2af02ce992e2 | 0 | 0 | 1180 | 2 | 0 | 0 | 1182 | p__Firmicutes; c__Bacilli; o__Lactobacillales; f__Enterococcaceae; g__Enterococcus; s__ |
| 09cb0bcce9ef9b5dcf6b9d7a6c663d83 | 0 | 0 | 785 | 11 | 0 | 0 | 796 | p__Firmicutes; c__Bacilli; o__Lactobacillales; f__Streptococcaceae; g__Streptococcus |
| 5160e83ed49366187ffd39b06bc14897 | 0 | 0 | 521 | 0 | 0 | 0 | 521 | p__Bacteroidetes; c__Bacteroidia; o__Bacteroidales; f__Bacteroidaceae; g__Bacteroides; s__ |
| 05f3c856ceb915f6606f1e0aca4fa8ae | 1485 | 1768 | 2 | 2 | 2507 | 3253 | 4 | p__Proteobacteria; c__Gammaproteobacteria; o__Enterobacteriales; f__Enterobacteriaceae; g__Gluconacetobacter; s__ |
| d641d1422558101f892611b0fe1194f0 | 70 | 188 | 2 | 0 | 84 | 258 | 2 | p__Bacteroidetes; c__Bacteroidia; o__Bacteroidales; f__Bacteroidaceae; g__Bacteroides; s__ |
| a32367cd48ba6aec652c13719f6d23f2 | 227 | 326 | 0 | 0 | 388 | 553 | 0 | p__Proteobacteria; c__Gammaproteobacteria; o__Pseudomonadales; f__Pseudomonadaceae; g__Pseudomonas; s__veronii |
| 3448d1bf6cccbbf7b14c71b09817cba4 | 245 | 123 | 0 | 0 | 310 | 368 | 0 | p__Proteobacteria; c__Gammaproteobacteria; o__Enterobacteriales; f__Enterobacteriaceae; g__Gluconacetobacter; s__ |
| 8bb6ba264b60216d22b221d5c6c0992e | 37 | 177 | 0 | 0 | 109 | 214 | 0 | p__Proteobacteria; c__Gammaproteobacteria; o__Pseudomonadales; f__Pseudomonadaceae; g__Pseudomonas |
| eb254f006a34d1ea0e5641906aa59747 | 54 | 120 | 0 | 0 | 81 | 174 | 0 | p__Firmicutes; c__Bacilli; o__Lactobacillales; f__Carnobacteriaceae; g__Carnobacterium; s__ |
| 70ed1793e682927f916f0b26a3881b14 | 0 | 93 | 0 | 0 | 0 | 93 | 0 | p__Bacteroidetes; c__Flavobacteriia; o__Flavobacteriales; f__Flavobacteriaceae; g__Capnocytophaga; s__ |
| 0e8bc97b3b6003e64d81ee7bab83c8a5 | 0 | 80 | 0 | 0 | 10 | 80 | 0 | p__Verrucomicrobia; c__Verrucomicrobiae; o__Verrucomicrobiales; f__Verrucomicrobiaceae; g__Akkermansia; s__muciniphila |
| d0ab2c15400fe710288526c9a33083fb | 0 | 72 | 0 | 0 | 17 | 72 | 0 | p__Firmicutes; c__Bacilli; o__Lactobacillales; f__Streptococcaceae; g__Streptococcus; s__infantis |
| 2de0f9fd09640e74b2b452a260409fab | 47 | 10 | 0 | 0 | 23 | 57 | 0 | p__Firmicutes; c__Bacilli; o__Bacillales; f__Bacillaceae |
| 4a91fbea81d4fc6aad46838da248268b | 0 | 57 | 0 | 0 | 12 | 57 | 0 | p__Proteobacteria; c__Gammaproteobacteria; o__Pseudomonadales; f__Pseudomonadaceae; g__Pseudomonas; s__veronii |
| 5a8c63a1400afebcd134e100c758112e | 48 | 0 | 0 | 0 | 0 | 48 | 0 | p__Bacteroidetes; c__Bacteroidia; o__Bacteroidales; f__Bacteroidaceae; g__Bacteroides; s__eggerthii |
| dde001cc50f5d8473e62de2fdb7d83c1 | 43 | 0 | 0 | 0 | 13 | 43 | 0 | p__Proteobacteria; c__Betaproteobacteria; o__Burkholderiales; f__Comamonadaceae; g__Curvibacter; s__ |
| 7700d116dce04ef0bbb3fc9a2deb328a | 41 | 0 | 0 | 0 | 3 | 41 | 0 | p__Firmicutes; c__Clostridia; o__Clostridiales; f__Lachnospiraceae; g__Shuttleworthia; s__ |
| 0072b0040e39e5d64afc5038682721dc | 40 | 0 | 0 | 0 | 0 | 40 | 0 | p__Firmicutes; c__Clostridia; o__Clostridiales; f__Ruminococcaceae; g__Oscillospira; s__ |
| 4ba5a86e53a755c6e3eb2f70a300339f | 0 | 38 | 0 | 0 | 0 | 38 | 0 | p__Firmicutes; c__Clostridia; o__Clostridiales; f__Ruminococcaceae |
| 12f940f9951b6fb0cbbb6b40d6b71776 | 0 | 37 | 0 | 0 | 0 | 37 | 0 | p__Proteobacteria; c__Alphaproteobacteria; o__Rhizobiales; f__Bradyrhizobiaceae |
| 6bd95460524b199c4a65914e8749de8c | 37 | 0 | 0 | 0 | 0 | 37 | 0 | p__Firmicutes; c__Clostridia; o__Clostridiales; f__Veillonellaceae; g__Phascolarctobacterium; s__ |
| bd0d1ce776efcbb887d89a9c31dd8707 | 36 | 0 | 0 | 0 | 0 | 36 | 0 | p__Bacteroidetes; c__Bacteroidia; o__Bacteroidales; f__S24-7; g__; s__ |
| 692ef3111a5143c64d5ec9d26603cc5a | 35 | 0 | 0 | 0 | 9 | 35 | 0 | p__Proteobacteria; c__Betaproteobacteria; o__Burkholderiales; f__Burkholderiaceae; g__Burkholderia; s__ |
| 4b58a918963708863f456bd44ae56b5b | 34 | 0 | 0 | 0 | 0 | 34 | 0 | p__Firmicutes; c__Clostridia; o__Clostridiales; f__Lachnospiraceae; g__Blautia; s__ |
| 2397358d6f6eaff8d40ccb98f4fbebd9 | 0 | 30 | 0 | 0 | 0 | 30 | 0 | p__Proteobacteria; c__Alphaproteobacteria; o__Rhizobiales; f__Brucellaceae; g__Pseudochrobactrum; s__ |
| 33db7b189dcfe16b551c393c56f3b5d5 | 30 | 0 | 0 | 0 | 0 | 30 | 0 | p__Firmicutes; c__Clostridia; o__Clostridiales; f__Lachnospiraceae; g__Coprococcus; s__ |
| 9f22da56e873ade2c09013cae621e715 | 0 | 30 | 0 | 0 | 0 | 30 | 0 | p__Proteobacteria; c__Betaproteobacteria; o__Burkholderiales; f__Oxalobacteraceae; g__Ralstonia; s__ |
| e748d50e065533ffb80c44c7ce9b78d4 | 29 | 0 | 0 | 0 | 2 | 29 | 0 | p__Proteobacteria; c__Gammaproteobacteria; o__Enterobacteriales; f__Enterobacteriaceae; g__Proteus; s__ |
| 0256efed814c10383ad0908793c5e2ff | 28 | 0 | 0 | 0 | 0 | 28 | 0 | p__Bacteroidetes; c__Bacteroidia; o__Bacteroidales; f__Bacteroidaceae; g__Bacteroides; s__caccae |
| 3c2934680d564f24be3099a620adb243 | 0 | 28 | 0 | 0 | 0 | 28 | 0 | p__Actinobacteria; c__Actinobacteria; o__Actinomycetales; f__Micrococcaceae; g__Rothia; s__mucilaginosa |
| e3c224e148980246988888a3bddbbb22 | 27 | 0 | 0 | 0 | 4 | 27 | 0 | p__Fusobacteria; c__Fusobacteriia; o__Fusobacteriales; f__Leptotrichiaceae; g__; s__ |
| d777bbaa70de5ae5aed5d5f715ce0201 | 0 | 26 | 0 | 0 | 16 | 26 | 0 | p__Actinobacteria; c__Actinobacteria; o__Bifidobacteriales; f__Bifidobacteriaceae; g__Bifidobacterium; s__thermacidophilum |
| 34588d48b1f866ee34ee3225e78009e8 | 26 | 0 | 0 | 0 | 14 | 26 | 0 | p__Proteobacteria; c__Gammaproteobacteria; o__Enterobacteriales; f__Enterobacteriaceae; g__Gluconacetobacter; s__ |
| a1c6504e8981cc8645a7f588e5f8408c | 0 | 25 | 0 | 0 | 0 | 25 | 0 | p__Firmicutes; c__Bacilli; o__Gemellales; f__Gemellaceae; g__; s__ |
| e9a43d976dfa1e002c02d5970c8b4255 | 0 | 25 | 0 | 0 | 0 | 25 | 0 | p__Proteobacteria; c__Betaproteobacteria; o__Burkholderiales; f__Alcaligenaceae; g__Denitrobacter; s__ |
| a769fd25cb4cb3e65fb0b5f752ddf4cb | 24 | 0 | 0 | 0 | 6 | 24 | 0 | p__Proteobacteria; c__Betaproteobacteria; o__Burkholderiales; f__Comamonadaceae; g__Pelomonas; s__ |
| 932d70b76d839b7553bb90df617aea66 | 0 | 21 | 0 | 0 | 15 | 21 | 0 | p__Proteobacteria; c__Gammaproteobacteria; o__Xanthomonadales; f__Xanthomonadaceae; g__Stenotrophomonas; s__geniculata |
| 454946e0565af9ca0cec72a700184f63 | 20 | 0 | 0 | 0 | 0 | 20 | 0 | p__Proteobacteria; c__Alphaproteobacteria; o__Rhizobiales; f__Methylobacteriaceae; g__Methylobacterium; s__ |
| d06dc00291933c496e93692f04461a33 | 0 | 18 | 0 | 0 | 0 | 18 | 0 | p__Firmicutes; c__Clostridia; o__Clostridiales; f__Lachnospiraceae; g__Butyrivibrio; s__ |
| 1f6b9f0a208495aae4792e174153c508 | 17 | 0 | 0 | 0 | 0 | 17 | 0 | p__Firmicutes; c__Clostridia; o__Clostridiales; f__Lachnospiraceae |
| ce732db6fdb0acd1dcfb1bc18665c8ce | 17 | 0 | 0 | 0 | 0 | 17 | 0 | p__Proteobacteria; c__Alphaproteobacteria; o__Sphingomonadales; f__Sphingomonadaceae; g__Sphingobium; s__ |
| e39d99da02868537980a95b2b42f5525 | 14 | 0 | 0 | 0 | 0 | 14 | 0 | p__Verrucomicrobia; c__Verrucomicrobiae; o__Verrucomicrobiales; f__Verrucomicrobiaceae; g__Akkermansia; s__muciniphila |
| e6d5a138e135922560122d359a014d1a | 0 | 10 | 0 | 0 | 3 | 10 | 0 | p__Proteobacteria; c__Gammaproteobacteria; o__Enterobacteriales; f__Enterobacteriaceae |

**Table S4:** List of ASVs that signifcantly discriminated among islets based on both seasonal datsets and according to both indval and LEfSe analyses

| | | LEFSE | | | | | | | |
|---|---|---|---|---|---|---|---|---|---|
| | | AUTUMN | | | | SPRING | | | |
| Figure_5_ID | ASV | log_high_class | Enrichment Class | LDA | p.value | log_high_ | Class | LDA | p.value |
| **ASV1** | 3bb7551ec97a9bacd2eb126b8521666a | 4,636 | NM | 4,31 | 0,00 | 4,67 | NM | 4,30 | 0,002 |
| **ASV2** | 7ae97f9f54cf6fc0d22f6b73b5346c76 | 4,070 | NM | 3,61 | 0,00 | 4,20 | NM | 3,82 | 0,001 |
| **ASV3** | b5c97bb118b54e8fc1afbba09dcb8d82 | 3,728 | NM | 3,42 | 0,00 | 4,05 | NM | 3,76 | 0,000 |
| **ASV4** | 59b5717fe298ca1e4e8c5f643993a1d2 | 4,328 | EC | 4,02 | 0,00 | 4,44 | EC | 4,13 | 0,000 |
| **ASV5** | d2103ec4413ebc45d4baced9b537d0a1 | 4,374 | EC | 4,06 | 0,00 | 3,83 | EC | 3,50 | 0,000 |
| **ASV6** | fe40f1a7a6ddfe082963689f85ae01cf | 3,674 | EC | 3,37 | 0,00 | 4,02 | EC | 3,68 | 0,000 |
| **ASV7** | 7575c5d1e36ca75c5a8171a29077f1d8 | 3,441 | EC | 3,13 | 0,00 | 4,03 | EC | 3,72 | 0,002 |
| **ASV8** | 657cf34751ede24a8bb901e90d390824 | 3,901 | EC | 3,59 | 0,00 | 3,91 | EC | 3,60 | 0,000 |
| **ASV9** | a18933a9733431c0323b81966e3e6c11 | 4,032 | EC | 3,76 | 0,00 | 3,48 | EC | 3,19 | 0,000 |
| **ASV10** | c44299bf6bc56e6d11986b8a7725db91 | 3,569 | EC | 3,30 | 0,00 | 3,83 | EC | 3,52 | 0,000 |
| **ASV11** | 32346b07c25298dace2646644e3dfca5 | 3,614 | EC | 3,30 | 0,00 | 3,47 | EC | 3,20 | 0,001 |
| **ASV12** | b62fd07e2c96423bab395852c88fb147 | 3,750 | EC | 3,41 | 0,00 | 3,33 | EC | 3,04 | 0,000 |
| **ASV13** | 87716c92b164605186d00734bea9e430 | 3,447 | EC | 3,20 | 0,00 | 3,46 | EC | 3,18 | 0,000 |
| **ASV14** | ae98aa63a4f9a9f9081e7e41b7db8af7 | 3,618 | EC | 3,28 | 0,00 | 3,34 | EC | 3,04 | 0,000 |

| | INDVAL | | | | | | | | | |
|---|---|---|---|---|---|---|---|---|---|---|
| | AUTUMN | | | | | | | | | |
| | indval | | | | Relfreq | | | Relabund | | |
| Figure_5_ID | EC | NM | NG | pval | EC | NM | NG | EC | NM | NG |
| **ASV1** | 0,018 | 0,518 | 0,2 | 0,009 | 0,25 | 0,897 | 0,571 | 0,072 | 0,578 | 0,349 |
| **ASV2** | 0,207 | 0,511 | 0,228 | 0,004 | 0,875 | 1 | 0,905 | 0,237 | 0,511 | 0,252 |
| **ASV3** | 0 | 0,531 | 0,071 | 0,01 | 0 | 0,759 | 0,238 | 0 | 0,7 | 0,3 |
| **ASV4** | 0,972 | 0,001 | 0,001 | 0,001 | 1 | 0,069 | 0,095 | 0,972 | 0,015 | 0,014 |
| **ASV5** | 1 | 0 | 0 | 0,001 | 1 | 0 | 0 | 1 | 0 | 0 |
| **ASV6** | 0,647 | 0,144 | 0 | 0,002 | 0,875 | 0,552 | 0 | 0,74 | 0,26 | 0 |
| **ASV7** | 0,604 | 0,02 | 0 | 0,001 | 0,75 | 0,103 | 0 | 0,805 | 0,195 | 0 |
| **ASV8** | 1 | 0 | 0 | 0,001 | 1 | 0 | 0 | 1 | 0 | 0 |
| **ASV9** | 0,625 | 0 | 0 | 0,001 | 0,625 | 0 | 0 | 1 | 0 | 0 |
| **ASV10** | 0,838 | 0,001 | 0 | 0,001 | 0,875 | 0,034 | 0 | 0,957 | 0,043 | 0 |
| **ASV11** | 0,864 | 0,001 | 0 | 0,001 | 0,875 | 0,103 | 0,048 | 0,988 | 0,008 | 0,004 |
| **ASV12** | 0,75 | 0 | 0 | 0,001 | 0,75 | 0 | 0 | 1 | 0 | 0 |
| **ASV13** | 0,724 | 0,001 | 0 | 0,001 | 0,75 | 0,034 | 0 | 0,966 | 0,034 | 0 |
| **ASV14** | 0,947 | 0,002 | 0 | 0,001 | 1 | 0,034 | 0 | 0,947 | 0,053 | 0 |

| | INDVAL | | | | | | | | | |
|---|---|---|---|---|---|---|---|---|---|---|
| | SPRING | | | | | | | | | |
| | indval | | | | Relfreq | | | Relabund | | |
| Figure_5_ID | EC | NM | NG | pval.1 | EC | NM | NG | EC | NM | NG |
| **ASV1** | 0,008 | 0,759 | 0,053 | 0,001 | 0,25 | 0,938 | 0,333 | 0,032 | 0,81 | 0,158 |
| **ASV2** | 0,064 | 0,576 | 0,304 | 0,003 | 0,625 | 1 | 0,944 | 0,103 | 0,576 | 0,321 |
| **ASV3** | 0 | 0,676 | 0,176 | 0,002 | 0 | 0,875 | 0,778 | 0 | 0,773 | 0,227 |
| **ASV4** | 0,818 | 0,006 | 0,003 | 0,001 | 0,875 | 0,125 | 0,167 | 0,935 | 0,044 | 0,021 |
| **ASV5** | 0,853 | 0,002 | 0 | 0,001 | 0,875 | 0,063 | 0 | 0,974 | 0,026 | 0 |
| **ASV6** | 0,633 | 0,079 | 0,005 | 0,002 | 0,75 | 0,625 | 0,167 | 0,844 | 0,127 | 0,029 |
| **ASV7** | 0,585 | 0,012 | 0 | 0,002 | 0,625 | 0,188 | 0 | 0,936 | 0,064 | 0 |
| **ASV8** | 0,875 | 0 | 0 | 0,001 | 0,875 | 0 | 0 | 1 | 0 | 0 |
| **ASV9** | 0,625 | 0 | 0 | 0,001 | 0,625 | 0 | 0 | 1 | 0 | 0 |
| **ASV10** | 0,875 | 0 | 0 | 0,001 | 0,875 | 0 | 0 | 1 | 0 | 0 |
| **ASV11** | 0,521 | 0,036 | 0,004 | 0,008 | 0,75 | 0,125 | 0,222 | 0,694 | 0,288 | 0,017 |
| **ASV12** | 0,578 | 0,001 | 0,012 | 0,002 | 0,75 | 0,063 | 0,056 | 0,77 | 0,019 | 0,211 |
| **ASV13** | 0,768 | 0 | 0,013 | 0,001 | 0,875 | 0,063 | 0,111 | 0,877 | 0,004 | 0,119 |
| **ASV14** | 0,98 | 0,001 | 0 | 0,001 | 1 | 0,063 | 0 | 0,98 | 0,02 | 0 |

| | Taxonomy | | | | | |
|---|---|---|---|---|---|---|
| Figure_5_ID | Phylum | Class | Order | Family | Genus | Species |
| **ASV1** | Bacteroidetes | Bacteroidia | Bacteroidales | Bacteroidaceae | Bacteroides | |
| **ASV2** | Bacteroidetes | Bacteroidia | Bacteroidales | Porphyromonadaceae | Parabacteroides | P. gordonii |
| **ASV3** | Bacteroidetes | Bacteroidia | Bacteroidales | [Odoribacteraceae] | Odoribacter | |
| **ASV4** | Bacteroidetes | Bacteroidia | Bacteroidales | Bacteroidaceae | Bacteroides | |
| **ASV5** | Bacteroidetes | Bacteroidia | Bacteroidales | Porphyromonadaceae | Parabacteroides | P. gordonii |
| **ASV6** | Bacteroidetes | Bacteroidia | Bacteroidales | Bacteroidaceae | Bacteroides | |
| **ASV7** | Bacteroidetes | Bacteroidia | Bacteroidales | Bacteroidaceae | Bacteroides | B. ovatus |
| **ASV8** | Bacteroidetes | Bacteroidia | Bacteroidales | Porphyromonadaceae | Parabacteroides | P. distasonis |
| **ASV9** | Verrucomicrobia | Verrucomicrobiae | Verrucomicrobiales | Verrucomicrobiaceae | Akkermansia | |
| **ASV10** | Bacteroidetes | Bacteroidia | Bacteroidales | Rikenellaceae | | |
| **ASV11** | Bacteroidetes | Bacteroidia | Bacteroidales | Porphyromonadaceae | Porphyromonas | |
| **ASV12** | Bacteroidetes | Bacteroidia | Bacteroidales | Porphyromonadaceae | Parabacteroides | |
| **ASV13** | Bacteroidetes | Bacteroidia | Bacteroidales | [Odoribacteraceae] | Odoribacter | |
| **ASV14** | Proteobacteria | Alphaproteobacteria | Rickettsiales | | | |

**Table S5**: Stable isotopes estimated on 71 specimens collected during spring 2016.

| Date | Islet | Specimen_ID | Delta 13C x 1000 | Mass Fraction C x 100 | Delta 15N x 1000 | Mass Fraction N x 100 | Sample Weight | ID_WEIGHT_gr | IF_LENGTH_cm | SEX |
|---|---|---|---|---|---|---|---|---|---|---|
| 14/04/2016 | NM | 1085 | -23,23 | 49,36 | 6,69 | 14,48 | 0.306 | 10 | 7,2 | M |
| 14/04/2016 | NM | 1086 | NA | NA | NA | NA | NA | 9 | 6,8 | M |
| 14/04/2016 | NM | 1088 | -22,61 | 43,71 | 6,83 | 10,44 | 0.173 | 9 | 6,9 | F |
| 14/04/2016 | NM | 1089 | NA | NA | NA | NA | NA | 3,9 | 5,1 | F |
| 14/04/2016 | NM | 1091 | NA | NA | NA | NA | NA | 10,3 | 7,1 | M |
| 14/04/2016 | NM | 1092 | -22,04 | 49,81 | 8,31 | 13,43 | 0.295 | NA | NA | NA |
| 14/04/2016 | NM | 1093 | NA | NA | NA | NA | NA | 6,7 | 6,5 | F |
| 14/04/2016 | NM | 1099 | -24,24 | 48,3 | 5,04 | 13,94 | 0.309 | 9,2 | 7,3 | M |
| 14/04/2016 | NM | 1100 | -24,14 | 47,59 | 6,04 | 14 | 0.130 | 4,2 | 6,1 | F |
| 14/04/2016 | NM | 1101 | -23,48 | 44,82 | 6,54 | 13,33 | 0.166 | 5,8 | 6,1 | F |
| 14/04/2016 | NM | 1102 | -24,55 | 48,69 | 6,75 | 12,59 | 0.309 | 6,5 | 6,7 | F |
| 14/04/2016 | NM | 1103 | -22,94 | 49,27 | 6,47 | 11,3 | 0.246 | 6,2 | 6,3 | F |
| 14/04/2016 | NM | 1104 | -23,37 | 47,97 | 6,85 | 13,38 | 0.298 | 10 | 7,4 | M |
| 14/04/2016 | NM | 1105 | NA | NA | NA | NA | NA | 4,4 | 6 | F |
| 14/04/2016 | NM | 1106 | -23,85 | 50,76 | 6,77 | 14,48 | 0.323 | 9,5 | 7 | M |
| 14/04/2016 | NM | 1107 | -24,25 | 48,96 | 5,99 | 14,57 | 0.300 | 6 | 6,5 | F |
| 14/04/2016 | NM | 1108 | -24,02 | 49,64 | 5,59 | 14,56 | 0.329 | 6 | 6,4 | F |
| 14/04/2016 | NM | 1109 | -23,43 | 47,56 | 6,05 | 13,36 | 0.287 | 8,2 | 7,1 | M |
| 14/04/2016 | NM | 1116 | -23,67 | 48,56 | 5,6 | 13,29 | 0.308 | 10 | 7,5 | M |
| 14/04/2016 | NM | 1133 | -23,82 | 47,85 | 6,13 | 14,27 | 0.295 | 7 | 6,9 | M |
| 14/04/2016 | EC | 185 | -21,73 | 51,58 | 10,6 | 14,97 | 0.299 | 9,5 | 7 | M |
| 14/04/2016 | EC | 187 | -23,79 | 50,6 | 13,42 | 14,47 | 0.141 | 6,4 | 6,5 | F |
| 14/04/2016 | EC | 188 | -23,96 | 49,83 | 10,55 | 14,26 | 0.244 | 7 | 6,5 | F |
| 14/04/2016 | EC | 190 | -23,43 | 50,44 | 10,8 | 14,48 | 0.255 | 7 | 6,6 | F |
| 14/04/2016 | EC | 191 | -24,62 | 53,16 | 11,8 | 12,28 | 0.385 | 7,6 | 7 | F |
| 14/04/2016 | EC | 193 | -23,85 | 45,36 | 12,21 | 12,93 | 0.300 | 6,8 | 6,8 | F |
| 14/04/2016 | EC | 194 | -21,82 | 56,23 | 10,11 | 11,98 | 0.185 | 8,7 | 6,8 | F |
| 14/04/2016 | EC | 195 | -22,49 | 48,27 | 11,09 | 14,06 | 0.238 | 6,4 | 6,5 | F |
| 14/04/2016 | EC | 196 | -22,89 | 49,05 | 10,51 | 13,8 | 0.350 | 7,2 | 6,8 | F |
| 14/04/2016 | EC | 197 | -23,58 | 47,67 | 9,44 | 14,39 | 0.213 | 11 | 7,4 | M |
| 14/04/2016 | EC | 198 | -21,77 | 36,18 | 8,75 | 11,05 | 0.141 | 8,4 | 6,7 | F |
| 14/04/2016 | EC | 199 | -23,13 | 50,01 | 9,45 | 15,22 | 0.302 | 9 | 7,1 | M |
| 14/04/2016 | EC | 204 | -21,26 | 50,56 | 8,74 | 15,33 | 0.303 | 11,8 | 7,3 | M |
| 14/04/2016 | EC | 205 | -21,83 | 48,96 | 9,55 | 14,71 | 0.294 | 9 | 7 | M |
| 15/04/2016 | EC | 85 | -22,95 | 49,32 | 9,99 | 14,09 | 0.283 | 10 | 7,2 | M |
| 15/04/2016 | EC | 86 | -23,37 | 47,94 | 10,09 | 13,41 | 0.290 | 6,7 | 6,5 | F |
| 15/04/2016 | EC | 87 | -23,19 | 51,7 | 9,92 | 14,9 | 0.299 | 10,7 | 7,4 | M |
| 15/04/2016 | EC | 91 | -21,13 | 49,12 | 10,34 | 14,13 | 0.242 | 10,6 | 7,1 | M |
| 15/04/2016 | EC | 92 | -24,34 | 39,53 | 11,2 | 11,6 | 0.354 | 6,7 | 6,5 | F |
| 15/04/2016 | EC | 93 | -21,96 | 53,66 | 8,78 | 15,32 | 0.307 | 11,2 | 7,7 | M |
| 15/04/2016 | EC | 94 | -23,37 | 47,33 | 10,05 | 12,89 | 0.235 | 9,1 | 6,8 | M |
| 15/04/2016 | EC | 96 | -23,87 | 53,05 | 10,32 | 13,93 | 0.378 | 7 | 6,4 | F |
| 15/04/2016 | EC | 1B | -22,49 | 51,79 | 9,63 | 14,99 | 0.336 | 9,6 | 7,1 | M |
| 15/04/2016 | EC | 3B | -19,33 | 49,71 | 9,84 | 14,61 | 0.316 | 9,7 | 6,7 | M |
| 15/04/2016 | EC | 4B | -23,04 | 50,01 | 10,15 | 14,3 | 0.303 | 11,3 | 7,2 | M |
| 15/04/2016 | EC | 8B | -21,82 | 51,13 | 9,68 | 14,61 | 0.309 | 10,4 | 7,2 | M |
| 15/04/2016 | EC | 11B | -21,05 | 48,94 | 9,52 | 13,46 | 0.325 | 9,5 | 7 | M |
| 15/04/2016 | EC | 13B | -22,55 | 50,05 | 9,73 | 14,75 | 0.230 | 9,2 | 6,8 | M |
| 15/04/2016 | EC | 14B | -21,84 | 49,83 | 10,22 | 14,12 | 0.335 | 7,2 | 6,4 | M |
| 15/04/2016 | EC | 15B | -22,62 | 54,49 | 10,18 | 12,55 | 0.270 | 5 | 6,4 | F |
| 18/05/2016 | NG | 3 | -24,46 | 50,36 | 5,6 | 15,21 | 0.211 | 6 | 5,9 | M |
| 18/05/2016 | NG | 4 | -24,17 | 48,73 | 5,88 | 14,53 | 0.296 | 9 | 7 | M |
| 18/05/2016 | NG | 8 | -24,4 | 50,35 | 5,44 | 15,34 | 0.306 | 8,6 | 7,6 | M |
| 18/05/2016 | NG | 9 | -24,17 | 44,05 | 5,48 | 13,34 | 0.329 | 7,8 | 6,5 | M |
| 18/05/2016 | NG | 10 | -24,91 | 51,77 | 6,17 | 15,13 | 0.249 | 6,2 | 6,3 | M |
| 18/05/2016 | NG | 11 | -23,6 | 51,57 | 5,7 | 14,62 | 0.298 | 5,9 | 6,2 | F |
| 19/05/2016 | NG | 14 | -24,23 | 50,25 | 5,66 | 14,54 | 0.163 | 8,5 | 6,8 | M |
| 19/05/2016 | NG | 15 | -24,22 | 51,05 | 5,08 | 15,13 | 0.289 | 8 | 6,8 | M |
| 19/05/2016 | NG | 16 | NA | NA | NA | NA | NA | 7,3 | 6,8 | F |
| 19/05/2016 | NG | 18 | -24,27 | 51,34 | 5,28 | 14,88 | 0.345 | 4,6 | 5,7 | F |
| 19/05/2016 | NG | 20 | -24,86 | 50,9 | 5,57 | 14,99 | 0.304 | 8 | 6,8 | M |
| 19/05/2016 | NG | 21 | -23,74 | 50,13 | 6,18 | 14,93 | 0.313 | 7,3 | 6,6 | M |
| 19/05/2016 | NG | 22 | -23,88 | 50,77 | 7,71 | 14,01 | 0.157 | 3,8 | 5,2 | F |
| 19/05/2016 | NG | 24 | -24,51 | 49,97 | 5,84 | 14,89 | 0.345 | 7,4 | 6,8 | M |
| 19/05/2016 | NG | 26 | -24,66 | 44,65 | 6,16 | 12,8 | 0.352 | 9 | 6,7 | M |
| 25/05/2016 | NG | 28 | -24,33 | 51,73 | 5,7 | 15,12 | 0.363 | 7 | 6,4 | M |
| 25/05/2016 | NG | 35 | -24,67 | 50,06 | 5 | 15,11 | 0.355 | 7,6 | 6,6 | M |
| 25/05/2016 | NG | 37 | -24,26 | 51,05 | 5,3 | 15,25 | 0.321 | 8,6 | 6,9 | M |
| 25/05/2016 | NG | 38 | -24,8 | 49,94 | 5,9 | 14,84 | 0.321 | 9 | 6,8 | M |
| 25/05/2016 | NG | 45 | NA | NA | NA | NA | NA | 6,2 | 6 | F |
| 25/05/2016 | NG | 49 | -24,5 | 51,05 | 5,89 | 15,1 | 0.290 | 8,3 | 6,8 | M |

**Table S6:** List of persistent ASVs found in NG.

| ASV | Phylum | Class | Order | Family | Genus | Species |
|---|---|---|---|---|---|---|
| 081a760329ff36deeb00c40b11af8cd6 | Firmicutes | Clostridia | Clostridiales | Veillonellaceae | | |
| aa24cd3ae457aa00fdcbc8ea4ce98e61 | Firmicutes | Clostridia | Clostridiales | Veillonellaceae | | |
| e09c1336c83f322b4c87ce1dfed1ad43 | Firmicutes | Clostridia | Clostridiales | Veillonellaceae | | |
| 036136e8ead714c63c347faa02ba5ff5 | Firmicutes | Clostridia | Clostridiales | Ruminococcaceae | Oscillospira | |
| 03e57e1f7ff88f0f5fe7426886c1e1ce | Firmicutes | Clostridia | Clostridiales | Ruminococcaceae | Oscillospira | |
| 06092ef5739bb73bae8e95de6f863256 | Firmicutes | Clostridia | Clostridiales | Ruminococcaceae | Oscillospira | |
| 07155fe648029be63758ee4f633dad1a | Firmicutes | Clostridia | Clostridiales | Ruminococcaceae | Oscillospira | |
| 087aae6fc14e2f9807532c46c73887ac | Firmicutes | Clostridia | Clostridiales | Ruminococcaceae | Ruminococcus | |
| 21bdcdbf32275700cf3fe244e64d30c1 | Firmicutes | Clostridia | Clostridiales | Ruminococcaceae | | |
| 41de14e523aff5bd65c2983b2dff7cd7 | Firmicutes | Clostridia | Clostridiales | Ruminococcaceae | Oscillospira | |
| 42dd3ae75c45270b0e029f7ecb542f8a | Firmicutes | Clostridia | Clostridiales | Ruminococcaceae | Oscillospira | |
| 46b595fb9d42b1e7a3e7f6c1fef264c8 | Firmicutes | Clostridia | Clostridiales | Ruminococcaceae | | |
| 4894f8be0657aff9ca3292e0cfdab79a | Firmicutes | Clostridia | Clostridiales | Ruminococcaceae | Oscillospira | |
| 4cadfae87ff6ef42975e04007f97c972 | Firmicutes | Clostridia | Clostridiales | Ruminococcaceae | Oscillospira | |
| 542ed6a3edfc1a64db378f18058e24e2 | Firmicutes | Clostridia | Clostridiales | Ruminococcaceae | Oscillospira | |
| 629600e92b9175ac52e765ca4824a8a2 | Firmicutes | Clostridia | Clostridiales | Ruminococcaceae | Oscillospira | |
| 7ca19bde2f95483c8a0823075bbd696b | Firmicutes | Clostridia | Clostridiales | Ruminococcaceae | | |
| 8d1e0a8ca56cbe479ac252b1eaa9bce3 | Firmicutes | Clostridia | Clostridiales | Ruminococcaceae | Oscillospira | |
| 929697b6c6b38ec9296c15a78ef858a1 | Firmicutes | Clostridia | Clostridiales | Ruminococcaceae | Oscillospira | |
| 92b8372223126d525b549ed23ece2de4 | Firmicutes | Clostridia | Clostridiales | Ruminococcaceae | Anaerofilum | |
| 941e0dcdc8442e099bf7fef5b92c758a | Firmicutes | Clostridia | Clostridiales | Ruminococcaceae | Oscillospira | |
| a3ab634bb4c3e266e9fa369353f42a7d | Firmicutes | Clostridia | Clostridiales | Ruminococcaceae | Oscillospira | |
| a6c6f96cc321d7d5522fb2476b79a2d5 | Firmicutes | Clostridia | Clostridiales | Ruminococcaceae | Oscillospira | |
| abb31b5c13e02bd8c596310ce738df08 | Firmicutes | Clostridia | Clostridiales | Ruminococcaceae | Oscillospira | |
| af55d1f16ccf85befde53df479defdb2 | Firmicutes | Clostridia | Clostridiales | Ruminococcaceae | Ruminococcus | |
| bdc2a0fdde88f16f46aae28f0655b773 | Firmicutes | Clostridia | Clostridiales | Ruminococcaceae | | |
| d0678b26f1b1adface9d0c1099e61425 | Firmicutes | Clostridia | Clostridiales | Ruminococcaceae | Oscillospira | |
| d347529956a17a92c99eb4ffa33ee441 | Firmicutes | Clostridia | Clostridiales | Ruminococcaceae | | |
| d3dc955ae8e79b9c9ab0273a6056a708 | Firmicutes | Clostridia | Clostridiales | Ruminococcaceae | Oscillospira | |
| da9659b21943ff56d5dd9adc56cad323 | Firmicutes | Clostridia | Clostridiales | Ruminococcaceae | Oscillospira | |
| e6e2607df619da709c9b9bd9204a5518 | Firmicutes | Clostridia | Clostridiales | Ruminococcaceae | Oscillospira | |
| f4969132c95e0442319f1e004b9fa864 | Firmicutes | Clostridia | Clostridiales | Ruminococcaceae | | |
| f9834a5b3d9f3017206a779b6f26379f | Firmicutes | Clostridia | Clostridiales | Ruminococcaceae | Ruminococcus | |
| b3c3b58775d030e62be1ded907bf3049 | Bacteroidetes | Bacteroidia | Bacteroidales | Rikenellaceae | | |
| c9df468a19fb7488b4b3748062988a49 | Bacteroidetes | Bacteroidia | Bacteroidales | Rikenellaceae | | |
| d4a8a2d25416eb82555ede62dfa6f801 | Bacteroidetes | Bacteroidia | Bacteroidales | Rikenellaceae | | |
| e4e57ce64e6fd2a7b52aae3343de982d | Bacteroidetes | Bacteroidia | Bacteroidales | Rikenellaceae | Rikenella | |
| 093a12421ef526ad04a8eb5fc7787c47 | Bacteroidetes | Bacteroidia | Bacteroidales | Porphyromonadaceae | Parabacteroides | gordonii |
| 1c38217653ccc0b533765fe1141cf5d2 | Bacteroidetes | Bacteroidia | Bacteroidales | Porphyromonadaceae | Parabacteroides | gordonii |
| 44a870333a7202203e144e681e8562e9 | Bacteroidetes | Bacteroidia | Bacteroidales | Porphyromonadaceae | Parabacteroides | |
| 7ae97f9f54cf6fc0d22f6b73b5346c76 | Bacteroidetes | Bacteroidia | Bacteroidales | Porphyromonadaceae | Parabacteroides | gordonii |
| 43d6557b41ba8adda3e6103fb9d0db8a | Firmicutes | Clostridia | Clostridiales | Peptococcaceae | | |
| cf30ae7250a326d515ee06ab165d7686 | Firmicutes | Clostridia | Clostridiales | Peptococcaceae | | |
| 0831db5b5309ed361907c514cbb278d9 | Firmicutes | Clostridia | Clostridiales | Lachnospiraceae | Dorea | |
| 0b056b5af335ddf8c5044b8fce87045c | Firmicutes | Clostridia | Clostridiales | Lachnospiraceae | | |
| 18b87e70f53ce2fbcb612413fa3888a7 | Firmicutes | Clostridia | Clostridiales | Lachnospiraceae | | |
| 1e704442d9676b27608745f8c00c7860 | Firmicutes | Clostridia | Clostridiales | Lachnospiraceae | | |
| 22e5d015e610da7277714a3ebf5000b4 | Firmicutes | Clostridia | Clostridiales | Lachnospiraceae | | |
| 2c9fb000c1c5ce35f2adc2c280e21a4e | Firmicutes | Clostridia | Clostridiales | Lachnospiraceae | Clostridium | |
| 3521cf575046d9249c5d0bcdc5a2eaa1 | Firmicutes | Clostridia | Clostridiales | Lachnospiraceae | | |
| 4b8f9c4141c2ec34339680e981e409cd | Firmicutes | Clostridia | Clostridiales | Lachnospiraceae | | |
| 562654ec58186222c341936e7ff1f82c | Firmicutes | Clostridia | Clostridiales | Lachnospiraceae | | |
| 64ca65803b97525889e7607c1aeaef64 | Firmicutes | Clostridia | Clostridiales | Lachnospiraceae | | |
| 665cf77fe4e20f3d809b716009c45013 | Firmicutes | Clostridia | Clostridiales | Lachnospiraceae | Roseburia | |
| 9496fec2565cbef69e99e590ab010689 | Firmicutes | Clostridia | Clostridiales | Lachnospiraceae | | |
| 995523e90f6cb421b7ac979223605799 | Firmicutes | Clostridia | Clostridiales | Lachnospiraceae | | |
| abab79e1fe980733546408cbf9fdbc0b | Firmicutes | Clostridia | Clostridiales | Lachnospiraceae | Dorea | |
| af849a4dae5a10136f008f623e1fb33b | Firmicutes | Clostridia | Clostridiales | Lachnospiraceae | | |
| b3109319445008804b634d387d46f198 | Firmicutes | Clostridia | Clostridiales | Lachnospiraceae | | |
| b8974b0cd72b2a5079fa3d58eec0cce5 | Firmicutes | Clostridia | Clostridiales | Lachnospiraceae | Coprococcus | |
| bd7fa538242e2349b0d400dc3f2ef77e | Firmicutes | Clostridia | Clostridiales | Lachnospiraceae | [Ruminococcus] | gnavus |
| d0d00eeb4f8ca57fdd13cd1a744fcbde | Firmicutes | Clostridia | Clostridiales | Lachnospiraceae | | |
| d8224527e62d14738a32c74f38e45060 | Firmicutes | Clostridia | Clostridiales | Lachnospiraceae | | |
| e368d24679804a8bea2bab39c2b3a555 | Firmicutes | Clostridia | Clostridiales | Lachnospiraceae | Dorea | |
| eb0ce025f750ad5e9839dc3c10602d78 | Firmicutes | Clostridia | Clostridiales | Lachnospiraceae | | |
| fbc50a8a333a1c1cb09f369f7f0f7bbe | Firmicutes | Clostridia | Clostridiales | Lachnospiraceae | Dorea | |
| ff067bb1428ba11a1250d867cb744298 | Firmicutes | Clostridia | Clostridiales | Lachnospiraceae | | |
| 29bc25fbae29180b45976abd2a3a116d | Firmicutes | Erysipelotrichi | Erysipelotrichales | Erysipelotrichaceae | Anaerorhabdus | furcosa |
| 5ca969e4e49c2150a3ab00f75e94daa0 | Firmicutes | Erysipelotrichi | Erysipelotrichales | Erysipelotrichaceae | | |
| 69d0f6b87c9018be06eef776c8b6f989 | Firmicutes | Erysipelotrichi | Erysipelotrichales | Erysipelotrichaceae | Anaerorhabdus | furcosa |
| 7ef65e7a88f775b4ad3e4ebcb448e331 | Firmicutes | Erysipelotrichi | Erysipelotrichales | Erysipelotrichaceae | Coprobacillus | |
| dc7dc6d036a3e4c7da76d9e33bbf8c78 | Firmicutes | Erysipelotrichi | Erysipelotrichales | Erysipelotrichaceae | | |
| e03f9292fb2b629dac6e6bd40704b2de | Firmicutes | Erysipelotrichi | Erysipelotrichales | Erysipelotrichaceae | Clostridium | ramosum |
| ecb83c1fd0d885366790df39ff61e472 | Firmicutes | Erysipelotrichi | Erysipelotrichales | Erysipelotrichaceae | | |
| f5ccb8e6959c242ae9753b5d77edfec4 | Firmicutes | Erysipelotrichi | Erysipelotrichales | Erysipelotrichaceae | | |
| 9ddbb9f3235a4d49679636c03acd5186 | Proteobacteria | Deltaproteobacteria | Desulfovibrionales | Desulfovibrionaceae | Desulfovibrio | |
| ad82b6c42a951f8c5940494b1c4c1877 | Proteobacteria | Deltaproteobacteria | Desulfovibrionales | Desulfovibrionaceae | | |
| eb26eefda4af0aeca7432a7561cc8d36 | Actinobacteria | Coriobacteriia | Coriobacteriales | Coriobacteriaceae | | |
| 16581f1341159085917e0b0c14834b12 | Firmicutes | Clostridia | Clostridiales | Christensenellaceae | | |
| 03d8e905c3acd9038bbea399fc9afea4 | Bacteroidetes | Bacteroidia | Bacteroidales | Bacteroidaceae | Bacteroides | |
| 0c66d5e224490b5fe7b5f9ee9ab6166e | Bacteroidetes | Bacteroidia | Bacteroidales | Bacteroidaceae | Bacteroides | |
| 28577d0c3cdd6e00da06f83a3bb164f7 | Bacteroidetes | Bacteroidia | Bacteroidales | Bacteroidaceae | Bacteroides | |
| 411d5a652e0c5bed13f0e0bf184d918b | Bacteroidetes | Bacteroidia | Bacteroidales | Bacteroidaceae | Bacteroides | |
| 428829cca0363b191c80f561f314f3ae | Bacteroidetes | Bacteroidia | Bacteroidales | Bacteroidaceae | Bacteroides | |
| 500f586b82a340be51437db122efe58f | Bacteroidetes | Bacteroidia | Bacteroidales | Bacteroidaceae | Bacteroides | |
| 615921a2952ff64b4b39e2630a18d479 | Bacteroidetes | Bacteroidia | Bacteroidales | Bacteroidaceae | Bacteroides | |
| c5e2344e6a032cd4d7db4c73bbfafd32 | Bacteroidetes | Bacteroidia | Bacteroidales | Bacteroidaceae | Bacteroides | |
| edeb07eb88e61abf3b088ed158b281ed | Bacteroidetes | Bacteroidia | Bacteroidales | Bacteroidaceae | Bacteroides | |
| af6834e6719155b144e4649c1ae805fe | Tenericutes | Mollicutes | Anaeroplasmatales | Anaeroplasmataceae | Anaeroplasma | |
| db2b5228af3edc1ec960afff42527116 | Bacteroidetes | Bacteroidia | Bacteroidales | [Odoribacteraceae] | Odoribacter | |
| 5b776dbaa20108557e01c959eb7c1bd1 | Firmicutes | Clostridia | Clostridiales | [Mogibacteriaceae] | | |
| 2aaae3842fd8be509b775740a4ec787e | Firmicutes | Clostridia | Clostridiales | | | |
| 2c4aee95cb2e0deb355f1e0afb3c7626 | Firmicutes | Clostridia | Clostridiales | | | |
| 4989fa9089225629c4007e53d3a3714f | Firmicutes | Clostridia | Clostridiales | | | |
| 6bb53cd3333a71f68d7ecabbabd58418 | Firmicutes | Clostridia | Clostridiales | | | |
| 76e5dafdee60ea522da07bccf36758c7 | Firmicutes | Clostridia | Clostridiales | | | |
| 86d33f33b14662a6a4e033d0b540278a | Firmicutes | Clostridia | Clostridiales | | | |
| 9bdbdf13a20c41052da53de3b4436ab8 | Firmicutes | Clostridia | Clostridiales | | | |
| a4b929d25334c34eb18fca9a5669ed52 | Firmicutes | Clostridia | Clostridiales | | | |
| d4ede4f69b87950cfc2a677dd7fc0d15 | Firmicutes | Clostridia | Clostridiales | | | |
| e7782b6827bb042621b9d8271e8a6660 | Firmicutes | Clostridia | Clostridiales | | | |
| e68f6ba852e82281e53b2bb128f04731 | Proteobacteria | Alphaproteobacteria | Rickettsiales | | | |

**Table S7:** List of persistent ASVs found in NM.

| ASV | Phylum | Class | Order | Family | Genus | Species |
|---|---|---|---|---|---|---|
| ec0d6515d924c1f0d0e807d9d9038d88 | Lentisphaerae | [Lentisphaeria] | Victivallales | Victivallaceae | | |
| aa24cd3ae457aa00fdcbc8ea4ce98e61 | Firmicutes | Clostridia | Clostridiales | Veillonellaceae | | |
| e09c1336c83f322b4c87ce1dfed1ad43 | Firmicutes | Clostridia | Clostridiales | Veillonellaceae | | |
| dc1d000d2ad856b1efbd9af1ce0c4568 | Bacteroidetes | Bacteroidia | Bacteroidales | S24-7 | | |
| 036136e8ead714c63c347faa02ba5ff5 | Firmicutes | Clostridia | Clostridiales | Ruminococcaceae | Oscillospira | |
| 03e57e1f7ff88f0f5fe7426886c1e1ce | Firmicutes | Clostridia | Clostridiales | Ruminococcaceae | Oscillospira | |
| 0602d20727fc87d1cd12850e5aa620b0 | Firmicutes | Clostridia | Clostridiales | Ruminococcaceae | | |
| 06092ef5739bb73bae8e95de6f863256 | Firmicutes | Clostridia | Clostridiales | Ruminococcaceae | Oscillospira | |
| 07155fe648029be63758ee4f633dad1a | Firmicutes | Clostridia | Clostridiales | Ruminococcaceae | Oscillospira | |
| 0878a69626dbc7078de8049603eb9bdd | Firmicutes | Clostridia | Clostridiales | Ruminococcaceae | Oscillospira | |
| 18697e8240afc0017f20fde5136a1490 | Firmicutes | Clostridia | Clostridiales | Ruminococcaceae | Oscillospira | |
| 1f28b3e656896666ff6b493769d89aae | Firmicutes | Clostridia | Clostridiales | Ruminococcaceae | | |
| 25a96134e15b97c172c162e860e8a326 | Firmicutes | Clostridia | Clostridiales | Ruminococcaceae | | |
| 3348c196d1bde3961288015f9a0deb38 | Firmicutes | Clostridia | Clostridiales | Ruminococcaceae | | |
| 3c80993b2cbd964f651ccedf74339b54 | Firmicutes | Clostridia | Clostridiales | Ruminococcaceae | | |
| 41de14e523aff5bd65c2983b2dff7cd7 | Firmicutes | Clostridia | Clostridiales | Ruminococcaceae | Oscillospira | |
| 42dd3ae75c45270b0e029f7ecb542f8a | Firmicutes | Clostridia | Clostridiales | Ruminococcaceae | Oscillospira | |
| 4559248c74d5709108c66a1053efa742 | Firmicutes | Clostridia | Clostridiales | Ruminococcaceae | Oscillospira | |
| 46b595fb9d42b1e7a3e7f6c1fef264c8 | Firmicutes | Clostridia | Clostridiales | Ruminococcaceae | | |
| 4894f8be0657aff9ca3292e0cfdab79a | Firmicutes | Clostridia | Clostridiales | Ruminococcaceae | Oscillospira | |
| 4cadfae87ff6ef42975e04007f97c972 | Firmicutes | Clostridia | Clostridiales | Ruminococcaceae | Oscillospira | |
| 629600e92b9175ac52e765ca4824a8a2 | Firmicutes | Clostridia | Clostridiales | Ruminococcaceae | Oscillospira | |
| 7ca19bde2f95483c8a0823075bbd696b | Firmicutes | Clostridia | Clostridiales | Ruminococcaceae | | |
| 880de7650bc43946b1eeaa9eb84c9afa | Firmicutes | Clostridia | Clostridiales | Ruminococcaceae | Butyricicoccus | pullicaecorum |
| 8d1e0a8ca56cbe479ac252b1eaa9bce3 | Firmicutes | Clostridia | Clostridiales | Ruminococcaceae | Oscillospira | |
| 929697b6c6b38ec9296c15a78ef858a1 | Firmicutes | Clostridia | Clostridiales | Ruminococcaceae | Oscillospira | |
| 92b8372223126d525b549ed23ece2de4 | Firmicutes | Clostridia | Clostridiales | Ruminococcaceae | Anaerofilum | |
| 941e0dcdc8442e099bf7fef5b92c758a | Firmicutes | Clostridia | Clostridiales | Ruminococcaceae | Oscillospira | |
| 9c849abf0fd1983fb81ee109c20b4f87 | Firmicutes | Clostridia | Clostridiales | Ruminococcaceae | Oscillospira | |
| a3ab634bb4c3e266e9fa369353f42a7d | Firmicutes | Clostridia | Clostridiales | Ruminococcaceae | Oscillospira | |
| a4b08afb41fd7c4ccf3c3365a98058f8 | Firmicutes | Clostridia | Clostridiales | Ruminococcaceae | Oscillospira | |
| ab3c094e8541a731e154432e2c0285c4 | Firmicutes | Clostridia | Clostridiales | Ruminococcaceae | Oscillospira | |
| abb31b5c13e02bd8c596310ce738df08 | Firmicutes | Clostridia | Clostridiales | Ruminococcaceae | Oscillospira | |
| af55d1f16ccf85befde53df479defdb2 | Firmicutes | Clostridia | Clostridiales | Ruminococcaceae | Ruminococcus | |
| b351470ecce44478b75ff6ccefc322e6 | Firmicutes | Clostridia | Clostridiales | Ruminococcaceae | Oscillospira | |
| bdc2a0fdde88f16f46aae28f0655b773 | Firmicutes | Clostridia | Clostridiales | Ruminococcaceae | | |
| c6555f27c699d01fa66b78a103559778 | Firmicutes | Clostridia | Clostridiales | Ruminococcaceae | | |
| d0678b26f1b1adface9d0c1099e61425 | Firmicutes | Clostridia | Clostridiales | Ruminococcaceae | Oscillospira | |
| d347529956a17a92c99eb4ffa33ee441 | Firmicutes | Clostridia | Clostridiales | Ruminococcaceae | | |
| d3dc955ae8e79b9c9ab0273a6056a708 | Firmicutes | Clostridia | Clostridiales | Ruminococcaceae | Oscillospira | |
| da9659b21943ff56d5dd9adc56cad323 | Firmicutes | Clostridia | Clostridiales | Ruminococcaceae | Oscillospira | |
| e6e2607df619da709c9b9bd9204a5518 | Firmicutes | Clostridia | Clostridiales | Ruminococcaceae | Oscillospira | |
| eebd291873d6fe80792d922968e729d8 | Firmicutes | Clostridia | Clostridiales | Ruminococcaceae | Ruminococcus | |
| f4969132c95e0442319f1e004b9fa864 | Firmicutes | Clostridia | Clostridiales | Ruminococcaceae | | |
| f9834a5b3d9f3017206a779b6f26379f | Firmicutes | Clostridia | Clostridiales | Ruminococcaceae | Ruminococcus | |
| fbce01f7cc604374247d1f8675bb29f2 | Firmicutes | Clostridia | Clostridiales | Ruminococcaceae | Oscillospira | |
| 00eea6adb849d7d173c19090cbdbfcfc | Bacteroidetes | Bacteroidia | Bacteroidales | Rikenellaceae | | |
| 5c43565006d70751ace565ef8b6d819c | Bacteroidetes | Bacteroidia | Bacteroidales | Rikenellaceae | Alistipes | indistinctus |
| 63380953905c02558cff39c2d1ce9d10 | Bacteroidetes | Bacteroidia | Bacteroidales | Rikenellaceae | | |
| a7a05bb9360908c2fc537444128aaa9c | Bacteroidetes | Bacteroidia | Bacteroidales | Rikenellaceae | | |
| b2ffca3d6d8c8ac2d2d938e1b5bb915f | Bacteroidetes | Bacteroidia | Bacteroidales | Rikenellaceae | | |
| c9df468a19fb7488b4b3748062988a49 | Bacteroidetes | Bacteroidia | Bacteroidales | Rikenellaceae | | |
| d8accc5224fcea846f8d2647b1bd3ae9 | Bacteroidetes | Bacteroidia | Bacteroidales | Rikenellaceae | | |
| e4e57ce64e6fd2a7b52aae3343de982d | Bacteroidetes | Bacteroidia | Bacteroidales | Rikenellaceae | Rikenella | |
| 093a12421ef526ad04a8eb5fc7787c47 | Bacteroidetes | Bacteroidia | Bacteroidales | Porphyromonadaceae | Parabacteroides | gordonii |
| 1c38217653ccc0b533765fe1141cf5d2 | Bacteroidetes | Bacteroidia | Bacteroidales | Porphyromonadaceae | Parabacteroides | gordonii |
| 342ac5cf320f718ca71dd4b823ffb9ed | Bacteroidetes | Bacteroidia | Bacteroidales | Porphyromonadaceae | Porphyromonas | |
| 44a870333a7202203e144e681e8562e9 | Bacteroidetes | Bacteroidia | Bacteroidales | Porphyromonadaceae | Parabacteroides | |
| 7ae97f9f54cf6fc0d22f6b73b5346c76 | Bacteroidetes | Bacteroidia | Bacteroidales | Porphyromonadaceae | Parabacteroides | gordonii |
| cf30ae7250a326d515ee06ab165d7686 | Firmicutes | Clostridia | Clostridiales | Peptococcaceae | | |
| 0b056b5af335ddf8c5044b8fce87045c | Firmicutes | Clostridia | Clostridiales | Lachnospiraceae | | |
| 18b87e70f53ce2fbcb612413fa3888a7 | Firmicutes | Clostridia | Clostridiales | Lachnospiraceae | | |
| 1c2204512a01b3e0708db9c91dd52f8b | Firmicutes | Clostridia | Clostridiales | Lachnospiraceae | | |
| 1e704442d9676b27608745f8c00c7860 | Firmicutes | Clostridia | Clostridiales | Lachnospiraceae | | |
| 22e5d015e610da7277714a3ebf5000b4 | Firmicutes | Clostridia | Clostridiales | Lachnospiraceae | | |
| 28fb140d46cf167a3f1976eb4d8855a0 | Firmicutes | Clostridia | Clostridiales | Lachnospiraceae | | |
| 2c9fb000c1c5ce35f2adc2c280e21a4e | Firmicutes | Clostridia | Clostridiales | Lachnospiraceae | Clostridium | |
| 369e064350f5b719ba67b7db0a257b5b | Firmicutes | Clostridia | Clostridiales | Lachnospiraceae | | |
| 4b8f9c4141c2ec34339680e981e409cd | Firmicutes | Clostridia | Clostridiales | Lachnospiraceae | | |
| 64ca65803b97525889e7607c1aeaef64 | Firmicutes | Clostridia | Clostridiales | Lachnospiraceae | | |
| 665cf77fe4e20f3d809b716009c45013 | Firmicutes | Clostridia | Clostridiales | Lachnospiraceae | Roseburia | |
| 701a62ed34ac3d0bb343337fffdecea3 | Firmicutes | Clostridia | Clostridiales | Lachnospiraceae | | |
| 8771b6279e16fc8aa2e2ba2646b03e33 | Firmicutes | Clostridia | Clostridiales | Lachnospiraceae | | |
| 9496fec2565cbef69e99e590ab010689 | Firmicutes | Clostridia | Clostridiales | Lachnospiraceae | | |
| 995523e90f6cb421b7ac979223605799 | Firmicutes | Clostridia | Clostridiales | Lachnospiraceae | | |
| 9f6619e957bc0557f07c6580ee325b9f | Firmicutes | Clostridia | Clostridiales | Lachnospiraceae | | |
| a7170ea6f21f65769bbcb0131625f4f3 | Firmicutes | Clostridia | Clostridiales | Lachnospiraceae | Blautia | producta |
| abab79e1fe980733546408cbf9fdbc0b | Firmicutes | Clostridia | Clostridiales | Lachnospiraceae | Dorea | |
| b3109319445008804b634d387d46f198 | Firmicutes | Clostridia | Clostridiales | Lachnospiraceae | | |
| b8974b0cd72b2a5079fa3d58eec0cce5 | Firmicutes | Clostridia | Clostridiales | Lachnospiraceae | Coprococcus | |
| bd7fa538242e2349b0d400dc3f2ef77e | Firmicutes | Clostridia | Clostridiales | Lachnospiraceae | [Ruminococcus] | gnavus |
| d0d00eeb4f8ca57fdd13cd1a744fcbde | Firmicutes | Clostridia | Clostridiales | Lachnospiraceae | | |
| dbfcf5d7bf1ec461d40c0befebf1835d | Firmicutes | Clostridia | Clostridiales | Lachnospiraceae | Coprococcus | |
| deb69b52e59acc99e6941d9c97f469f0 | Firmicutes | Clostridia | Clostridiales | Lachnospiraceae | | |
| e368d24679804a8bea2bab39c2b3a555 | Firmicutes | Clostridia | Clostridiales | Lachnospiraceae | Dorea | |
| eb0ce025f750ad5e9839dc3c10602d78 | Firmicutes | Clostridia | Clostridiales | Lachnospiraceae | | |
| fbc50a8a333a1c1cb09f369f7f0f7bbe | Firmicutes | Clostridia | Clostridiales | Lachnospiraceae | Dorea | |
| ff067bb1428ba11a1250d867cb744298 | Firmicutes | Clostridia | Clostridiales | Lachnospiraceae | | |
| aa98dc927a4f154d35a001085ca9bcce | Proteobacteria | Epsilonproteobacteria | Campylobacterales | Helicobacteraceae | Flexispira | |
| 6e81c93a891bd6660cee4be66e70aef4 | Firmicutes | Clostridia | Clostridiales | Eubacteriaceae | Pseudoramibacter_Eubacterium | |
| 5ca969e4e49c2150a3ab00f75e94daa0 | Firmicutes | Erysipelotrichi | Erysipelotrichales | Erysipelotrichaceae | | |
| 7ef65e7a88f775b4ad3e4ebcb448e331 | Firmicutes | Erysipelotrichi | Erysipelotrichales | Erysipelotrichaceae | Coprobacillus | |
| 85ea3ab37ef0ba4230d5e18ff5eebcb4 | Firmicutes | Erysipelotrichi | Erysipelotrichales | Erysipelotrichaceae | Anaerorhabdus | furcosa |
| 8ba1003dee1708fc6fd6246977a42bc4 | Firmicutes | Erysipelotrichi | Erysipelotrichales | Erysipelotrichaceae | | |
| c8db65cd923eb078d3082e7f6ffe7ddc | Firmicutes | Erysipelotrichi | Erysipelotrichales | Erysipelotrichaceae | | |
| dc7dc6d036a3e4c7da76d9e33bbf8c78 | Firmicutes | Erysipelotrichi | Erysipelotrichales | Erysipelotrichaceae | | |
| e03f9292fb2b629dac6e6bd40704b2de | Firmicutes | Erysipelotrichi | Erysipelotrichales | Erysipelotrichaceae | Clostridium | ramosum |
| ecb83c1fd0d885366790df39ff61e472 | Firmicutes | Erysipelotrichi | Erysipelotrichales | Erysipelotrichaceae | | |
| f11e992935bfb9edcd1c151eec52970a | Firmicutes | Erysipelotrichi | Erysipelotrichales | Erysipelotrichaceae | | |
| f5ccb8e6959c242ae9753b5d77edfec4 | Firmicutes | Erysipelotrichi | Erysipelotrichales | Erysipelotrichaceae | | |
| 28c9e399296808ddd37ea2feaf092d12 | Proteobacteria | Deltaproteobacteria | Desulfovibrionales | Desulfovibrionaceae | Bilophila | |
| 7c2d30a0057a38e19e314bf6c729514b | Proteobacteria | Deltaproteobacteria | Desulfovibrionales | Desulfovibrionaceae | | |
| 9ddbb9f3235a4d49679636c03acd5186 | Proteobacteria | Deltaproteobacteria | Desulfovibrionales | Desulfovibrionaceae | Desulfovibrio | |
| fbdf2b1282fe99b4c707f69887a30aa6 | Firmicutes | Clostridia | Clostridiales | Dehalobacteriaceae | Dehalobacterium | |
| 1c95c7c381d22117c65fa3d804000ce4 | Actinobacteria | Coriobacteriia | Coriobacteriales | Coriobacteriaceae | Paraeggerthella | hongkongensis |
| 957bc4f51f13dcf1c85785d3d60e4777 | Actinobacteria | Coriobacteriia | Coriobacteriales | Coriobacteriaceae | | |
| d186abf5e3f0d65a7dc7a866a6458b3d | Actinobacteria | Coriobacteriia | Coriobacteriales | Coriobacteriaceae | Adlercreutzia | |
| eb26eefda4af0aeca7432a7561cc8d36 | Actinobacteria | Coriobacteriia | Coriobacteriales | Coriobacteriaceae | | |
| ebc6118f1e70c631be2fb9094dbf00dc | Actinobacteria | Coriobacteriia | Coriobacteriales | Coriobacteriaceae | Enterococcus | casseliflavus |
| 16581f1341159085917e0b0c14834b12 | Firmicutes | Clostridia | Clostridiales | Christensenellaceae | | |
| 03d8e905c3acd9038bbea399fc9afea4 | Bacteroidetes | Bacteroidia | Bacteroidales | Bacteroidaceae | Bacteroides | |
| 072cd9606ef2440fbb7dd3f585d38dbc | Bacteroidetes | Bacteroidia | Bacteroidales | Bacteroidaceae | Bacteroides | |
| 0c66d5e224490b5fe7b5f9ee9ab6166e | Bacteroidetes | Bacteroidia | Bacteroidales | Bacteroidaceae | Bacteroides | |
| 1210c67494f234869c42b7c98d2ec758 | Bacteroidetes | Bacteroidia | Bacteroidales | Bacteroidaceae | Bacteroides | |
| 28577d0c3cdd6e00da06f83a3bb164f7 | Bacteroidetes | Bacteroidia | Bacteroidales | Bacteroidaceae | Bacteroides | |
| 3ae938ca339d3b75e9b1180eabe6f47d | Bacteroidetes | Bacteroidia | Bacteroidales | Bacteroidaceae | Bacteroides | |
| 3bb7551ec97a9bacd2eb126b8521666a | Bacteroidetes | Bacteroidia | Bacteroidales | Bacteroidaceae | Bacteroides | |
| 411d5a652e0c5bed13f0e0bf184d918b | Bacteroidetes | Bacteroidia | Bacteroidales | Bacteroidaceae | Bacteroides | |
| 428829cca0363b191c80f561f314f3ae | Bacteroidetes | Bacteroidia | Bacteroidales | Bacteroidaceae | Bacteroides | |
| 4c31b9f379fb5ef4845299340e4ecace | Bacteroidetes | Bacteroidia | Bacteroidales | Bacteroidaceae | Bacteroides | |
| 5f65c177995eee5dea4d04aef2753009 | Bacteroidetes | Bacteroidia | Bacteroidales | Bacteroidaceae | Bacteroides | |
| 615921a2952ff64b4b39e2630a18d479 | Bacteroidetes | Bacteroidia | Bacteroidales | Bacteroidaceae | Bacteroides | |
| c5e2344e6a032cd4d7db4c73bbfafd32 | Bacteroidetes | Bacteroidia | Bacteroidales | Bacteroidaceae | Bacteroides | |
| af6834e6719155b144e4649c1ae805fe | Tenericutes | Mollicutes | Anaeroplasmatales | Anaeroplasmataceae | Anaeroplasma | |
| ec3ef6d3fcb45d763873ac9417e91430 | Tenericutes | Mollicutes | Anaeroplasmatales | Anaeroplasmataceae | | |
| 9f2c0bcef5b3acc9f882e18a5f51e305 | Bacteroidetes | Bacteroidia | Bacteroidales | [Odoribacteraceae] | Odoribacter | |
| b5c97bb118b54e8fc1afbba09dcb8d82 | Bacteroidetes | Bacteroidia | Bacteroidales | [Odoribacteraceae] | Odoribacter | |
| ef5a2ca667bd274b643f1e1d4458a955 | Bacteroidetes | Bacteroidia | Bacteroidales | [Odoribacteraceae] | Odoribacter | |
| 14fbfa9a624e36f83d008152b8ad120e | Firmicutes | Clostridia | Clostridiales | [Mogibacteriaceae] | Anaerovorax | |
| 4b12e7a3b19303f42b4db94503cba1bd | Firmicutes | Clostridia | Clostridiales | [Mogibacteriaceae] | | |
| 5b776dbaa20108557e01c959eb7c1bd1 | Firmicutes | Clostridia | Clostridiales | [Mogibacteriaceae] | | |
| 7b23d4c9eda60f616d6b4adc7b21dcbf | Firmicutes | Clostridia | Clostridiales | [Mogibacteriaceae] | | |
| 8dbc4df00378a18b2c8d1d7debdd6300 | Firmicutes | Clostridia | Clostridiales | [Mogibacteriaceae] | | |
| 67a55f8f72d812e5dcf6e03a405efb56 | Tenericutes | Mollicutes | RF39 | | | |
| 262bcf8e78a28ce7adf83ffe3cf66588 | Proteobacteria | Alphaproteobacteria | RF32 | | | |
| e68f6ba852e82281e53b2bb128f04731 | Proteobacteria | Alphaproteobacteria | Rickettsiales | | | |
| 263b81aff509464acfad59da91fa90f8 | Firmicutes | Clostridia | Clostridiales | | | |
| 2aaae3842fd8be509b775740a4ec787e | Firmicutes | Clostridia | Clostridiales | | | |
| 2c4aee95cb2e0deb355f1e0afb3c7626 | Firmicutes | Clostridia | Clostridiales | | | |
| 4989fa9089225629c4007e53d3a3714f | Firmicutes | Clostridia | Clostridiales | | | |
| 6bb53cd3333a71f68d7ecabbabd58418 | Firmicutes | Clostridia | Clostridiales | | | |
| 7ef1f0e4c8d4afe5d15893fa737a4ee1 | Firmicutes | Clostridia | Clostridiales | | | |
| 86d33f33b14662a6a4e033d0b540278a | Firmicutes | Clostridia | Clostridiales | | | |
| 8d9db6bf4f91eb03410f4b65e5992d1a | Firmicutes | Clostridia | Clostridiales | | | |
| 9bdbdf13a20c41052da53de3b4436ab8 | Firmicutes | Clostridia | Clostridiales | | | |
| a4b929d25334c34eb18fca9a5669ed52 | Firmicutes | Clostridia | Clostridiales | | | |
| afd6c9a08f045d12b64b439021f3337d | Firmicutes | Clostridia | Clostridiales | | | |
| d4ede4f69b87950cfc2a677dd7fc0d15 | Firmicutes | Clostridia | Clostridiales | | | |
| dbcc1b08e37598d13406803bf6825606 | Firmicutes | Clostridia | Clostridiales | | | |
| e7782b6827bb042621b9d8271e8a6660 | Firmicutes | Clostridia | Clostridiales | | | |
| fefeae219591f8b87c12533e30566824 | Firmicutes | Clostridia | Clostridiales | | | |

**Table S8:** List of ASVs that signifcantly discriminated between seasons within single islets (NG and NM) according to both indval and LEfSe analyses

| | LEFSE | | | | INDVAL | Indval | | | Relfreq | | Relabund | | Taxonomy | | | | | |
|---|---|---|---|---|---|---|---|---|---|---|---|---|---|---|---|---|---|---|
| **ASV** | **log_high_class** | **Class** | **LDA** | **p.value** | **Islet** | **Autumn** | **Spring** | **pval** | **Autumn** | **Spring** | **Autumn** | **Spring** | **Phylum** | **Class** | **Order** | **Family** | **Genus** | **Species** |
| 605ab0359371a294f8108c26d4c4230a | 3,450 | Spring | 3,045 | 0,000 | NG | 0,03 | 0,73 | 0 | 0,24 | 0,83 | 0,12 | 0,88 | Firmicutes | Clostridia | Clostridiales | Ruminococcaceae | | |
| 7ef1f0e4c8d4afe5d15893fa737a4ee1 | 4,238 | Spring | 3,692 | 0,001 | NG | 0,07 | 0,59 | 0 | 0,24 | 0,83 | 0,29 | 0,71 | Firmicutes | Clostridia | Clostridiales | | | |
| 866d02483b53b5ecc63102c27534ceef | 3,461 | Spring | 3,139 | 0,000 | NG | 0,01 | 0,75 | 0 | 0,19 | 0,78 | 0,03 | 0,97 | Firmicutes | Clostridia | Clostridiales | Ruminococcaceae | Ruminococcus | |
| 8771b6279e16fc8aa2e2ba2646b03e33 | 3,904 | Spring | 3,597 | 0,001 | NG | 0,02 | 0,8 | 0 | 0,48 | 0,83 | 0,04 | 0,96 | Firmicutes | Clostridia | Clostridiales | Lachnospiraceae | | |
| 95ea322f33388879b977a9c7e71c584e | 4,042 | Spring | 3,762 | 0,000 | NG | 0 | 0,61 | 0 | 0 | 0,61 | 0 | 1 | Tenericutes | Mollicutes | Anaeroplasmatales | Anaeroplasmataceae | | |
| d8accc5224fcea846f8d2647b1bd3ae9 | 3,509 | Spring | 3,097 | 0,001 | NG | 0,04 | 0,57 | 0 | 0,29 | 0,67 | 0,14 | 0,86 | Bacteroidetes | Bacteroidia | Bacteroidales | Rikenellaceae | | |
| edbd029a7d064892a5ca5a1c4fbcb09b | 3,937 | Spring | 3,577 | 0,000 | NG | 0,03 | 0,76 | 0 | 0,33 | 0,83 | 0,09 | 0,91 | Bacteroidetes | Bacteroidia | Bacteroidales | Rikenellaceae | | |
| ef5a2ca667bd274b643f1e1d4458a955 | 4,142 | Spring | 3,711 | 0,000 | NG | 0,05 | 0,73 | 0 | 0,29 | 0,89 | 0,18 | 0,82 | Bacteroidetes | Bacteroidia | Bacteroidales | [Odoribacteraceae] | Odoribacter | |
| 1e704442d9676b27608745f8c00c7860 | 3,953 | Fall | 3,567 | 0,000 | NG | 0,84 | 0,1 | 0 | 1 | 0,61 | 0,84 | 0,16 | Firmicutes | Clostridia | Clostridiales | Lachnospiraceae | | |
| 2c4aee95cb2e0deb355f1e0afb3c7626 | 4,310 | Fall | 3,963 | 0,001 | NG | 0,88 | 0,09 | 0 | 1 | 0,78 | 0,88 | 0,12 | Firmicutes | Clostridia | Clostridiales | | | |
| 4b8f9c4141c2ec34339680e981e409cd | 3,606 | Fall | 3,008 | 0,002 | NG | 0,77 | 0,22 | 0 | 1 | 0,94 | 0,77 | 0,23 | Firmicutes | Clostridia | Clostridiales | Lachnospiraceae | | |
| 4cadfae87ff6ef42975e04007f97c972 | 3,947 | Fall | 3,451 | 0,000 | NG | 0,73 | 0,27 | 0 | 1 | 1 | 0,73 | 0,27 | Firmicutes | Clostridia | Clostridiales | Ruminococcaceae | Oscillospira | |
| 59e3f9a4091eac22ffe16b2167af6bfa | 3,913 | Fall | 3,586 | 0,000 | NG | 0,86 | 0,05 | 0 | 0,95 | 0,56 | 0,9 | 0,1 | Firmicutes | Clostridia | Clostridiales | Lachnospiraceae | | |
| 6b10014e0bf7c76d6096fba0e5055f3a | 3,564 | Fall | 3,200 | 0,003 | NG | 0,66 | 0,02 | 0 | 0,71 | 0,22 | 0,92 | 0,08 | Bacteroidetes | Bacteroidia | Bacteroidales | Porphyromonadaceae | Parabacteroides | Parabacteroides gordonii |
| 995523e90f6cb421b7ac979223605799 | 4,243 | Fall | 3,874 | 0,000 | NG | 0,84 | 0,08 | 0 | 0,95 | 0,72 | 0,88 | 0,12 | Firmicutes | Clostridia | Clostridiales | Lachnospiraceae | | |
| af849a4dae5a10136f008f623e1fb33b | 3,455 | Fall | 3,082 | 0,006 | NG | 0,88 | 0,05 | 0 | 0,95 | 0,67 | 0,93 | 0,07 | Firmicutes | Clostridia | Clostridiales | Lachnospiraceae | | |
| b0d703f37798eb4720000dbb35113165 | 3,556 | Fall | 3,145 | 0,002 | NG | 0,72 | 0,1 | 0 | 0,9 | 0,5 | 0,79 | 0,21 | Firmicutes | Clostridia | Clostridiales | Lachnospiraceae | | |
| deb69b52e59acc99e6941d9c97f469f0 | 3,790 | Fall | 3,400 | 0,000 | NG | 0,82 | 0,06 | 0 | 0,9 | 0,61 | 0,9 | 0,1 | Firmicutes | Clostridia | Clostridiales | Lachnospiraceae | | |
| f11e992935bfb9edcd1c151eec52970a | 4,043 | Fall | 3,699 | 0,000 | NG | 0,9 | 0,07 | 0 | 1 | 0,72 | 0,9 | 0,1 | Firmicutes | Erysipelotrichi | Erysipelotrichales | Erysipelotrichaceae | | |
| f5ccb8e6959c242ae9753b5d77edfec4 | 4,043 | Fall | 3,619 | 0,000 | NG | 0,83 | 0,17 | 0 | 1 | 1 | 0,83 | 0,17 | Firmicutes | Erysipelotrichi | Erysipelotrichales | Erysipelotrichaceae | | |
| 0070c81cb25fd9a1dec14534c39aef0a | 3,531 | Spring | 3,125 | 0,006 | NM | 0,1 | 0,72 | 0 | 0,55 | 0,88 | 0,17 | 0,83 | Firmicutes | Clostridia | Clostridiales | Lachnospiraceae | Clostridium | |
| 38ce9310bd47517f19bf91451f371907 | 3,416 | Spring | 3,092 | 0,001 | NM | 0,03 | 0,69 | 0 | 0,41 | 0,75 | 0,07 | 0,93 | Bacteroidetes | Bacteroidia | Bacteroidales | [Odoribacteraceae] | Odoribacter | |
| a058e645f47175cd6d59b24705cc73dd | 3,697 | Spring | 3,372 | 0,000 | NM | 0,01 | 0,72 | 0 | 0,31 | 0,75 | 0,04 | 0,96 | Bacteroidetes | Bacteroidia | Bacteroidales | [Odoribacteraceae] | Odoribacter | |
| d0d00eeb4f8ca57fdd13cd1a744fcbde | 3,979 | Spring | 3,597 | 0,000 | NM | 0,05 | 0,91 | 0 | 0,59 | 1 | 0,09 | 0,91 | Firmicutes | Clostridia | Clostridiales | Lachnospiraceae | | |
| d96e00dd528c1e3a12f8103877bc6f07 | 3,499 | Spring | 3,063 | 0,002 | NM | 0,1 | 0,69 | 0,01 | 0,48 | 0,88 | 0,22 | 0,78 | Firmicutes | Clostridia | Clostridiales | Lachnospiraceae | | |
| 1e704442d9676b27608745f8c00c7860 | 3,958 | Fall | 3,556 | 0,003 | NM | 0,69 | 0,12 | 0,01 | 0,83 | 0,69 | 0,83 | 0,17 | Firmicutes | Clostridia | Clostridiales | Lachnospiraceae | | |
| 21bdcdbf32275700cf3fe244e64d30c1 | 3,671 | Fall | 3,293 | 0,000 | NM | 0,72 | 0,11 | 0 | 0,9 | 0,56 | 0,8 | 0,2 | Firmicutes | Clostridia | Clostridiales | Ruminococcaceae | | |
| 2c4aee95cb2e0deb355f1e0afb3c7626 | 4,162 | Fall | 3,798 | 0,002 | NM | 0,85 | 0,11 | 0 | 0,97 | 0,94 | 0,88 | 0,12 | Firmicutes | Clostridia | Clostridiales | | | |
| 629600e92b9175ac52e765ca4824a8a2 | 3,842 | Fall | 3,508 | 0,000 | NM | 0,95 | 0,04 | 0 | 1 | 0,75 | 0,95 | 0,05 | Firmicutes | Clostridia | Clostridiales | Ruminococcaceae | Oscillospira | |
| 76e5dafdee60ea522da07bccf36758c7 | 3,947 | Fall | 3,568 | 0,000 | NM | 0,75 | 0,06 | 0,01 | 0,86 | 0,44 | 0,87 | 0,13 | Firmicutes | Clostridia | Clostridiales | | | |
| 929697b6c6b38ec9296c15a78ef858a1 | 3,690 | Fall | 3,316 | 0,001 | NM | 0,8 | 0,17 | 0 | 1 | 0,88 | 0,8 | 0,2 | Firmicutes | Clostridia | Clostridiales | Ruminococcaceae | Oscillospira | |
