## Supporting Information, Figure S1 for "Insular holobionts: persistence and seasonal plasticity of the Balearic wall lizard (*Podarcis lilfordi*) gut microbiota"

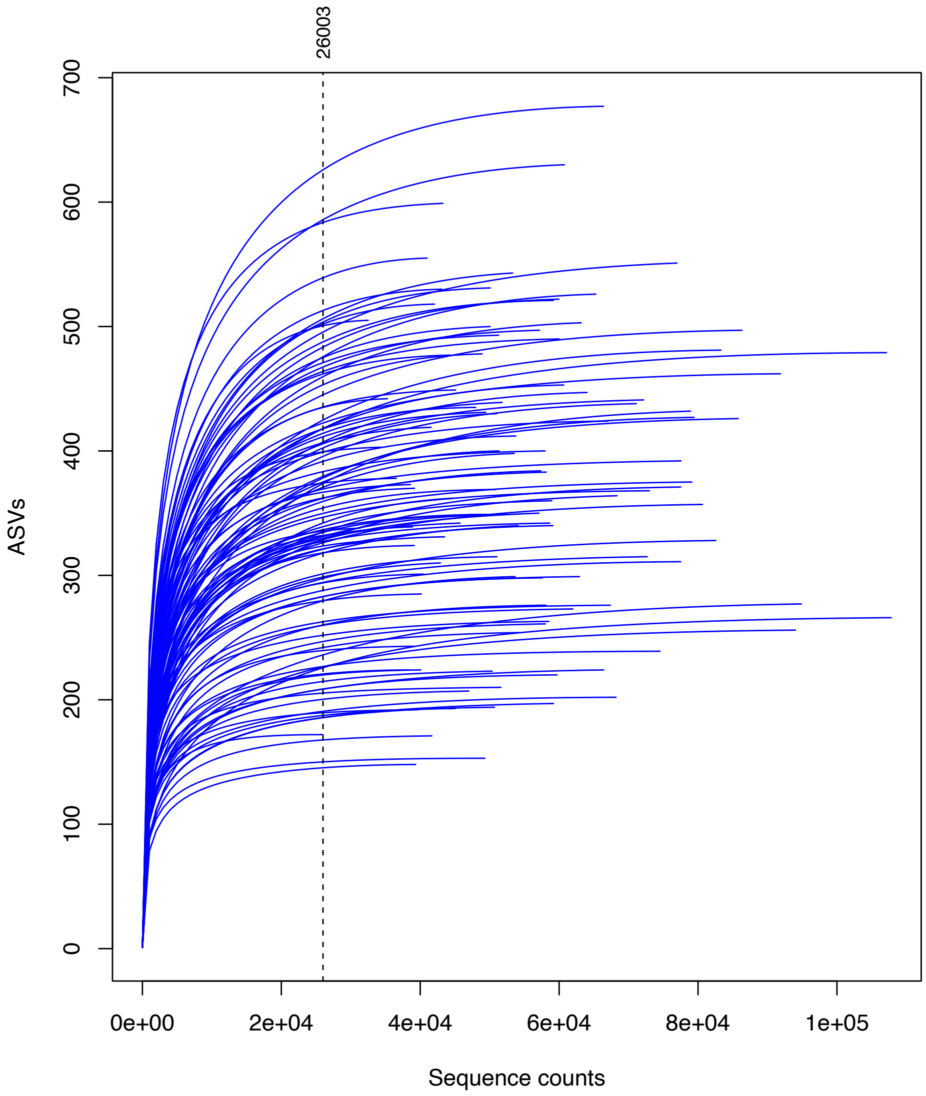


**Figure S1:** Rarefaction curves (step= 1000 counts) summarizing sequencing effort per sample/specimen (109). The dashed line shows the sample with minimum sequence coverage (260003 sequence counts).
