## Supporting Information, Figure S2 for "Insular holobionts: persistence and seasonal plasticity of the Balearic wall lizard (*Podarcis lilfordi*) gut microbiota"

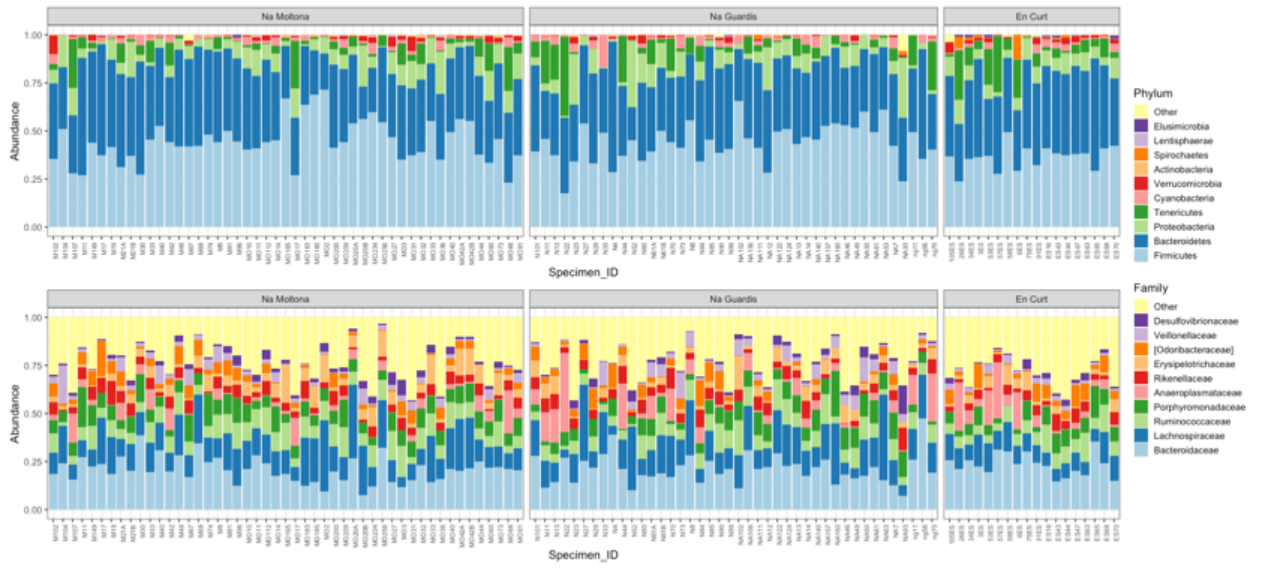


**Figure S2:** Microbiota taxonomic composition (phylum and family) at specimen level. Legends list only the top ten most abundant taxa. The remaining were included in "Others".
