## Supporting Information, Figure S3 for "Insular holobionts: persistence and seasonal plasticity of the Balearic wall lizard (*Podarcis lilfordi*) gut microbiota"

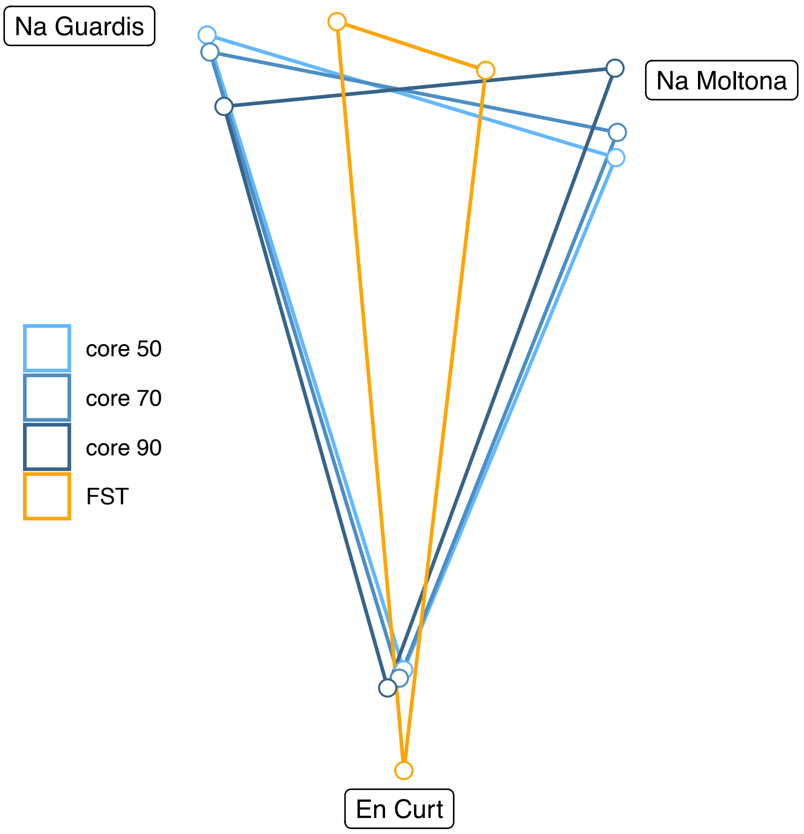


**Figure S3:** Microbiota centroid distances according to unweighted Unifrac, estimated on different core subsets, and host genetic distances according to *Fst* values estimated from microsatellites data (Rotger et al. 2021).
