## Supporting Information, Figure S4 for "Insular holobionts: persistence and seasonal plasticity of the Balearic wall lizard (*Podarcis lilfordi*) gut microbiota"

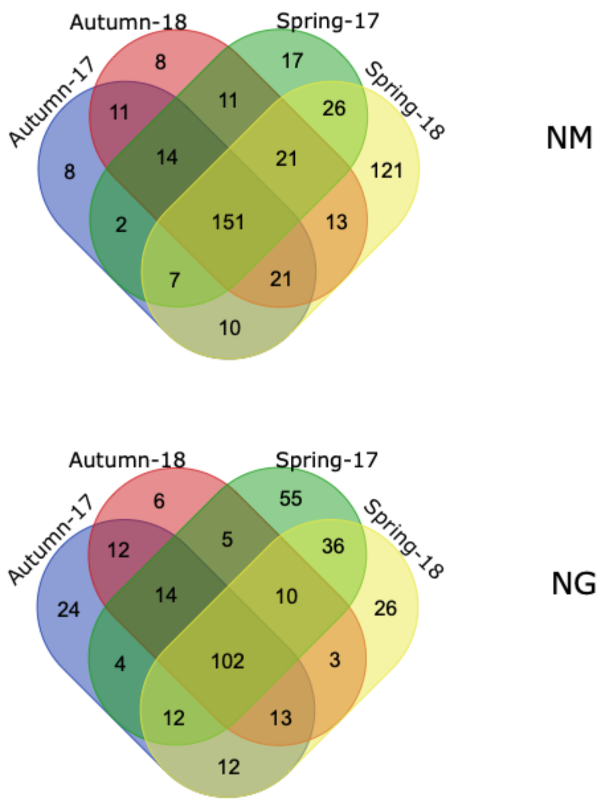


**Figure S4:** Venn diagrams showing shared ASVs across the four sampling dates. For each date, we considered only ASVs found in at least 50% of the specimens.
