## Supporting Information, Figure S5 for "Insular holobionts: persistence and seasonal plasticity of the Balearic wall lizard (*Podarcis lilfordi*) gut microbiota"

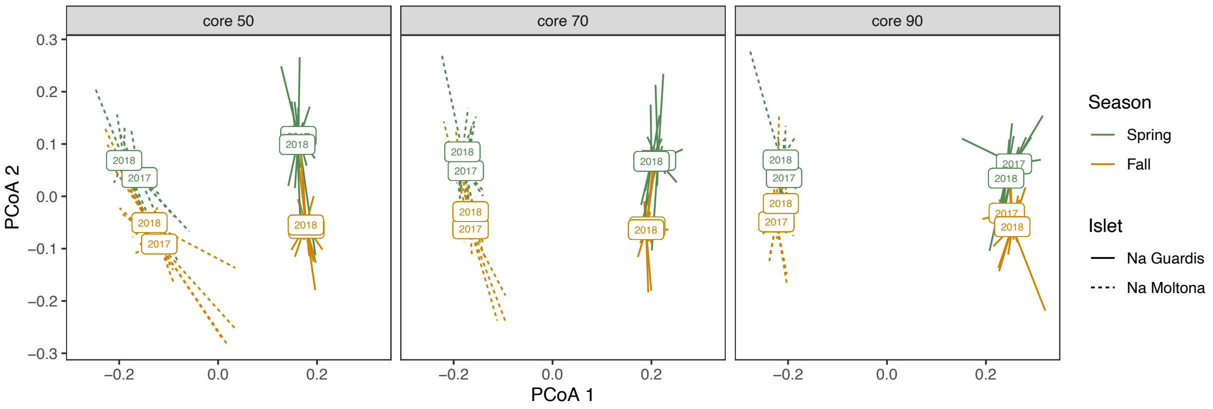


**Figure S5:** PCoA of microbiota Bray-Curtis distances according to Date, estimated on different core subsets (50, 70, 80 and 90%). The rectangular box represents the centroid per date.
