## Supporting Information, Figure S6 for "Insular holobionts: persistence and seasonal plasticity of the Balearic wall lizard (*Podarcis lilfordi*) gut microbiota"

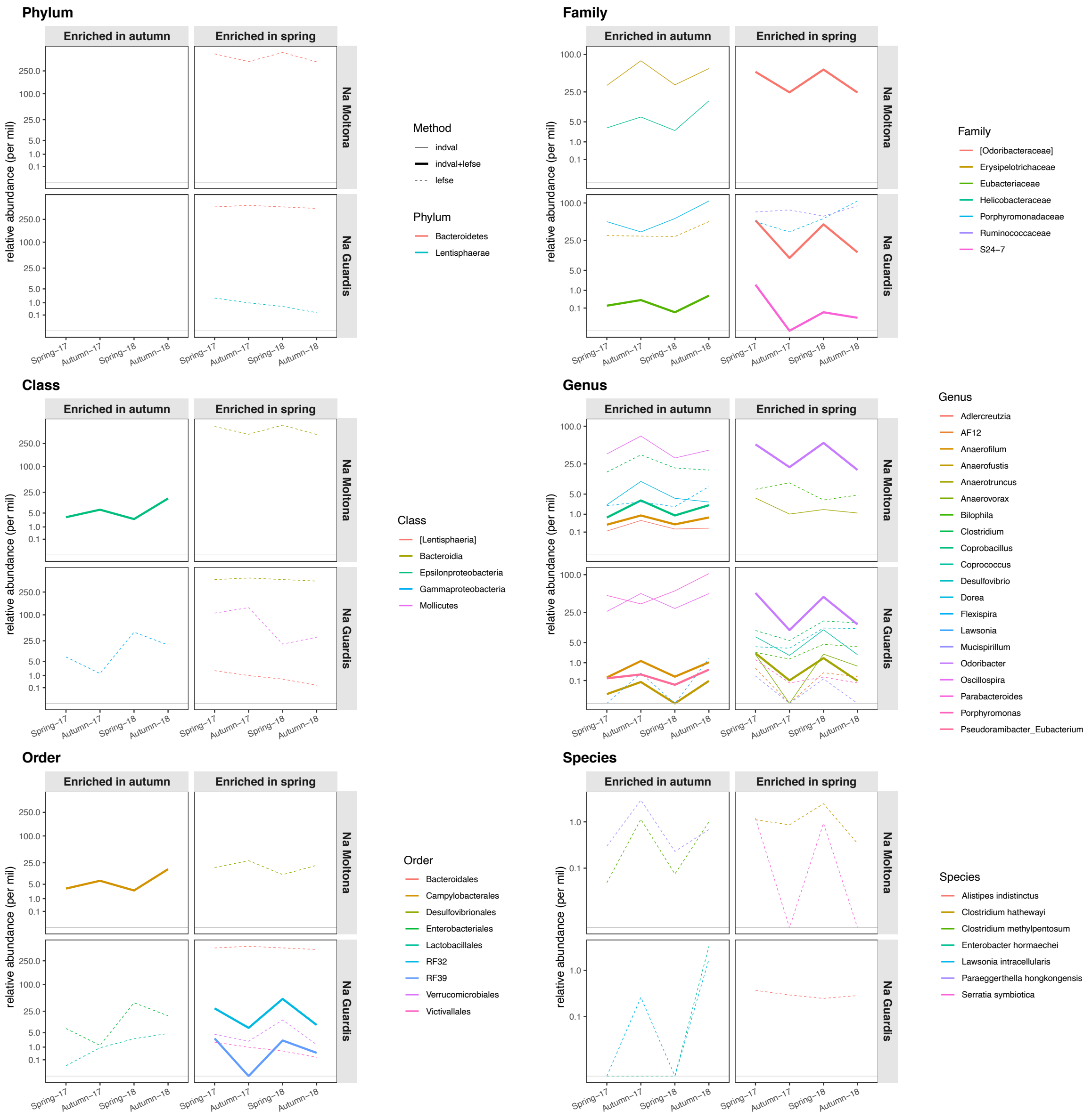


**Figure S6:** Variation in relative abundance along "dates" of taxa that were significantly enriched in either spring or autumn according to LEfSe and/or indval analyses.
